# AlphaFold prediction and analysis of Adhesion-family G protein coupled receptor (aGPCR) structures

**DOI:** 10.1101/2024.12.17.628075

**Authors:** Xiao Wen (Kelly) Zhou, Eddie Chen, Rithwik Ramachandran

## Abstract

Adhesion GPCRs (aGPCRs), a family of 33 receptors, regulate crucial physiological processes through a unique self-activation mechanism. We employed AlphaFold, a state-of-the-art AI system, to predict 3D structures of aGPCRs in active and inactive conformations. Full-length models were obtained from the AlphaFold Protein Structure Database, while tethered-ligand (TL) exposed structures were predicted using AlphaFold. Comparison with available experimentally solved structures were made to validate accuracy of Alphafold aGPCR model. For receptors without experimentally solved structures, we compare structures of inactive (TL masked) and active (TL exposed) aGPCRs. In the active structures, the tethered-ligand was docked in the transmembrane bundle for several receptors but surprisingly not all receptors. This AI driven analysis of aGPCR structures provides a framework for exploring alternative receptor activation mechanisms. These models also offer valuable insights into receptor conformational changes and TL interactions, paving the way for the development of novel pharmacological tools and accelerating drug discovery efforts targeting this important receptor family.

---

G protein-coupled receptors (GPCRs) are the largest class of cell membrane receptors and are regulators of diverse physiological and pathological processes. GPCRs are also valuable and highly tractable targets for the development of therapeutic drugs^1^. Adhesion GPCRs (aGPCRs) are the second largest family of GPCRs (Family B2, 33 members) and are emerging as important regulators of many physiological and pathophysiological processes^2–4^. Despite this growing recognition of aGPCRs regulated cellular events, there are limited pharmacological tools for studying these receptors and currently no approved therapies that directly target them. This can be largely attributed to the unusual activation mechanism in these receptors that involves mechanical force mediated separation of the receptor into two fragments to initiate signaling.

aGPCRs are made up of a large extracellular N-terminal fragment (NTF) and a seven-transmembrane spanning domain C-terminal fragment (CTF) that signals through intracellular effector proteins^1^. aGPCR NTFs have diverse structural features that consist of motifs, often repeating, that allow interaction with cellular or matrix components^5^. Apart from ADGRA1, all aGPCRs also have a conserved GPCR autoproteolysis-inducing (GAIN) domain that encapsulates a hydrophobic region of the C-terminal Fragment (CTF) referred to as the statchel sequence or tethered-ligand (TL) (Figure 1). During biosynthesis and trafficking of aGPCRs, the TL region is released at the GPCR proteolysis site (GPS) in an autocatalytic manner through nucleophilic attack on the Leucine-Threonine peptide bond by the Threonine R-group^6^. In inactive receptors at the cell surface, the TL remains encapsulated in the NTF through weak non-covalent forces.

**Figure 1.**
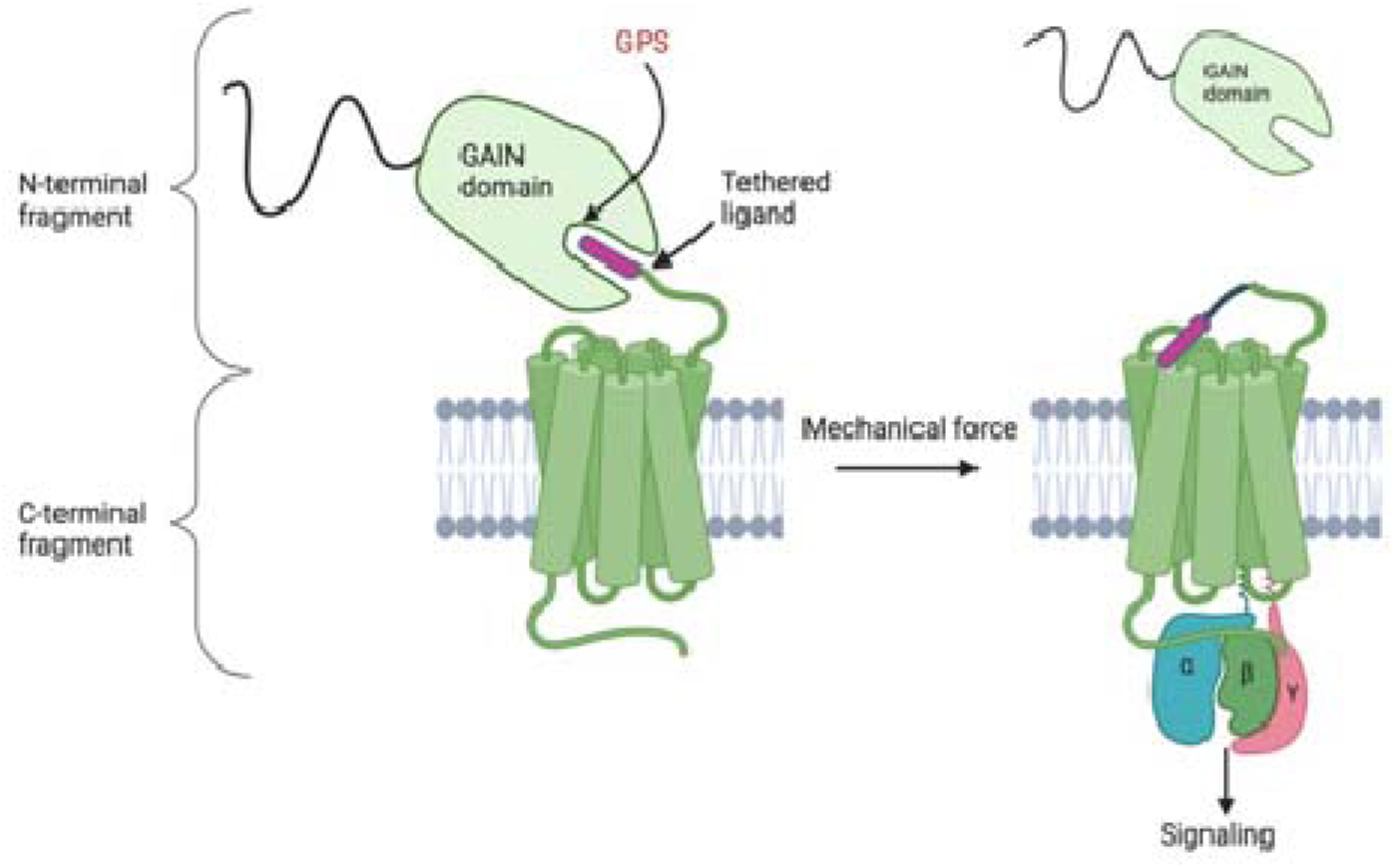
General Structures of Adhesion Family G protein-coupled receptors (aGPCRs). The N-terminal fragment (NTF) and C-terminal fragment (CTF) in aGPCRs are non-covalently bonded in the inactive receptor. Cell-cell or cell-matrix interactions and mechanical forces dissociate the NTF from the membrane embedded CTF at the GPCR proteolysis site (GPS) within the autoproteolysis-inducing (GAIN) domain to activate the receptor. NTF and CTF separation reveals a sequence of amino acids referred to as the tethered-ligand (TL) and is thought to act as a self-activator. Members of the aGPCRs family have varied NTF domains that can interact with various ligands, though the specific binding properties and precise activation mechanism are still unclear.

It is now well established that aGPCR activation requires NTF interaction with an extracellular binding partner and mechanical force mediated separation of the NTF and CTF fragments at the GPS site^5^. Whether the unmasked TL is important for activation of aGPCRs is still debated, however, there is now evidence in some aGPCRs to show that the unmasked TL docks into the transmembrane bundle of the aGPCR CTF, in what may be considered as the orthosteric ligand binding pocket, to initiate receptor mediated signaling. Conversely, studies have demonstrated that aGPCRs with the entire TL deleted retain the ability to activate signaling responses^7^, and that GPS cleavage is not essential for receptor function^8^, which support a TL independent mode of activation that involves the receptor adopting an active signaling competent confirmation upon removal of the NTF.

Given the unusual mechanical force mediated activation, studying aGPCR signaling and the development of pharmacological tools to study receptor function remains a challenging task. Mechanical dissociation of aGPCR fragments is not readily compatible with traditional ligand screening assays and has limited progress. Recent advances in computational screening of ligands could provide a solution but these approaches are restricted by the limited number of solved aGPCR protein structures. The hydrophobic and dynamic nature of aGPCRs and the limited knowledge of ligand-binding partners to form stable receptor-ligand complexes for have presented a challenge for achieving high and accurate resolution of aGPCR structures, however, recent advances in Cryo-electron microscopy (Cyro-EM) have enabled structure elucidation for several aGPCRs^9–12^. In addition to traditional experimental approaches for determining protein structures, recent advances in AI systems for predicting protein structures enables a novel avenue for aGPCR protein structure determination.

AlphaFold is an artificial intelligence neural network-based model that was developed by DeepMind to predict the three-dimensional structure of proteins based on the amino acid sequences. AlphaFold was first introduced in the 2020 Critical Assessment of protein Structure Prediction (CASP14) demonstrating accuracy comparable to experimental methods and outperforming other computational methods^9^. The latest version of AlphaFold incorporates physical and biological knowledge of protein structures along with multiple-sequence alignments into a deep learning algorithm to predict protein structures. AlphaFold works by analyzing the amino acid of proteins and comparing it to a large database of proteins that share homology and evolutionary relationships. It then performs a multiple sequence alignment (MSA) by aligning the target protein with homologous sequences. The MSA provides information about patterns of amino acid conservation and variation across different species and predicts residue-residue contacts within a protein^13^. For every predicted structure, AlphaFold produces a per-residue model confidence score (pLDDT) between 0-100. This score measures how well the predicted structure would agree with an experimental structure based on the local distance test Cα (lDDT- Cα)^10^. This study will use ColabFold, an implementation of AlphaFold, that provides faster prediction of protein structures and complexes using homology searches with MMseqs2 in AlphaFold2^11^.

Since most aGPCR structures have not been experimentally solved, we used AlphaFold to predict aGPCR structures in NTF separated TL exposed “active” state and compared it to the full-length structure where the TL is encapsulated within the GAIN domain as the “inactive” model. We hypothesized that AlphaFold would predict aGPCR structures and reveal structural rearrangements that occur when aGPCRs are activated. We further sought to compare the outputs from AlphaFold to the experimentally solved aGPCR structures that are available in the protein databank (PDB).

## RESULTS

### AlphaFold Predictions Align with Solved Structures

As an initial test of the accuracy of AlphaFold prediction of aGPCR structures, we compared predicted TL exposed structural models of ADGRD1, ADGRG1, ADGRG2, ADGRG3, ADGRG4, ADGRG5, ADGRF1, ADGRE5, and ADGRL3 to experimentally solved structures for these receptors. In all cases, we find that the AlphaFold predicted models showed high structural similarity to the previously published solved structures (Figure 2). The predicted and solved structures were superimposed and the Root Mean Squared Deviation (RMSD) values were measured to quantify the similarity between them by comparing the distance between atoms. An RSMD value less than 2.0 Å typically indicates high structural similarity^12^. For all seven receptors assessed here, the RMSD values for the TL exposed receptors were around 1.0 Å (Supplementary Table 1), indicating that the AlphaFold predictions reliably predicted the structures of these receptors. The predicted TL exposed receptors to the inactive receptor where the TL is encapsulated in the NTF gain domain were also superimposed and compared (Figure 5). Surprisingly, the predicted models of both TL exposed active and FL inactive models also showed a high degree of structural similarity. Regions with the highest RMSD shifts between the TL exposed and FL models are the intracellular and extracellular loops, TM6, and as expected, the TL region (Figure 5). The TL regions of ADGRG2 and ADGRG3 are absent in the solved structures, therefore these structures could not be assessed in this manner. In support of the accuracy of AlphaFold Predictions, the structure of recently solved ADGRE5 (CD97)^13^ showed structural similarity to the predicted FL and TL exposed models that were generated prior to the publication and addition of these models to the PDB (Figure 2). In comparison to the solved ADGRE5 structure, the most prominent RMSD shifts were shown in TM6 and TM5 (Figure 2) for both the predicted TL exposed and FL structures.

**Figure 2.**
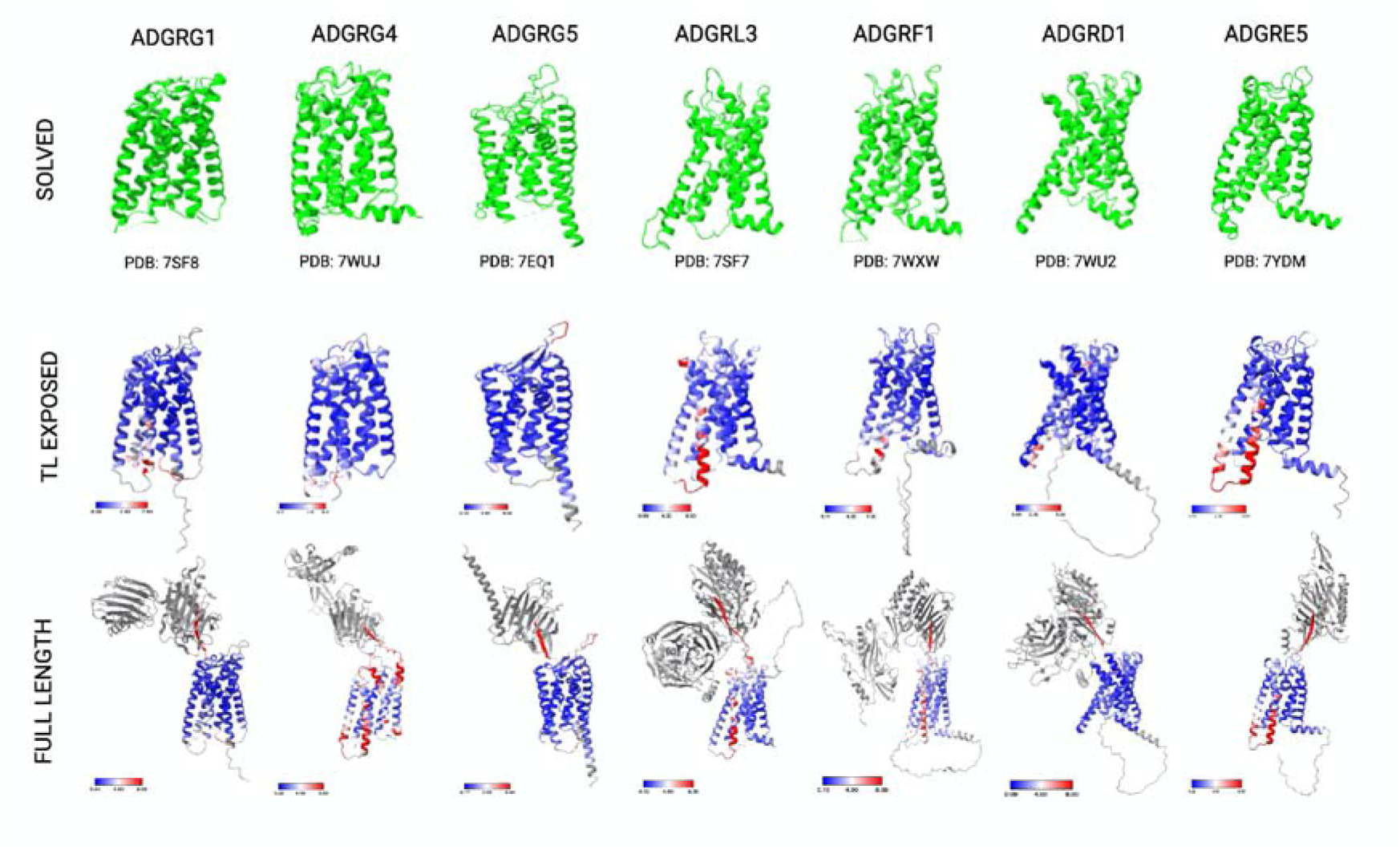
RMSD Shifts Between Solved aGPCR structures and AlphaFold Predicted Models. Structures represented in green are solved structures (left), tethered-ligand exposed predictions overlayed with solved structures are represented in the middle, and full-length structures overlayed with solved structures are represented on the right. Root Mean Squared Deviation (RMSD) values were calculated for each figure. RMSD colour bar is shown for all overlayed figures with blue areas representing areas with a smaller RMSD shifts, and red areas represent higher RMSD compared to the solved structure. Here, we see that the predicted tethered-ligand exposed model and the full-length structures adopt similar conformations compared to the solved structure. The RMSD shift for all predicted structures were less than 2.0 Å, suggesting similarity to solved model. As expected, the TL region in the full-length structure has the highest RMSD value compared to the solved structure. This suggests that AlphaFold was able to predict structural models that are similar to known solved models.

### Structural Changes Shown in Active and Inactive aGPCRs

The RMSD shifts show that the full-length predicted structures have more regions with higher RMSD shifts compared to the solved structures (Figure 2). This could be indicative of possible structural changes associated with the transition from full-length inactive to TL exposed active structures. However, one caveat is that AlphaFold pLDDT confidence scores varied between different models and may contribute to the perceived structural difference. For example, the AlphaFold predicted ADGRG4 structure (Supplementary Figure S-1) has several regions with low confidence scores which may contribute to why the RMSD shifts of the predicted full-length structure differs from the solved structures. Conversely, the predicted structure of ADGRG3 has a fairly high pLDDT score indicative of a high structural confidence (Supplementary Figure S-1) and still exhibits high RMSD shifts indicative of the structural changes accompanying receptor activation. When assessing RMSD shifts between solved and predicted structures, it is therefore important to consider prediction confidence scores to assess whether high RMSD shifts are a reflection of structural changes or a consequence of low model confidence.

The TL in the full-length predictions are all encapsulated within the GAIN domain and in the TL exposed model, the TL is docked in the TMD binding pocket for the majority of aGPCRs (22 out of 33). When the GAIN domain is mechanically separated from the TL, AlphaFold shows that the most favourable structure for many aGPCRs to adopt is to have the hydrophobic TL bind back into the TMD orthosteric pocket. This supports the notion that aGPCRs may be TL activated, though exceptions to this conformation were also seen for several aGPCRs (Figure 4). In the case of ADGRL1-2, ADGRE4, ADGRA1-3, ADGRC1-3, ADGRB1-3, and ADGRV1, we see that the exposed tethered ligand is positioned outside the TMD and in most cases adopts a helical conformation. For ADGRA1, there is no apparent TL sequence and the NTF will be highlighted as the TL. This observation is significant and suggests that multiple mechanisms may exist for aGPCR activation following NTF and CTF separation.

Finally, we also see that the TMD regions of the TL exposed active state and FL inactive states do not significantly differ with the exception of TM6. This is in keeping with shifts seen in class B1 receptors where the most pronounced change between active and inactive conformations are found in the outward swing of TM6^14^. For example, Glucagon-like peptide 1 receptor (GLP-1R) shows a pronounced TM6 shift in the active form compared to the inactive form, however the other TMD regions do not show extensive movement (Supplementary Figure S-2).

### Tethered-Ligand Binding Pocket of Solved Structures Align with Predicted Models

As discussed, following activation of aGPCRs, the TL undergoes a significant reorientation and is proposed to bind an orthosteric pocket in the transmembrane bundle. The predicted AlphaFold models of active ADGRD1, ADGRG2, ADGRG3, ADGRG4, ADGRG5, ADGRF1, ADGRL3, and ADGRE5 show the TL docked into the orthosteric pocket of the TMD. This is comparable to what was observed in the solved structures for these receptors which were also obtained in the active TL exposed conformation. The overlay of the solved TL binding pocket with the predicted TL binding pocket showed structural similarities as well as comparable alignments in the TL binding position and interactions with the TMD (Figure 3). In all cases where the exposed TL predicted active model docks the TL into the orthosteric binding site, the TL forms a partial helical conformation similar to that reported in the solved structures. The solved structures for ADGRL3 and ADGRD1 showed unstructured TLs, however, AlphaFold predicted a structured helical TL conformation. Here we see that the TL docks and interacts with all the transmembrane domains with little to no interactions with TM4 (Figure 3). TL binding contacts were determined for the remaining 27 unsolved aGPCRs (Figure 4). The unsolved aGPCRs that have the TL docked into the orthosteric pocket showed similar TL binding contacts with the solved models where the TL interacts with all the transmembrane domains with no interactions with TM4.

**Figure 3.**
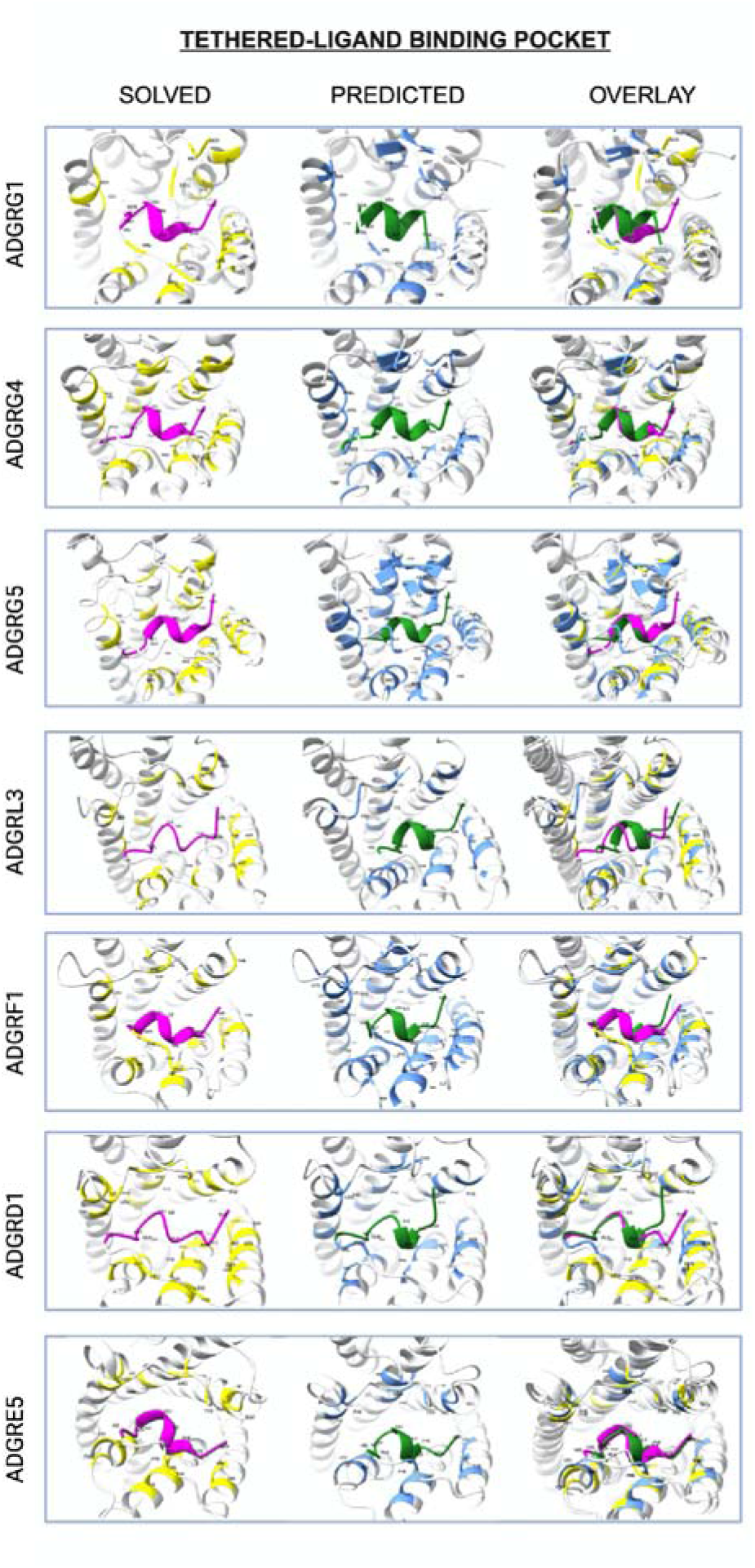
Comparison of Tethered-Ligand Binding Pocket for Solved aGPCR structures and AlphaFold Predicted Models. Tethered-Ligand (TL) binding contacts were computed using ChimeraX to identify tethered-ligand interactions with the transmembrane bundle that are less than 4 Å apart. Solved structures are in active forms where the tethered ligand is docked into an orthosteric site on the receptor. Predicted AlphaFold structure of TL exposed receptors follow similar conformation where the TL is docked into a binding pocket within the transmembrane domain. TL contacts of the solved structures matched with predicted models. Predicted models often show additional contacts that solved structures did not show.

**Figure 4.**
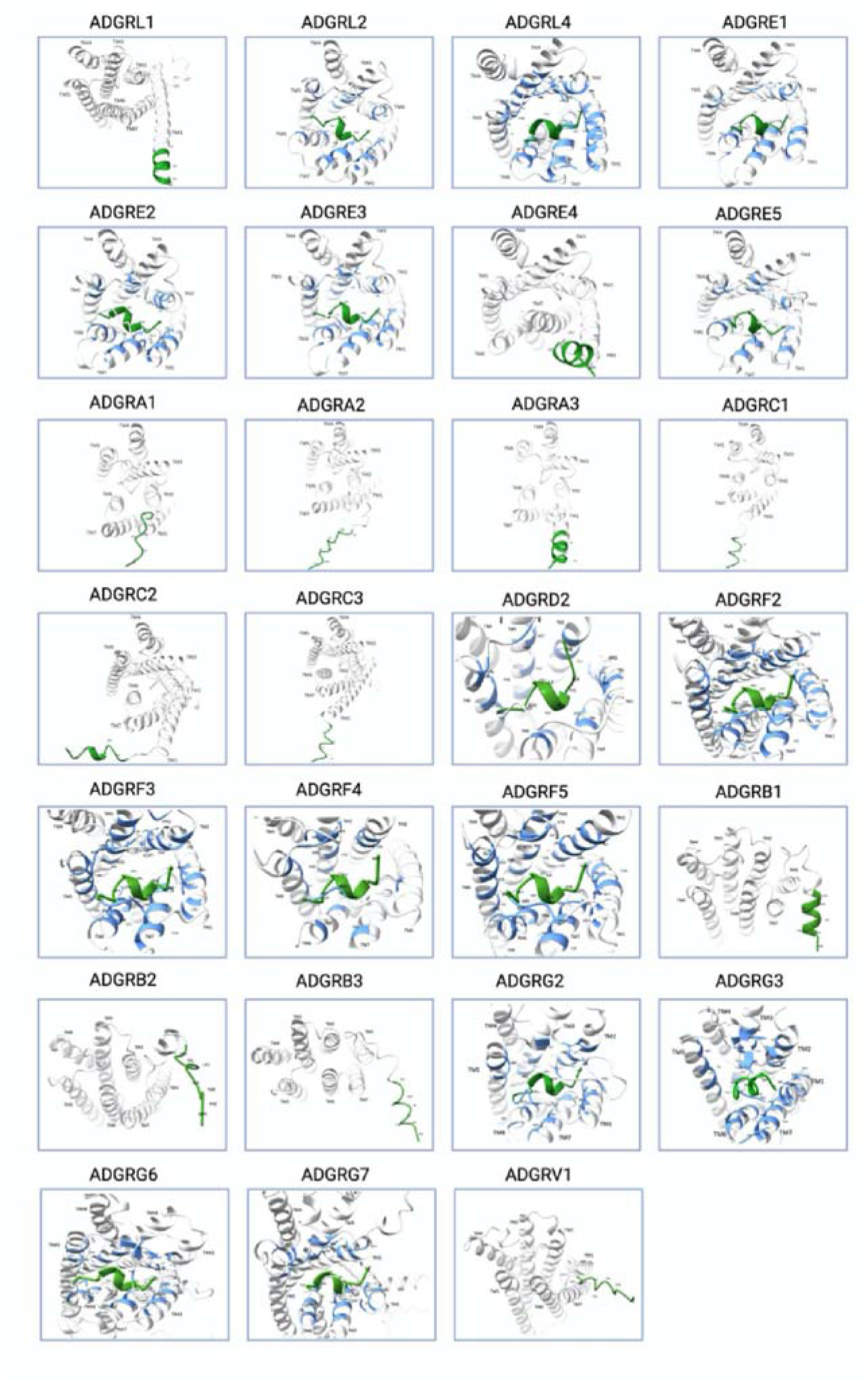
AlphaFold Prediction of Tethered-Ligand Binding Pocket for Unsolved aGPCR Structures. TL binding contacts were computed using ChimeraX to identify tethered-ligand interactions with the transmembrane bundle that are less than 4 Å apart. The tethered ligand is coloured in green, and the TL binding contacts are coloured in blue. The unsolved predicted TL binding contacts are similar to the solved predicted TL binding contacts. Most of the structures have the TL docked into the orthosteric pocket, however, the TL lies outside the transmembrane domain for some aGPCRs which include: ADGRB1-3, ADGRV1, ADGRC1-3, ADGRA1-3, ADGRE4-5, ADGRL1. This could suggest that exposing the TL may not initiate TL docking into the orthosteric pocket in these aGPCRs.

As mentioned previously, several aGPCRs have their TL positioned outside the pocket, including ADGRL1-2, ADGRE4, ADGRA1-3, ADGRC1-3, ADGRB1-3, and ADGRV1 (Figure 4). This could be a result of the low pLDDT confidence for some of these AlphaFold predicted structures (Supplementary Figure S-1) resulting in low incorrect positioning of the exposed TL. For example, ADGRV1, ADGRB2, and ADGRB3 have unstructured TLs, however, the TL for ADGRB1 and ADGRL1 shows the expected helical TL conformation, but the TL is still positioned outside the TMD bundle when exposed. This could indicate that the exposed TL may not favour docking into the TMD binding pocket.

## DISCUSSION

Current studies that elucidated aGPCR structures suggest a TL dependent mechanism of activation. Computational and functional studies also propose TL dependent activation for different aGPCRs^15,16^. Cryo-EM active-state structures of GPR56 and show that the decrypted TL bends downward into the core of the 7TM adopting a partial alpha-helical fold and stabilizing an active conformation through interacting with several conserved regions with TM1, TM2, TM6, TM7 and extracellular loop 2 (ECL2)^11^. Findings of TL dependent activity were further supported in functional studies which found that mutations to the YFLM residues on the TL of GPR56 significantly reduced receptor activity as measured through GTPase turnover assays^11^. Similar active-state conformation were seen for several aGPCRs including ADGRL3 (LPHN3), ADGRD1, ADGRE5, and ADGRF1^13,16,17^. These studies have led to the current consensus that TL exposure initiates aGPCR activation by orthosteric binding. Interestingly however, the AlphaFold predictions for several aGPCRs do not exhibit TL docking into the orthosteric pocket when exposed. This pattern was often seen in several aGPCRs with large N-terminal fragments. For example, one of the larger aGPCRs, ADGRB1 (BAI1) has the TL positioned outside of the TMD in the active model. Interestingly previous studies with engineered BAI1 mutants that lacked the entire NTF (including the TL) showed signaling activity which provides support for an alternative model where conformational change triggered by the removal of the NTF rather than the TL docking enables receptor activation and signalling^7^. Several other studies have also shown TL independent signaling of aGPCRs including one which showed that monobodies that targeted various NTF regions were still able to initiate signaling activity in mutants that were deficient in autoproteolysis and TL exposure^18^. A recent study presented differences in tethered agonist signalling in aGPCRs where they showed that almost half of human aGPCRs lack TL dependent activity as shown in functional profiling assaying including transcriptional report responses (CRE, SRE, SRF-RE, NFAT-RE) and G protein interaction analyses^19^. This study also ran AlphaFold predictions that align with the current study’s data on TL docking of TL exposed models. Both studies did not show TL binding into the orthosteric pocket for ADGRL1, ADGRA1-3, ADGRB1-3, ADGRC1-3, and ADGRV1 in the structural AlphaFold analysis^19^.

This suggests that even though all aGPCRs (except ADGRA1) have a conserved TL region, the exposed TL and it’s interaction with the TMD ligand binding pocket may not be necessary for the activation of aGPCRs. The interactions between the NTF and the 7TM domain for aGPCRs is also not well characterized and the large NTF of these aGPCRs may be involved in allosteric communication to activate or inhibit signaling from these receptors.

Such TL independent pattern of activation in aGPCRs could be reminiscent of mechanisms seen in the Class B1 or secretin-like GPCR family which are GPCRs that are the most closely evolutionarily related to Class B2 aGPCRs. Class B1 GPCRs are activated by peptide hormone agonists that first bind to the N-terminus of the receptor, causing a conformational change that directs the extracellular loops to open up at the extracellular face of the receptor, allowing the peptide N-terminus to the dock into the TMD orthosteric ligand binding site to initiate signaling (Figure S2)^14^. Overall, the AlphaFold predictions showing that TL docking is not seen in all aGPCRs support the need to explore different models of aGPCR activation in order to consider different strategies for aGPCR targeted drug development.

The majority of aGPCRs do show the TL docked into the orthosteric binding pocket consistent with the current predominant model and in these cases, show similar interacting residues within the TMD (Figure 5). The solved structures obtained from Barros-Alvarez et al., (2022) found that TMD residues in TM1, TM2, ECL2, and TM6 were areas of TL binding for GPR56 (C411^1.47^, F454^2.64^, W577^45.51^, W617^6.53^, and I620^6.56^) and LPHN3 (I872^1.47^, F914^2.64^, F938^3.4^, W1000^45.51^, W1068^6.53^ and, L1072^6.57^). This matches what we see in both the predicted and solved structures (Figure 3). More TL binding contacts were also evident in both the predicted and solved structures, which may be indicative of additional TL TMD interaction sites (Figure S-3). Solved structures for ADGRE5 (CD97) found that the TL interacted with four highly conserved aromatic residues across solved aGPCR structures on the 7TM domain: F597^2.64^, W685^ECL^, W753^6.53^, and F771^7.42^. These interacting residues were also highlighted in the predicted CD97 structure (Figure 2), was predicted prior to the publication of the solved structure. Identifying interactions of the TL to the TMD binding domains is an obvious avenue for drug development as several studies have found that TL mimicking peptides can initiate signalling^20^. One caveat to the development of drug therapies that mimic the TL and target the orthosteric pocket is that the TL region is highly conserved across all aGPCRs thus making target specificity a challenge. Experiments showed that the TL mimicking peptide can activate more than one member of the aGPCR subfamily^21^, therefore, AlphaFold predicted models provides a tool for identifying TL interacting residues across the aGPCR family develop drugs that are more specific to the receptor of interest. Since we saw previously that some aGPCRs may not exhibit TL docking when exposed, drugs can also be developed to target the NTF or allosteric sites in the TMD to overcome the challenge of identifying specific ligands.

**Figure 5.**
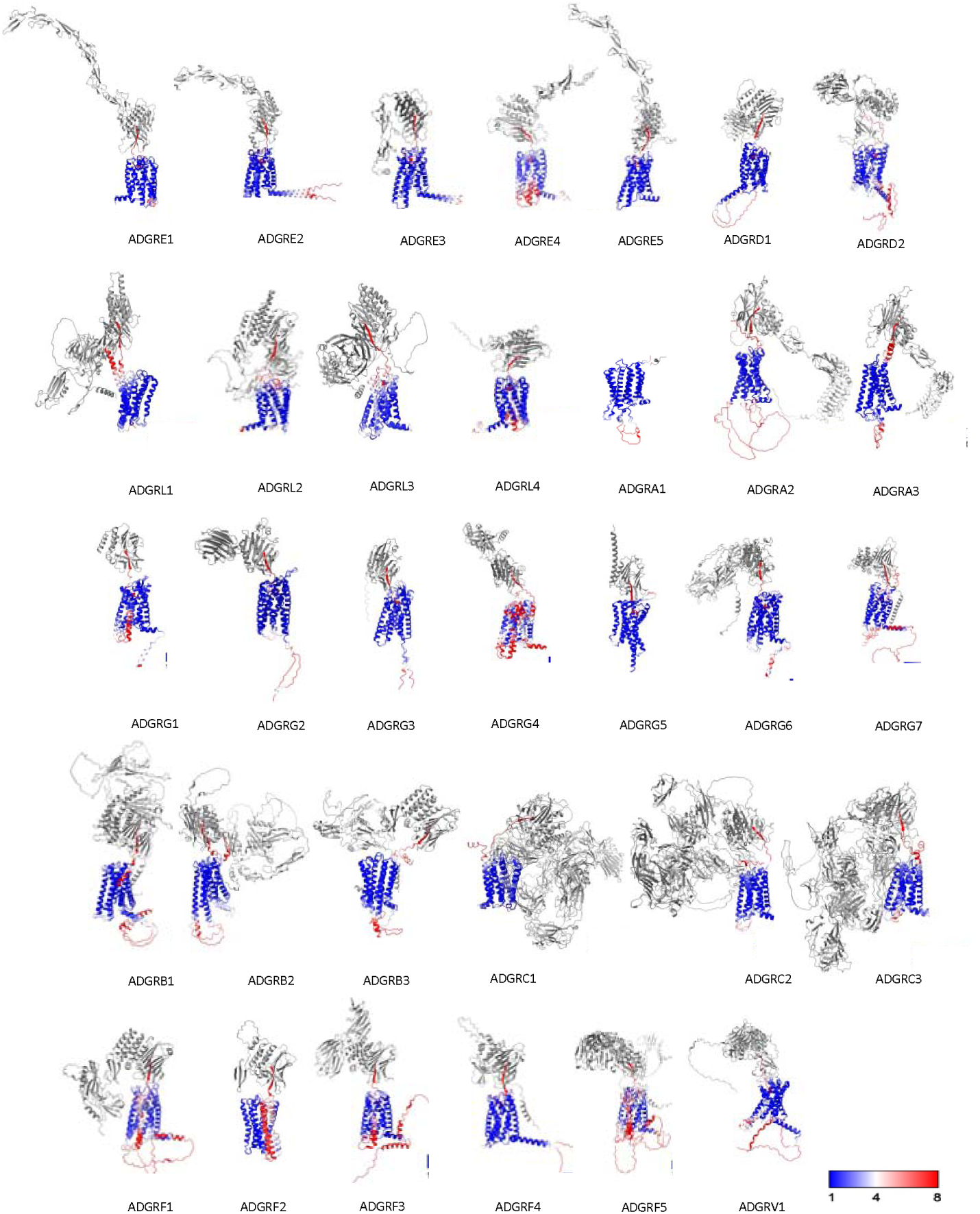
Overlayed Structures of Tethered-Ligand Exposed and Full-Length aGPCR Structures. Predictions showed high confidence for the transmembrane core and had lower structural confidence for extracellular and intracellular domains. Root Mean Square Deviation (RMSD) colour bar shown for the overlayed image to represent the RMSD shift between TL exposed and FL prediction. Blue regions represent smaller RMSD shifts, red regions represent larger shifts, and grey areas represent regions that have RMSD values greater than 8. Regions that have higher RMSD shifts are the TL region, and the TMD regions and TM6 and TM7. AlphaFold predictions show that full-length structure and TL exposed structure adopt similar conformations.

Overall AlphaFold predictions appear to reliably predict aGPCR structures and show good agreement to solved structure with values less than 2.0 Å. One concern of using AlphaFold is whether it reproduces or replicate previously experimentally determined structures as the predicted structure. Since the predicted structures were not identical to solved structures, this is however unlikely. AlphaFold generated structures also accurately predicted the interactions of the TL to the 7TM domain within the receptor for several receptors. Interestingly, the predicted CD97 structure that was generated prior to the publication of the solved structure exhibited very similar structural properties as well further supporting the accuracy and utility of AlphaFold predictions. The solved structure of CD97 was however in complex with an engineered G_q_ protein which may account for the shifts seen in TM5 and TM6. This suggests that AlphaFold predictions of aGPCRs with TL exposed or not may not reflect their fully active states and additional prediction and analysis of the structures in complex with different effectors is warranted.

AlphaFold appears to be a valuable tool for predicting aGPCR structures and can provide valuable insights into this complex receptor systems. While our current study, along with many others, has shown the accuracy of AlphaFold protein predictions, it is important to acknowledge the significance of low confidence pLDDT scores in the predictions. Low pLDDT scores indicate regions where predicted structures are less reliable and are less likely to be correctly positioned. However, low pLDDT scores may also correlate with protein structures that are highly mobile and intrinsically unstructured^22^. These areas are a challenge for experimental structure determination by cryo-EM and crystallography and are very often stabilized through additional protein engineering approached. AlphaFold provides a starting point to analyzing these dynamic regions though further efforts are needed to fully comprehend their contribution to receptor function. aGPCRs initiate various signaling pathways upon activation through the recruitment of various G proteins. A study done with ADGRL3 in complex with 4 major G proteins revealed coupling to G_q_, G_i_, G_s_, and G ^23^. Similar computational modelling can be done using AlphaFold to predict structural characteristics of aGPCRs and assess effector docking in the CTF. The potential for virtual docking and drug screening with AlphaFold is also promising. Since most aGPCRs are still unsolved, leveraging AlphaFold to uncover common binding residues in the TMD presents an avenue for functional studies and pharmacological tool development for aGPCRs.

## Conclusion

The comparison of solved aGPCR protein structures to the AlphaFold predicted structures displayed several key conclusions. The exposed TL predominately docks in orthosteric binding pocket for several aGPCRs supporting the notion of TL meditated activation. The identification of undocked TL structures suggests additional determinants of aGPCR activation. Overall AlphaFold modelling accurately predicted aGPCR structures and uncovered TL binding contacts allowing the integration of additional molecular docking and experimental methodologies to advance aGPCR targeted ligand discovery.

### Experimental procedures

#### aGPCR selection of solved structures

Experimentally determined aGPCR structures were chosen from the list of cryo-EM solved structures recorded by Seufer et al., (2023). Only human aGPCRs were chosen and those with the tethered agonist/statchel included the solved structure. The predicted AlphaFold structures were compared with the following solved aGPCR structures from the protein databank: ADGRG1 (PDB: 7SF8), ADGRD1 (PDB: 7UW2), ADGRG2 (PDB: 7XKE), ADGRG3 (PDB: 7D77), ADGRG4 (PDB: 7WUJ), ADGRG5 (PDB: 7EQ1), ADGRF1 (PDB: 7WXW), ADGRL3 (PDB: 7SF7), and ADGRE5 (7YDM). Additional proteins bound to solved structure were removed using ChimeraX hide tool for better depiction of receptor structures. These structures include G protein complexes as well as bound ligands (See Supplementary Data). Regions with very low confidence and unstructured regions were also hidden (See Supplementary Data).

#### Selection of Full-Length and Tethered Ligand Exposed Sequences

Full protein sequences for all 33 aGPCRs were obtained from UniProt. In each case, the top-ranked reviewed human isoform 1 of each receptor was selected, and its entry identifier was noted. TL exposed models were obtained by predicting structures of protein sequences starting from the first TL residue as defined by Xiao, P. et al. (2022) down to the end of the CTF.

#### Generation of Full-Length and Tethered Ligand Exposed Structures

Colabfold v.1.5.3 implementation of AlphaFold2 installed on ChimeraX v.1.6.1 was used to make structural predictions of all aGPCR human proteins. To generate homology models of each of the aGPCRs in both inactive and active signaling conformations, full-length and TL- exposed sequences respectively were submitted to the ColabFold software server, which predicted the structures. In each case, ColabFold generated 5 models based on the ‘alphafold2_ptm’ modeling mode using an ‘unpaired_paired’ pair mode. Each model was run for 3 recycles with default ColabFold parameters: without PDB templates and without amber relaxation/energy minimization. Multiple sequence alignments were generated using MMSeqs2. Each protein’s models were ranked based on their Predicted Template Modeling Score (pTM) and the top-ranked model as determined by ColabFold for each inactive and active signaling state aGPCR was designated as ‘best_model’ and selected for further analysis. A per-residue estimate of confidence (pLDDT) score was computed for each protein prediction. This measure corresponds to the models’ predicted score on the IDDT- Cα metric. Regions with high confidence for model accuracy are coloured in dark blue (pLDDT >90), regions that are moderately confident are yellow (pLDDT =70-90), and regions with low confidence are coloured in yellow (pLDDT < 50).

### Structural Comparisons

#### RMSD Shift

To calculate differences in solved and predicted structures, Root Mean Squared Deviation (RMSD) values were calculated. ChimeraX alignment tool was used to align solved and predicted structures and the RMSD values were also calculated using ChimeraX. Predicted TL exposed model and predicted FL models were also aligned and RMSD values were calculated as above using ChimeraX. The overlay of structures was coloured using RMSD colour bar with areas of lower RMSD shifts represented in blue, and areas with higher RMSD shifts represented in red.

#### Tethered-Ligand Binding Contacts

Solved structures and predicted structures were aligned using ChimeraX. TL region was recoloured in pink for solved structures and green for predicted TL exposed structures. The binding contacts were calculated using ChimeraX “contacts” tool and coloured in yellow for solved structures and blue for predicted structures. Contacts were filtered using center-center distance less than or equal to 4 Å. Interactions between atoms 4 or fewer bonds apart and interactions between residues <5 apart in sequences were ignored. Solved structures were compared to identify common TL binding contacts within the TMD. Contacts were considered as common TMD binding contacts if they were present in 4 or more of the solved structures.

## Data availability

All data are available in the manuscript or appended supplementary files. All AlphaFold predictions and associated information is provided in the supplementary data folders.

## Supporting information

Full-Length Predictions

TL Exposed Predictions

Figure S-1

Figure S-2

Figure S-3

Table S-1

Hidden Sequences AlphaFold

## Author contributions

XWZ, EC and RR designed the experiments. EC ran ColabFold predictions. XWZ and EC analysed predicted structures. XWZ, EC and RR wrote the manuscript.

## Funding and additional information

These studied were funded by a discovery grant from the Natural Sciences and Engineering Research Council of Canada (NSERC). EC was the recipient of a Schulich school of Medicine and Dentistry Dean’s undergraduate research opportunity (DUROP) scholarship.

## Conflict of interest

The authors declare no conflict of interest.

## Notes

### Competing Interest Statement

The authors have declared no competing interest.

