## Supplementary material for "AlphaFold prediction and analysis of Adhesion-family G protein coupled receptor (aGPCR) structures": AlphaFold Supplementary Data.docx

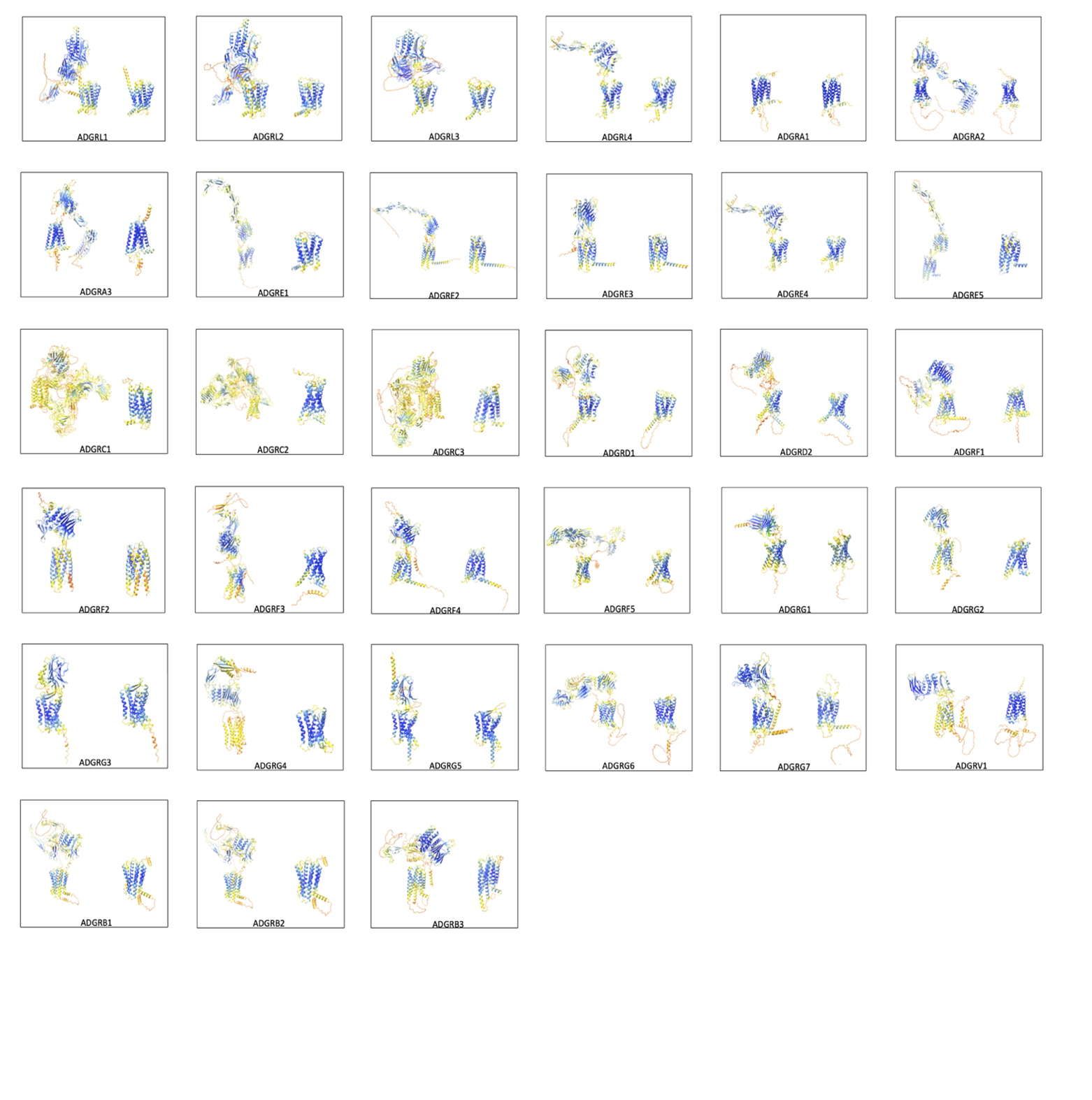

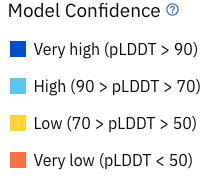

***Figure S-1. Tethered-Ligand Exposed and Full Length Structures of Adhesion Family G Protein-Coupled Receptors.*** The active and inactive aGPCRs predicted models were compared. In the active structures, the sequence downstream of the first TL residue was entered and predicted in AlphaFold. The entire aGPCR sequence was retrieved for the inactive/full-length structure. Most of the aGPCRs have high pLDDT scores, with the transmembrane domain having the highest structural confidence. Full-length structures that have the lowest structural confidence is ADCRC1-3. ADGRG4, and ADGRV1. In all cases, the TL is docked within the GAIN domain of receptors in the inactive form. When the TL is exposed in the inactive form, most aGPCRs will have the TL dock within the orthosteric pocket of the transmembrane domain, however some aGPCRs do not adopt this conformation.

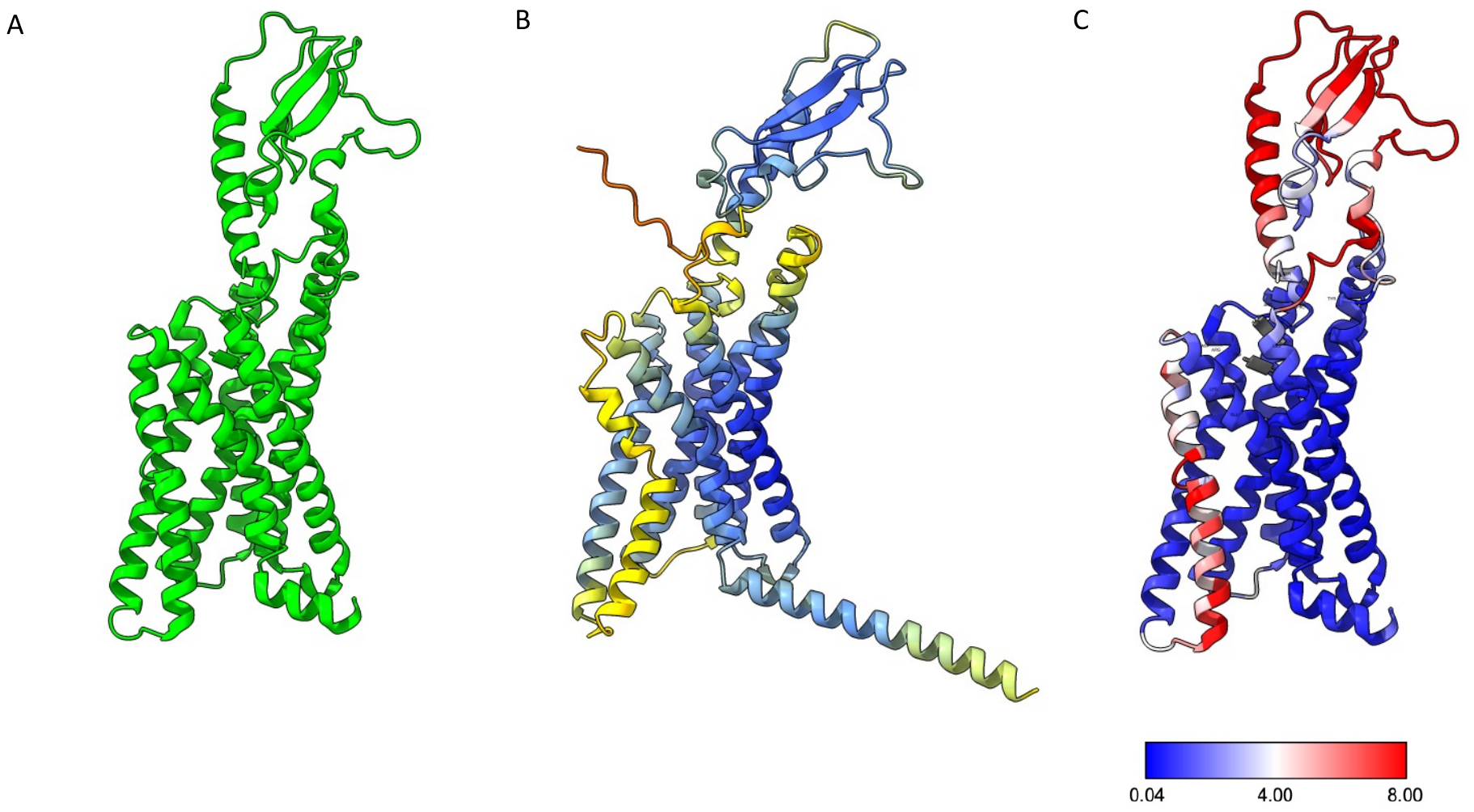

***Figure S-2. Comparison of Solved and Predicted Structure of Glucagon-like peptide 1 receptor***

***(GLP-1R).*** Solved structure of Glucagon-like peptide 1 receptor (GLP-1R) (A) (PDB: 5NX2) shown in green was compared to AlphaFold predicted model (B). The predicted model was colored using pLDDT scores where regions with higher structural confidence (pLDDT score greater than 90) is coloured in blue and regions with lower confidence (pLDDT score less than 70) were coloured in yellow and orange. Solved and predicted model were compared (C) and regions with higher Root Mean Square Deviation (R.M.S.D) shifts are shown in red and regions with lower R.M.S.D shifts are shown in blue. The highest R.M.S.D values were shown in the NTF region and transmembrane domain 6. The difference may be due to that fact that the predicted model is not in complex with a peptide agonist and may represent an inactive form.

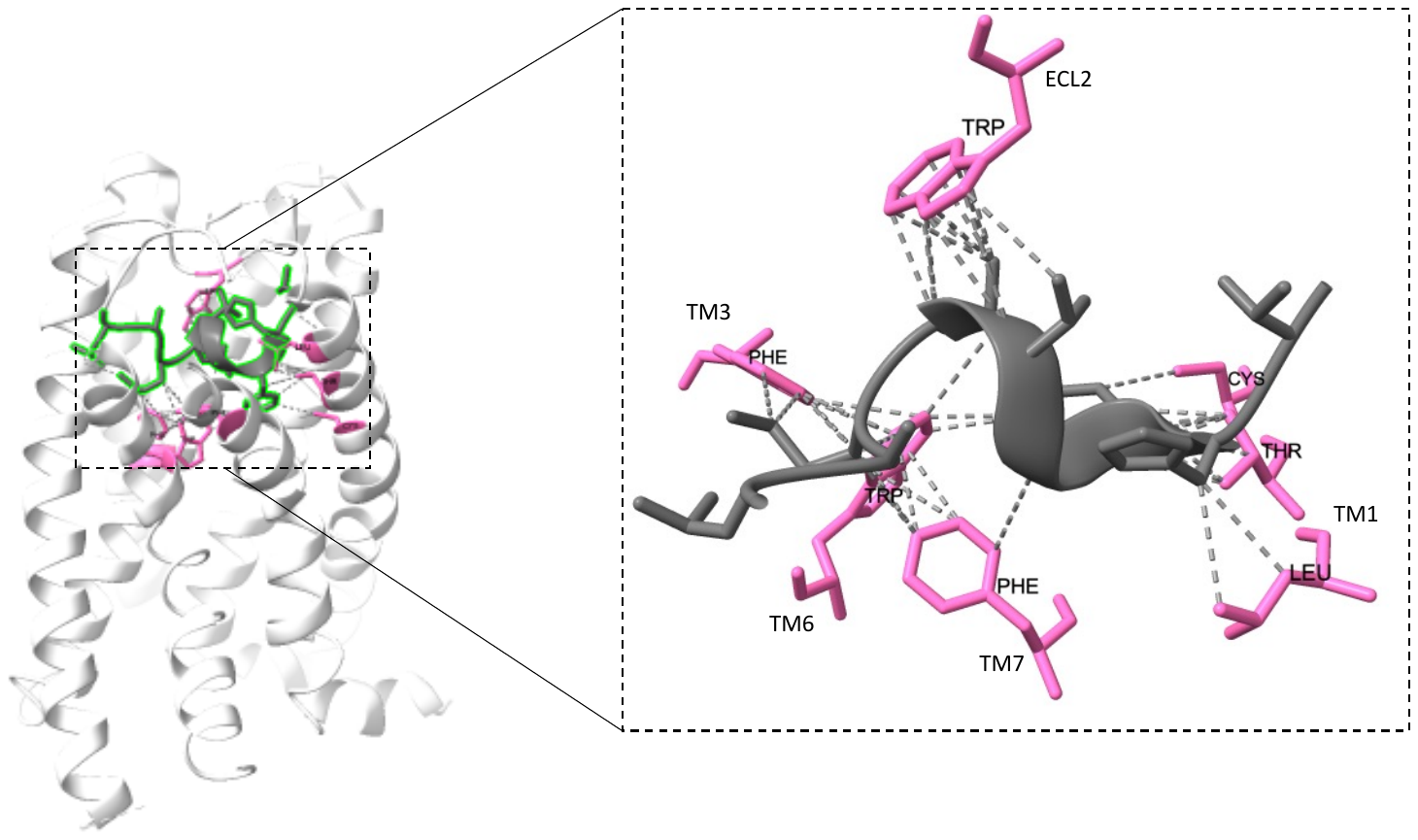

***Figure S-3. Common binding contacts of solved aGPCR structures.*** TL binding contacts were computed using ChimeraX to identify tethered-ligand interactions with the transmembrane bundle that are less than 4 Å apart. Tethered ligand is coloured in grey and the binding contacts were coloured in pink. Comparison of binding contacts for solved structures showed common TMD residues in TM1 (CYS, THR, LEU), ECL2 (TRP), TM3 (PHE), TM6 (TRP), and TM7 (PHE).

***Table S-1. Root-Mean Squared Deviation Values between Solved and Predicted Structures***. Calculated values of Root-Mean Squared Deviation (RMSD) values of solved aGPCR structures to the predict tethered-ligand exposed or full-length model. All RMSD values were less than the value of 2.0 Å, indicating high structural similarity.

| aGPCR | RMSD of Solved vs Tethered-Ligand Exposed Model (Å) | RMSD value of Solved vs Full-Length Model (Å) |
| --- | --- | --- |
| ADGRD1 | 0.875 | 0.753 |
| ADGRG1 | 0.968 | 0.885 |
| ADGRG4 | 0.908 | 1.083 |
| ADGRG5 | 0.949 | 0.919 |
| ADGRF1 | 1.018 | 1.085 |
| ADGRL3 | 1.117 | 1.057 |
| ADGRE5 | 1.118 | 1.058 |

***Table S-2. Amino Acid Sequences of Full- length and Tethered-Ligand Exposed aGPCRs***

**aGPCR Amino Acid Sequences: putative tethered-ligand is highlighted.**

**Sequences highlighted in red are unfolded/disordered regions hidden in figures**

**>sp|Q86SQ6|AGRA1_HUMAN Adhesion G protein-coupled receptor A1 OS=Homo sapiens OX=9606 GN=ADGRA1 PE=2 SV=3**

MDLKTVLSLPRYPGEFLHPVVYACTAVMLLCLLASFVTYIVHQSAIRISRKGRHTLLNFC

FHAALTFTVFAGGINRTKYPILCQAVGIVLHYSTLSTMLWIGVTARNIYKQVTKKAPLCL

DTDQPPYPRQPLLRFYLVSGGVPFIICGVTAATNIRNYGTEDEDTAYCWMAWEPSLGAFY

GPAAIITLVTCVYFLGTYVQLRRHPGRRYELRTQPEEQRRLATPEGGRGIRPGTPPAHDA

PGASVLQNEHSFQAQLRAAAFTLFLFTATWAFGALAVSQGHFLDMVFSCLYGAFCVTLGL

FVLIHHCAKREDVWQCWWACCPPRKDAHPALDANGAALGRAACLHSPGLGQPRGFAHPPG

PCKMTNLQAAQGHASCLSPATPCCAKMHCEPLTADEAHVHLQEEGAFGHDPHLHGCLQGR

TKPPYFSRHPAEEPEYAYHIPSSLDGSPRSSRTDSPPSSLDGPAGTHTLACCTQGDPFPM

VTQPEGSDGSPALYSCPTQPGREAALGPGHLEMLRRTQSLPFGGPSQNGLPKGKLLEGLP

FGTDGTGNIRTGPWKNETTV

**AF structure NTF is misfolded. No apparent TL sequence**.

**>sp|Q86SQ6|AGRA1_HUMAN Adhesion G protein-coupled receptor A1 OS=Homo sapiens OX=9606 GN=ADGRA1 PE=2 SV=3 – TL EXPOSED**

SLPRYPGEFLHPVVYACTAVMLLCLLASFVTYIVHQSAIRISRKGRHTLLNFC

FHAALTFTVFAGGINRTKYPILCQAVGIVLHYSTLSTMLWIGVTARNIYKQVTKKAPLCL

DTDQPPYPRQPLLRFYLVSGGVPFIICGVTAATNIRNYGTEDEDTAYCWMAWEPSLGAFY

GPAAIITLVTCVYFLGTYVQLRRHPGRRYELRTQPEEQRRLATPEGGRGIRPGTPPAHDA

PGASVLQNEHSFQAQLRAAAFTLFLFTATWAFGALAVSQGHFLDMVFSCLYGAFCVTLGL

FVLIHHCAKREDVWQCWWACCPPRKDAHPALDANGAALGRAACLHSPGLGQPRGFAHPPG

PCKMTNLQAAQGHASCLSPATPCCAKMHCEPLTADEAHVHLQEEGAFGHDPHLHGCLQGR

TKPPYFSRHPAEEPEYAYHIPSSLDGSPRSSRTDSPPSSLDGPAGTHTLACCTQGDPFPM

VTQPEGSDGSPALYSCPTQPGREAALGPGHLEMLRRTQSLPFGGPSQNGLPKGKLLEGLP

FGTDGTGNIRTGPWKNETTV

**AF structure NTF is misfolded. No apparent TL sequence**.

**>sp|Q96PE1|AGRA2_HUMAN Adhesion G protein-coupled receptor A2 OS=Homo sapiens OX=9606 GN=ADGRA2 PE=1 SV=2**

MGAGGRRMRGAPARLLLPLLPWLLLLLAPEARGAPGCPLSIRSCKCSGERPKGLSGGVPG

PARRRVVCSGGDLPEPPEPGLLPNGTVTLLLSNNKITGLRNGSFLGLSLLEKLDLRNNII

STVQPGAFLGLGELKRLDLSNNRIGCLTSETFQGLPRLLRLNISGNIFSSLQPGVFDELP

ALKVVDLGTEFLTCDCHLRWLLPWAQNRSLQLSEHTLCAYPSALHAQALGSLQEAQLCCE

GALELHTHHLIPSLRQVVFQGDRLPFQCSASYLGNDTRIRWYHNRAPVEGDEQAGILLAE

SLIHDCTFITSELTLSHIGVWASGEWECTVSMAQGNASKKVEIVVLETSASYCPAERVAN

NRGDFRWPRTLAGITAYQSCLQYPFTSVPLGGGAPGTRASRRCDRAGRWEPGDYSHCLYT

NDITRVLYTFVLMPINASNALTLAHQLRVYTAEAASFSDMMDVVYVAQMIQKFLGYVDQI

KELVEVMVDMASNLMLVDEHLLWLAQREDKACSRIVGALERIGGAALSPHAQHISVNARN

VALEAYLIKPHSYVGLTCTAFQRREGGVPGTRPGSPGQNPPPEPEPPADQQLRFRCTTGR

PNVSLSSFHIKNSVALASIQLPPSLFSSLPAALAPPVPPDCTLQLLVFRNGRLFHSHSNT

SRPGAAGPGKRRGVATPVIFAGTSGCGVGNLTEPVAVSLRHWAEGAEPVAAWWSQEGPGE

AGGWTSEGCQLRSSQPNVSALHCQHLGNVAVLMELSAFPREVGGAGAGLHPVVYPCTALL

LLCLFATIITYILNHSSIRVSRKGWHMLLNLCFHIAMTSAVFAGGITLTNYQMVCQAVGI

TLHYSSLSTLLWMGVKARVLHKELTWRAPPPQEGDPALPTPSPMLRFYLIAGGIPLIICG

ITAAVNIHNYRDHSPYCWLVWRPSLGAFYIPVALILLITWIYFLCAGLRLRGPLAQNPKA

GNSRASLEAGEELRGSTRLRGSGPLLSDSGSLLATGSARVGTPGPPEDGDSLYSPGVQLG

ALVTTHFLYLAMWACGALAVSQRWLPRVVCSCLYGVAASALGLFVFTHHCARRRDVRASW

RACCPPASPAAPHAPPRALPAAAEDGSPVFGEGPPSLKSSPSGSSGHPLALGPCKLTNLQ

LAQSQVCEAGAAAGGEGEPEPAGTRGNLAHRHPNNVHHGRRAHKSRAKGHRAGEACGKNR

LKALRGGAAGALELLSSESGSLHNSPTDSYLGSSRNSPGAGLQLEGEPMLTPSEGSDTSA

APLSEAGRAGQRRSASRDSLKGGGALEKESHRRSYPLNAASLNGAPKGGKYDDVTLMGAE

VASGGCMKTGLWKSETTV

**>sp|Q96PE1|AGRA2_HUMAN Adhesion G protein-coupled receptor A2 OS=Homo sapiens OX=9606 GN=ADGRA2 PE=1 SV=2** **– TL EXPOSED**

GNVAVLMELSAFPREVGGAGAGLHPVVYPCTALL

LLCLFATIITYILNHSSIRVSRKGWHMLLNLCFHIAMTSAVFAGGITLTNYQMVCQAVGI

TLHYSSLSTLLWMGVKARVLHKELTWRAPPPQEGDPALPTPSPMLRFYLIAGGIPLIICG

ITAAVNIHNYRDHSPYCWLVWRPSLGAFYIPVALILLITWIYFLCAGLRLRGPLAQNPKA

GNSRASLEAGEELRGSTRLRGSGPLLSDSGSLLATGSARVGTPGPPEDGDSLYSPGVQLG

ALVTTHFLYLAMWACGALAVSQRWLPRVVCSCLYGVAASALGLFVFTHHCARRRDVRASW

RACCPPASPAAPHAPPRALPAAAEDGSPVFGEGPPSLKSSPSGSSGHPLALGPCKLTNLQ

LAQSQVCEAGAAAGGEGEPEPAGTRGNLAHRHPNNVHHGRRAHKSRAKGHRAGEACGKNR

LKALRGGAAGALELLSSESGSLHNSPTDSYLGSSRNSPGAGLQLEGEPMLTPSEGSDTSA

APLSEAGRAGQRRSASRDSLKGGGALEKESHRRSYPLNAASLNGAPKGGKYDDVTLMGAE

VASGGCMKTGLWKSETTV

**>sp|Q8IWK6|AGRA3_HUMAN Adhesion G protein-coupled receptor A3 OS=Homo sapiens OX=9606 GN=ADGRA3 PE=1 SV=2**

MEPPGRRRGRAQPPLLLPLSLLALLALLGGGGGGGAAALPAGCKHDGRPRGAGRAAGAAE

GKVVCSSLELAQVLPPDTLPNRTVTLILSNNKISELKNGSFSGLSLLERLDLRNNLISSI

DPGAFWGLSSLKRLDLTNNRIGCLNADIFRGLTNLVRLNLSGNLFSSLSQGTFDYLASLR

SLEFQTEYLLCDCNILWMHRWVKEKNITVRDTRCVYPKSLQAQPVTGVKQELLTCDPPLE

LPSFYMTPSHRQVVFEGDSLPFQCMASYIDQDMQVLWYQDGRIVETDESQGIFVEKNMIH

NCSLIASALTISNIQAGSTGNWGCHVQTKRGNNTRTVDIVVLESSAQYCPPERVVNNKGD

FRWPRTLAGITAYLQCTRNTHGSGIYPGNPQDERKAWRRCDRGGFWADDDYSRCQYANDV

TRVLYMFNQMPLNLTNAVATARQLLAYTVEAANFSDKMDVIFVAEMIEKFGRFTKEEKSK

ELGDVMVDIASNIMLADERVLWLAQREAKACSRIVQCLQRIATYRLAGGAHVYSTYSPNI

ALEAYVIKSTGFTGMTCTVFQKVAASDRTGLSDYGRRDPEGNLDKQLSFKCNVSNTFSSL

ALKNTIVEASIQLPPSLFSPKQKRELRPTDDSLYKLQLIAFRNGKLFPATGNSTNLADDG

KRRTVVTPVILTKIDGVNVDTHHIPVNVTLRRIAHGADAVAARWDFDLLNGQGGWKSDGC

HILYSDENITTIQCYSLSNYAVLMDLTGSELYTQAASLLHPVVYTTAIILLLCLLAVIVS

YIYHHSLIRISLKSWHMLVNLCFHIFLTCVVFVGGITQTRNASICQAVGIILHYSTLATV

LWVGVTARNIYKQVTKKAKRCQDPDEPPPPPRPMLRFYLIGGGIPIIVCGITAAANIKNY

GSRPNAPYCWMAWEPSLGAFYGPASFITFVNCMYFLSIFIQLKRHPERKYELKEPTEEQQ

RLAANENGEINHQDSMSLSLISTSALENEHTFHSQLLGASLTLLLYVALWMFGALAVSLY

YPLDLVFSFVFGATSLSFSAFFVVHHCVNREDVRLAWIMTCCPGRSSYSVQVNVQPPNSN

GTNGEAPKCPNSSAESSCTNKSASSFKNSSQGCKLTNLQAAAAQCHANSLPLNSTPQLDN

SLTEHSMDNDIKMHVAPLEVQFRTNVHSSRHHKNRSKGHRASRLTVLREYAYDVPTSVEG

SVQNGLPKSRLGNNEGHSRSRRAYLAYRERQYNPPQQDSSDACSTLPKSSRNFEKPVSTT

SKKDALRKPAVVELENQQKSYGLNLAIQNGPIKSNGQEGPLLGTDSTGNVRTGLWKHETT

V

**>sp|Q8IWK6|AGRA3_HUMAN Adhesion G protein-coupled receptor A3 OS=Homo sapiens OX=9606 GN=ADGRA3 PE=1 SV=2 – TL EXPOSED**

SNYAVLMDLTGSELYTQAASLLHPVVYTTAIILLLCLLAVIVS

YIYHHSLIRISLKSWHMLVNLCFHIFLTCVVFVGGITQTRNASICQAVGIILHYSTLATV

LWVGVTARNIYKQVTKKAKRCQDPDEPPPPPRPMLRFYLIGGGIPIIVCGITAAANIKNY

GSRPNAPYCWMAWEPSLGAFYGPASFITFVNCMYFLSIFIQLKRHPERKYELKEPTEEQQ

RLAANENGEINHQDSMSLSLISTSALENEHTFHSQLLGASLTLLLYVALWMFGALAVSLY

YPLDLVFSFVFGATSLSFSAFFVVHHCVNREDVRLAWIMTCCPGRSSYSVQVNVQPPNSN

GTNGEAPKCPNSSAESSCTNKSASSFKNSSQGCKLTNLQAAAAQCHANSLPLNSTPQLDN

SLTEHSMDNDIKMHVAPLEVQFRTNVHSSRHHKNRSKGHRASRLTVLREYAYDVPTSVEG

SVQNGLPKSRLGNNEGHSRSRRAYLAYRERQYNPPQQDSSDACSTLPKSSRNFEKPVSTT

SKKDALRKPAVVELENQQKSYGLNLAIQNGPIKSNGQEGPLLGTDSTGNVRTGLWKHETT

V

**>sp|O14514|AGRB1_HUMAN Adhesion G protein-coupled receptor B1 OS=Homo sapiens OX=9606 GN=ADGRB1 PE=1 SV=2**

MRGQAAAPGPVWILAPLLLLLLLLGRRARAAAGADAGPGPEPCATLVQGKFFGYFSAAAV

FPANASRCSWTLRNPDPRRYTLYMKVAKAPVPCSGPGRVRTYQFDSFLESTRTYLGVESF

DEVLRLCDPSAPLAFLQASKQFLQMRRQQPPQHDGLRPRAGPPGPTDDFSVEYLVVGNRN

PSRAACQMLCRWLDACLAGSRSSHPCGIMQTPCACLGGEAGGPAAGPLAPRGDVCLRDAV

AGGPENCLTSLTQDRGGHGATGGWKLWSLWGECTRDCGGGLQTRTRTCLPAPGVEGGGCE

GVLEEGRQCNREACGPAGRTSSRSQSLRSTDARRREELGDELQQFGFPAPQTGDPAAEEW

SPWSVCSSTCGEGWQTRTRFCVSSSYSTQCSGPLREQRLCNNSAVCPVHGAWDEWSPWSL

CSSTCGRGFRDRTRTCRPPQFGGNPCEGPEKQTKFCNIALCPGRAVDGNWNEWSSWSACS

ASCSQGRQQRTRECNGPSYGGAECQGHWVETRDCFLQQCPVDGKWQAWASWGSCSVTCGA

GSQRRERVCSGPFFGGAACQGPQDEYRQCGTQRCPEPHEICDEDNFGAVIWKETPAGEVA

AVRCPRNATGLILRRCELDEEGIAYWEPPTYIRCVSIDYRNIQMMTREHLAKAQRGLPGE

GVSEVIQTLVEISQDGTSYSGDLLSTIDVLRNMTEIFRRAYYSPTPGDVQNFVQILSNLL

AEENRDKWEEAQLAGPNAKELFRLVEDFVDVIGFRMKDLRDAYQVTDNLVLSIHKLPASG

ATDISFPMKGWRATGDWAKVPEDRVTVSKSVFSTGLTEADEASVFVVGTVLYRNLGSFLA

LQRNTTVLNSKVISVTVKPPPRSLRTPLEIEFAHMYNGTTNQTCILWDETDVPSSSAPPQ

LGPWSWRGCRTVPLDALRTRCLCDRLSTFAILAQLSADANMEKATLPSVTLIVGCGVSSL

TLLMLVIIYVSVWRYIRSERSVILINFCLSIISSNALILIGQTQTRNKVVCTLVAAFLHF

FFLSSFCWVLTEAWQSYMAVTGHLRNRLIRKRFLCLGWGLPALVVAISVGFTKAKGYSTM

NYCWLSLEGGLLYAFVGPAAAVVLVNMVIGILVFNKLVSKDGITDKKLKERAGASLWSSC

VVLPLLALTWMSAVLAVTDRRSALFQILFAVFDSLEGFVIVMVHCILRREVQDAVKCRVV

DRQEEGNGDSGGSFQNGHAQLMTDFEKDVDLACRSVLNKDIAACRTATITGTLKRPSLPE

EEKLKLAHAKGPPTNFNSLPANVSKLHLHGSPRYPGGPLPDFPNHSLTLKRDKAPKSSFV

GDGDIFKKLDSELSRAQEKALDTSYVILPTATATLRPKPKEEPKYSIHIDQMPQTRLIHL

STAPEASLPARSPPSRQPPSGGPPEAPPAQPPPPPPPPPPPPQQPLPPPPNLEPAPPSLG

DPGEPAAHPGPSTGPSTKNENVATLSVSSLERRKSRYAELDFEKIMHTRKRHQDMFQDLN

RKLQHAAEKDKEVLGPDSKPEKQQTPNKRPWESLRKAHGTPTWVKKELEPLQPSPLELRS

VEWERSGATIPLVGQDIIDLQTEV

**>sp|O14514|AGRB1_HUMAN Adhesion G protein-coupled receptor B1 OS=Homo sapiens OX=9606 GN=ADGRB1 PE=1 SV=2- TL EXPOSED**

STFAILAQLSADANMEKATLPSVTLIVGCGVSSL

TLLMLVIIYVSVWRYIRSERSVILINFCLSIISSNALILIGQTQTRNKVVCTLVAAFLHF

FFLSSFCWVLTEAWQSYMAVTGHLRNRLIRKRFLCLGWGLPALVVAISVGFTKAKGYSTM

NYCWLSLEGGLLYAFVGPAAAVVLVNMVIGILVFNKLVSKDGITDKKLKERAGASLWSSC

VVLPLLALTWMSAVLAVTDRRSALFQILFAVFDSLEGFVIVMVHCILRREVQDAVKCRVV

DRQEEGNGDSGGSFQNGHAQLMTDFEKDVDLACRSVLNKDIAACRTATITGTLKRPSLPE

EEKLKLAHAKGPPTNFNSLPANVSKLHLHGSPRYPGGPLPDFPNHSLTLKRDKAPKSSFV

GDGDIFKKLDSELSRAQEKALDTSYVILPTATATLRPKPKEEPKYSIHIDQMPQTRLIHL

STAPEASLPARSPPSRQPPSGGPPEAPPAQPPPPPPPPPPPPQQPLPPPPNLEPAPPSLG

DPGEPAAHPGPSTGPSTKNENVATLSVSSLERRKSRYAELDFEKIMHTRKRHQDMFQDLN

RKLQHAAEKDKEVLGPDSKPEKQQTPNKRPWESLRKAHGTPTWVKKELEPLQPSPLELRS

VEWERSGATIPLVGQDIIDLQTEV

**>sp|O60241|AGRB2_HUMAN Adhesion G protein-coupled receptor B2 OS=Homo sapiens OX=9606 GN=ADGRB2 PE=1 SV=2**

MENTGWMGKGHRMTPACPLLLSVILSLRLATAFDPAPSACSALASGVLYGAFSLQDLFPT

IASGCSWTLENPDPTKYSLYLRFNRQEQVCAHFAPRLLPLDHYLVNFTCLRPSPEEAVAQ

AESEVGRPEEEEAEAAAGLELCSGSGPFTFLHFDKNFVQLCLSAEPSEAPRLLAPAALAF

RFVEVLLINNNNSSQFTCGVLCRWSEECGRAAGRACGFAQPGCSCPGEAGAGSTTTTSPG

PPAAHTLSNALVPGGPAPPAEADLHSGSSNDLFTTEMRYGEEPEEEPKVKTQWPRSADEP

GLYMAQTGDPAAEEWSPWSVCSLTCGQGLQVRTRSCVSSPYGTLCSGPLRETRPCNNSAT

CPVHGVWEEWGSWSLCSRSCGRGSRSRMRTCVPPQHGGKACEGPELQTKLCSMAACPVEG

QWLEWGPWGPCSTSCANGTQQRSRKCSVAGPAWATCTGALTDTRECSNLECPATDSKWGP

WNAWSLCSKTCDTGWQRRFRMCQATGTQGYPCEGTGEEVKPCSEKRCPAFHEMCRDEYVM

LMTWKKAAAGEIIYNKCPPNASGSASRRCLLSAQGVAYWGLPSFARCISHEYRYLYLSLR

EHLAKGQRMLAGEGMSQVVRSLQELLARRTYYSGDLLFSVDILRNVTDTFKRATYVPSAD

DVQRFFQVVSFMVDAENKEKWDDAQQVSPGSVHLLRVVEDFIHLVGDALKAFQSSLIVTD

NLVISIQREPVSAVSSDITFPMRGRRGMKDWVRHSEDRLFLPKEVLSLSSPGKPATSGAA

GSPGRGRGPGTVPPGPGHSHQRLLPADPDESSYFVIGAVLYRTLGLILPPPRPPLAVTSR

VMTVTVRPPTQPPAEPLITVELSYIINGTTDPHCASWDYSRADASSGDWDTENCQTLETQ

AAHTRCQCQHLSTFAVLAQPPKDLTLELAGSPSVPLVIGCAVSCMALLTLLAIYAAFWRF

IKSERSIILLNFCLSILASNILILVGQSRVLSKGVCTMTAAFLHFFFLSSFCWVLTEAWQ

SYLAVIGRMRTRLVRKRFLCLGWGLPALVVAVSVGFTRTKGYGTSSYCWLSLEGGLLYAF

VGPAAVIVLVNMLIGIIVFNKLMARDGISDKSKKQRAGSERCPWASLLLPCSACGAVPSP

LLSSASARNAMASLWSSCVVLPLLALTWMSAVLAMTDRRSVLFQALFAVFNSAQGFVITA

VHCFLRREVQDVVKCQMGVCRADESEDSPDSCKNGQLQILSDFEKDVDLACQTVLFKEVN

TCNPSTITGTLSRLSLDEDEEPKSCLVGPEGSLSFSPLPGNILVPMAASPGLGEPPPPQE

ANPVYMCGEGGLRQLDLTWLRPTEPGSEGDYMVLPRRTLSLQPGGGGGGGEDAPRARPEG

TPRRAAKTVAHTEGYPSFLSVDHSGLGLGPAYGSLQNPYGMTFQPPPPTPSARQVPEPGE

RSRTMPRTVPGSTMKMGSLERKKLRYSDLDFEKVMHTRKRHSELYHELNQKFHTFDRYRS

QSTAKREKRWSVSSGGAAERSVCTDKPSPGERPSLSQHRRHQSWSTFKSMTLGSLPPKPR

ERLTLHRAAAWEPTEPPDGDFQTEV

**>sp|O60241|AGRB2_HUMAN Adhesion G protein-coupled receptor B2 OS=Homo sapiens OX=9606 GN=ADGRB2 PE=1 SV=2- TL EXPOSED**

STFAVLAQPPKDLTLELAGSPSVPLVIGCAVSCMALLTLLAIYAAFWRF

IKSERSIILLNFCLSILASNILILVGQSRVLSKGVCTMTAAFLHFFFLSSFCWVLTEAWQ

SYLAVIGRMRTRLVRKRFLCLGWGLPALVVAVSVGFTRTKGYGTSSYCWLSLEGGLLYAF

VGPAAVIVLVNMLIGIIVFNKLMARDGISDKSKKQRAGSERCPWASLLLPCSACGAVPSP

LLSSASARNAMASLWSSCVVLPLLALTWMSAVLAMTDRRSVLFQALFAVFNSAQGFVITA

VHCFLRREVQDVVKCQMGVCRADESEDSPDSCKNGQLQILSDFEKDVDLACQTVLFKEVN

TCNPSTITGTLSRLSLDEDEEPKSCLVGPEGSLSFSPLPGNILVPMAASPGLGEPPPPQE

ANPVYMCGEGGLRQLDLTWLRPTEPGSEGDYMVLPRRTLSLQPGGGGGGGEDAPRARPEG

TPRRAAKTVAHTEGYPSFLSVDHSGLGLGPAYGSLQNPYGMTFQPPPPTPSARQVPEPGE

RSRTMPRTVPGSTMKMGSLERKKLRYSDLDFEKVMHTRKRHSELYHELNQKFHTFDRYRS

QSTAKREKRWSVSSGGAAERSVCTDKPSPGERPSLSQHRRHQSWSTFKSMTLGSLPPKPR

ERLTLHRAAAWEPTEPPDGDFQTEV

**>sp|O60242|AGRB3_HUMAN Adhesion G protein-coupled receptor B3 OS=Homo sapiens OX=9606 GN=ADGRB3 PE=1 SV=2**

MKAVRNLLIYIFSTYLLVMFGFNAAQDFWCSTLVKGVIYGSYSVSEMFPKNFTNCTWTLE

NPDPTKYSIYLKFSKKDLSCSNFSLLAYQFDHFSHEKIKDLLRKNHSIMQLCNSKNAFVF

LQYDKNFIQIRRVFPTNFPGLQKKGEEDQKSFFEFLVLNKVSPSQFGCHVLCTWLESCLK

SENGRTESCGIMYTKCTCPQHLGEWGIDDQSLILLNNVVLPLNEQTEGCLTQELQTTQVC

NLTREAKRPPKEEFGMMGDHTIKSQRPRSVHEKRVPQEQADAAKFMAQTGESGVEEWSQW

STCSVTCGQGSQVRTRTCVSPYGTHCSGPLRESRVCNNTALCPVHGVWEEWSPWSLCSFT

CGRGQRTRTRSCTPPQYGGRPCEGPETHHKPCNIALCPVDGQWQEWSSWSQCSVTCSNGT

QQRSRQCTAAAHGGSECRGPWAESRECYNPECTANGQWNQWGHWSGCSKSCDGGWERRIR

TCQGAVITGQQCEGTGEEVRRCNEQRCPAPYEICPEDYLMSMVWKRTPAGDLAFNQCPLN

ATGTTSRRCSLSLHGVAFWEQPSFARCISNEYRHLQHSIKEHLAKGQRMLAGDGMSQVTK

TLLDLTQRKNFYAGDLLMSVEILRNVTDTFKRASYIPASDGVQNFFQIVSNLLDEENKEK

WEDAQQIYPGSIELMQVIEDFIHIVGMGMMDFQNSYLMTGNVVASIQKLPAASVLTDINF

PMKGRKGMVDWARNSEDRVVIPKSIFTPVSSKELDESSVFVLGAVLYKNLDLILPTLRNY

TVINSKIIVVTIRPEPKTTDSFLEIELAHLANGTLNPYCVLWDDSKTNESLGTWSTQGCK

TVLTDASHTKCLCDRLSTFAILAQQPREIIMESSGTPSVTLIVGSGLSCLALITLAVVYA

ALWRYIRSERSIILINFCLSIISSNILILVGQTQTHNKSICTTTTAFLHFFFLASFCWVL

TEAWQSYMAVTGKIRTRLIRKRFLCLGWGLPALVVATSVGFTRTKGYGTDHYCWLSLEGG

LLYAFVGPAAAVVLVNMVIGILVFNKLVSRDGILDKKLKHRAGQMSEPHSGLTLKCAKCG

VVSTTALSATTASNAMASLWSSCVVLPLLALTWMSAVLAMTDKRSILFQILFAVFDSLQG

FVIVMVHCILRREVQDAFRCRLRNCQDPINADSSSSFPNGHAQIMTDFEKDVDIACRSVL

HKDIGPCRAATITGTLSRISLNDDEEEKGTNPEGLSYSTLPGNVISKVIIQQPTGLHMPM

SMNELSNPCLKKENSELRRTVYLCTDDNLRGADMDIVHPQERMMESDYIVMPRSSVNNQP

SMKEESKMNIGMETLPHERLLHYKVNPEFNMNPPVMDQFNMNLEQHLAPQEHMQNLPFEP

RTAVKNFMASELDDNAGLSRSETGSTISMSSLERRKSRYSDLDFEKVMHTRKRHMELFQE

LNQKFQTLDRFRDIPNTSSMENPAPNKNPWDTFKNPSEYPHYTTINVLDTEAKDALELRP

AEWEKCLNLPLDVQEGDFQTEV

**>sp|O60242|AGRB3_HUMAN Adhesion G protein-coupled receptor B3 OS=Homo sapiens OX=9606 GN=ADGRB3 PE=1 SV=2- TL EXPOSED**

STFAILAQQPREIIMESSGTPSVTLIVGSGLSCLALITLAVVYA

ALWRYIRSERSIILINFCLSIISSNILILVGQTQTHNKSICTTTTAFLHFFFLASFCWVL

TEAWQSYMAVTGKIRTRLIRKRFLCLGWGLPALVVATSVGFTRTKGYGTDHYCWLSLEGG

LLYAFVGPAAAVVLVNMVIGILVFNKLVSRDGILDKKLKHRAGQMSEPHSGLTLKCAKCG

VVSTTALSATTASNAMASLWSSCVVLPLLALTWMSAVLAMTDKRSILFQILFAVFDSLQG

FVIVMVHCILRREVQDAFRCRLRNCQDPINADSSSSFPNGHAQIMTDFEKDVDIACRSVL

HKDIGPCRAATITGTLSRISLNDDEEEKGTNPEGLSYSTLPGNVISKVIIQQPTGLHMPM

SMNELSNPCLKKENSELRRTVYLCTDDNLRGADMDIVHPQERMMESDYIVMPRSSVNNQP

SMKEESKMNIGMETLPHERLLHYKVNPEFNMNPPVMDQFNMNLEQHLAPQEHMQNLPFEP

RTAVKNFMASELDDNAGLSRSETGSTISMSSLERRKSRYSDLDFEKVMHTRKRHMELFQE

LNQKFQTLDRFRDIPNTSSMENPAPNKNPWDTFKNPSEYPHYTTINVLDTEAKDALELRP

AEWEKCLNLPLDVQEGDFQTEV

**>sp|Q6QNK2|AGRD1_HUMAN Adhesion G-protein coupled receptor D1 OS=Homo sapiens OX=9606 GN=ADGRD1 PE=1 SV=1**

MEKLLRLCCWYSWLLLFYYNFQVRGVYSRSQDHPGFQVLASASHYWPLENVDGIHELQDT

TGDIVEGKVNKGIYLKEEKGVTLLYYGRYNSSCISKPEQCGPEGVTFSFFWKTQGEQSRP

IPSAYGGQVISNGFKVCSSGGRGSVELYTRDNSMTWEASFSPPGPYWTHVLFTWKSKEGL

KVYVNGTLSTSDPSGKVSRDYGESNVNLVIGSEQDQAKCYENGAFDEFIIWERALTPDEI

AMYFTAAIGKHALLSSTLPSLFMTSTASPVMPTDAYHPIITNLTEERKTFQSPGVILSYL

QNVSLSLPSKSLSEQTALNLTKTFLKAVGEILLLPGWIALSEDSAVVLSLIDTIDTVMGH

VSSNLHGSTPQVTVEGSSAMAEFSVAKILPKTVNSSHYRFPAHGQSFIQIPHEAFHRHAW

STVVGLLYHSMHYYLNNIWPAHTKIAEAMHHQDCLLFATSHLISLEVSPPPTLSQNLSGS

PLITVHLKHRLTRKQHSEATNSSNRVFVYCAFLDFSSGEGVWSNHGCALTRGNLTYSVCR

CTHLTNFAILMQVVPLELARGHQVALSSISYVGCSLSVLCLVATLVTFAVLSSVSTIRNQ

RYHIHANLSFAVLVAQVLLLISFRLEPGTTPCQVMAVLLHYFFLSAFAWMLVEGLHLYSM

VIKVFGSEDSKHRYYYGMGWGFPLLICIISLSFAMDSYGTSNNCWLSLASGAIWAFVAPA

LFVIVVNIGILIAVTRVISQISADNYKIHGDPSAFKLTAKAVAVLLPILGTSWVFGVLAV

NGCAVVFQYMFATLNSLQGLFIFLFHCLLNSEVRAAFKHKTKVWSLTSSSARTSNAKPFH

SDLMNGTRPGMASTKLSPWDKSSHSAHRVDLSAV

**>sp|Q6QNK2|AGRD1_HUMAN Adhesion G-protein coupled receptor D1 OS=Homo sapiens OX=9606 GN=ADGRD1 PE=1 SV=1 – TL EXPOSED**

TNFAILMQVVPLELARGHQVALSSISYVGCSLSVLCLVATLVTFAVLSSVSTIRNQ

RYHIHANLSFAVLVAQVLLLISFRLEPGTTPCQVMAVLLHYFFLSAFAWMLVEGLHLYSM

VIKVFGSEDSKHRYYYGMGWGFPLLICIISLSFAMDSYGTSNNCWLSLASGAIWAFVAPA

LFVIVVNIGILIAVTRVISQISADNYKIHGDPSAFKLTAKAVAVLLPILGTSWVFGVLAV

NGCAVVFQYMFATLNSLQGLFIFLFHCLLNSEVRAAFKHKTKVWSLTSSSARTSNAKPFH

SDLMNGTRPGMASTKLSPWDKSSHSAHRVDLSAV

**>sp|Q7Z7M1|AGRD2_HUMAN Adhesion G-protein coupled receptor D2 OS=Homo sapiens OX=9606 GN=ADGRD2 PE=2 SV=1**

MDAPWGAGERWLHGAAVDRSGVSLGPPPTPQVNQGTLGPQVAPVAAGEVVKTAGGVCKFS

GQRLSWWQAQESCEQQFGHLALQPPDGVLASRLRDPVWVGQREAPLRRPPQRRARTTAVL

VFDERTADRAARLRSPLPELAALTACTHVQWDCASPDPAALFSVAAPALPNALQLRAFAE

PGGVVRAALVVRGQHAPFLAAFRADGRWHHVCATWEQRGGRWALFSDGRRRAGARGLGAG

HPVPSGGILVLGQDQDSLGGGFSVRHALSGNLTDFHLWARALSPAQLHRARACAPPSEGL

LFRWDPGALDVTPSLLPTVWVRLLCPVPSEECPTWNPGPRSEGSELCLEPQPFLCCYRTE

PYRRLQDAQSWPGQDVISRVNALANDIVLLPDPLSEVHGALSPAEASSFLGLLEHVLAME

MAPLGPAALLAVVRFLKRVVALGAGDPELLLTGPWEQLSQGVVSVASLVLEEQVADTWLS

LREVIGGPMALVASVQRLAPLLSTSMTSERPRMRIQHRHAGLSGVTVIHSWFTSRVFQHT

LEGPDLEPQAPASSEEANRVQRFLSTQVGSAIISSEVWDVTGEVNVAMTFHLQHRAQSPL

FPPHPPSPYTGGAWATTGCSVAALYLDSTACFCNHSTSFAILLQIYEVQRGPEEESLLRT

LSFVGCGVSFCALTTTFLLFLVAGVPKSERTTVHKNLTFSLASAEGFLMTSEWAKANEVA

CVAVTVAMHFLFLVAFSWMLVEGLLLWRKVVAVSMHPGPGMRLYHATGWGVPVGIVAVTL

AMLPHDYVAPGHCWLNVHTNAIWAFVGPVLFVLTANTCILARVVMITVSSARRRARMLSP

QPCLQQQIWTQIWATVKPVLVLLPVLGLTWLAGILVHLSPAWAYAAVGLNSIQGLYIFLV

YAACNEEVRSALQRMAEKKVAEVLRALGVWGGAAKEHSLPFSVLPLFLPPKPSTPRHPLK

APA

**>sp|Q7Z7M1|AGRD2_HUMAN Adhesion G-protein coupled receptor D2 OS=Homo sapiens OX=9606 GN=ADGRD2 PE=2 SV=1**

TSFAILLQIYEVQRGPEEESLLRTLSFVGCGVSFCALTTTFLLFLVAGVPKSERTTVHKN

LTFSLASAEGFLMTSEWAKANEVA CVAVTVAMHFLFLVAFSWMLVEGLLLWRKVVAVSM

HPGPGMRLYHATGWGVPVGIVAVTLAMLPHDYVAPGHCWLNVHTNAIWAFVGPVLFVLTA

NTCILARVVMITVSSARRRARMLSP QPCLQQQIWTQIWATVKPVLVLLPVLGLTWLAGI

LVHLSPAWAYAAVGLNSIQGLYIFLVYAACNEEVRSALQRMAEKKVAEVLRALGVWGGAA

KEHSLPFSVLPLFLPPKPSTPRHPLKAPA

**>sp|Q14246|AGRE1_HUMAN Adhesion G protein-coupled receptor E1 OS=Homo sapiens OX=9606 GN=ADGRE1 PE=2 SV=3- TL EXPOSED**

MRGFNLLLFWGCCVMHSWEGHIRPTRKPNTKGNNCRDSTLCPAYATCTNTVDSYYCACKQ

GFLSSNGQNHFKDPGVRCKDIDECSQSPQPCGPNSSCKNLSGRYKCSCLDGFSSPTGNDW

VPGKPGNFSCTDINECLTSSVCPEHSDCVNSMGSYSCSCQVGFISRNSTCEDVDECADPR

ACPEHATCNNTVGNYSCFCNPGFESSSGHLSFQGLKASCEDIDECTEMCPINSTCTNTPG

SYFCTCHPGFAPSNGQLNFTDQGVECRDIDECRQDPSTCGPNSICTNALGSYSCGCIAGF

HPNPEGSQKDGNFSCQRVLFKCKEDVIPDNKQIQQCQEGTAVKPAYVSFCAQINNIFSVL

DKVCENKTTVVSLKNTTESFVPVLKQISTWTKFTKEETSSLATVFLESVESMTLASFWKP

SANITPAVRTEYLDIESKVINKECSEENVTLDLVAKGDKMKIGCSTIEESESTETTGVAF

VSFVGMESVLNERFFKDHQAPLTTSEIKLKMNSRVVGGIMTGEKKDGFSDPIIYTLENIQ

PKQKFERPICVSWSTDVKGGRWTSFGCVILEASETYTICSCNQMANLAVIMASGELTMDF

SLYIISHVGIIISLVCLVLAIATFLLCRSIRNHNTYLHLHLCVCLLLAKTLFLAGIHKTD

NKMGCAIIAGFLHYLFLACFFWMLVEAVILFLMVRNLKVVNYFSSRNIKMLHICAFGYGL

PMLVVVISASVQPQGYGMHNRCWLNTETGFIWSFLGPVCTVIVINSLLLTWTLWILRQRL

SSVNAEVSTLKDTRLLTFKAFAQLFILGCSWVLGIFQIGPVAGVMAYLFTIINSLQGAFI

FLIHCLLNGQVREEYKRWITGKTKPSSQSQTSRILLSSMPSASKTG

**>sp|Q14246|AGRE1_HUMAN Adhesion G protein-coupled receptor E1 OS=Homo sapiens OX=9606 GN=ADGRE1 PE=2 SV=3- TL EXPOSED**

ANLAVIMASGELTMDFSLYIISHVGIIISLVCLVLAIATFLLCRSIRNHNTYLHLHLCVC

LLLAKTLFLAGIHKTDNKMGCAIIAGFLHYLFLACFFWMLVEAVILFLMVRNLKVVNYFS

SRNIKMLHICAFGYGLPMLVVVISASVQPQGYGMHNRCWLNTETGFIWSFLGPVCTVIVI

NSLLLTWTLWILRQRLSSVNAEVSTLKDTRLLTFKAFAQLFILGCSWVLGIFQIGPVAGV

MAYLFTIINSLQGAFIFLIHCLLNGQVREEYKRWITGKTKPSSQSQTSRILLSSMPSASK

TG

**>sp|Q9UHX3|AGRE2_HUMAN Adhesion G protein-coupled receptor E2 OS=Homo sapiens OX=9606 GN=ADGRE2 PE=1 SV=2**

MGGRVFLVFLAFCVWLTLPGAETQDSRGCARWCPQDSSCVNATACRCNPGFSSFSEIITT

PMETCDDINECATLSKVSCGKFSDCWNTEGSYDCVCSPGYEPVSGAKTFKNESENTCQDV

DECQQNPRLCKSYGTCVNTLGSYTCQCLPGFKLKPEDPKLCTDVNECTSGQNPCHSSTHC

LNNVGSYQCRCRPGWQPIPGSPNGPNNTVCEDVDECSSGQHQCDSSTVCFNTVGSYSCRC

RPGWKPRHGIPNNQKDTVCEDMTFSTWTPPPGVHSQTLSRFFDKVQDLGRDYKPGLANNT

IQSILQALDELLEAPGDLETLPRLQQHCVASHLLDGLEDVLRGLSKNLSNGLLNFSYPAG

TELSLEVQKQVDRSVTLRQNQAVMQLDWNQAQKSGDPGPSVVGLVSIPGMGKLLAEAPLV

LEPEKQMLLHETHQGLLQDGSPILLSDVISAFLSNNDTQNLSSPVTFTFSHRSVIPRQKV

LCVFWEHGQNGCGHWATTGCSTIGTRDTSTICRCTHLSSFAVLMAHYDVQEEDPVLTVIT

YMGLSVSLLCLLLAALTFLLCKAIQNTSTSLHLQLSLCLFLAHLLFLVAIDQTGHKVLCS

IIAGTLHYLYLATLTWMLLEALYLFLTARNLTVVNYSSINRFMKKLMFPVGYGVPAVTVA

ISAASRPHLYGTPSRCWLQPEKGFIWGFLGPVCAIFSVNLVLFLVTLWILKNRLSSLNSE

VSTLRNTRMLAFKATAQLFILGCTWCLGILQVGPAARVMAYLFTIINSLQGVFIFLVYCL

LSQQVREQYGKWSKGIRKLKTESEMHTLSSSAKADTSKPSTVN

**>sp|Q9UHX3|AGRE2_HUMAN Adhesion G protein-coupled receptor E2 OS=Homo sapiens OX=9606 GN=ADGRE2 PE=1 SV=2- TL EXPOSED**

SSFAVLMAHYDVQEEDPVLTVITYMGLSVSLLCLLLAALTFLLCKAIQNTSTSLHLQLSL

CLFLAHLLFLVAIDQTGHKVLCSIIAGTLHYLYLATLTWMLLEALYLFLTARNLTVVNYS

SINRFMKKLMFPVGYGVPAVTVAISAASRPHLYGTPSRCWLQPEKGFIWGFLGPVCAIFS

VNLVLFLVTLWILKNRLSSLNSE VSTLRNTRMLAFKATAQLFILGCTWCLGILQVGPAA

RVMAYLFTIINSLQGVFIFLVYCLLSQQVREQYGKWSKGIRKLKTESEMHTLSSSAKADT

SKPSTVN

**>sp|Q9BY15|AGRE3_HUMAN Adhesion G protein-coupled receptor E3 OS=Homo sapiens OX=9606 GN=ADGRE3 PE=2 SV=2**

MQGPLLLPGLCFLLSLFGAVTQKTKTSCAKCPPNASCVNNTHCTCNHGYTSGSGQKLFTF

PLETCNDINECTPPYSVYCGFNAVCYNVEGSFYCQCVPGYRLHSGNEQFSNSNENTCQDT

TSSKTTEGRKELQKIVDKFESLLTNQTLWRTEGRQEISSTATTILRDVESKVLETALKDP

EQKVLKIQNDSVAIETQAITDNCSEERKTFNLNVQMNSMDIRCSDIIQGDTQGPSAIAFI

SYSSLGNIINATFFEEMDKKDQVYLNSQVVSAAIGPKRNVSLSKSVTLTFQHVKMTPSTK

KVFCVYWKSTGQGSQWSRDGCFLIHVNKSHTMCNCSHLSSFAVLMALTSQEEDPVLTVIT

YVGLSVSLLCLLLAALTFLLCKAIRNTSTSLHLQLSLCLFLAHLLFLVGIDRTEPKVLCS

IIAGALHYLYLAAFTWMLLEGVHLFLTARNLTVVNYSSINRLMKWIMFPVGYGVPAVTVA

ISAASWPHLYGTADRCWLHLDQGFMWSFLGPVCAIFSANLVLFILVFWILKRKLSSLNSE

VSTIQNTRMLAFKATAQLFILGCTWCLGLLQVGPAAQVMAYLFTIINSLQGFFIFLVYCL

LSQQVQKQYQKWFREIVKSKSESETYTLSSKMGPDSKPSEGDVFPGQVKRKY

**>sp|Q9BY15|AGRE3_HUMAN Adhesion G protein-coupled receptor E3 OS=Homo sapiens OX=9606 GN=ADGRE3 PE=2 SV=2 – TL EXPOSED**

SSFAVLMALTSQEEDPVLTVIT

YVGLSVSLLCLLLAALTFLLCKAIRNTSTSLHLQLSLCLFLAHLLFLVGIDRTEPKVLCS

IIAGALHYLYLAAFTWMLLEGVHLFLTARNLTVVNYSSINRLMKWIMFPVGYGVPAVTVA

ISAASWPHLYGTADRCWLHLDQGFMWSFLGPVCAIFSANLVLFILVFWILKRKLSSLNSE

VSTIQNTRMLAFKATAQLFILGCTWCLGLLQVGPAAQVMAYLFTIINSLQGFFIFLVYCL

LSQQVQKQYQKWFREIVKSKSESETYTLSSKMGPDSKPSEGDVFPGQVKRKY

**>sp|Q86SQ3|AGRE4_HUMAN Putative adhesion G protein-coupled receptor E4P OS=Homo sapiens OX=9606 GN=ADGRE4P PE=5 SV=1**

MGSRFLLVLLSGASCPPCPKYASCHNSTHCTCEDGFRARSGRTYFHDSSEKCEDINECET

GLAKCKYKAYCRNKVGGYICSCLVKYTLFNFLAGIIDYDHPDCYENNSQGTTQSNVDIWV

SGVKPGFGKQLPGDKRTKHICVYWEGSEGGWSTEGCSHVHSNGSYTKCKCFHLSSFAVLV

ALAPKEDPVLTVITQVGLTISLLCLFLAILTFLLCRPIQNTSTSLHLELSLCLFLAHLLF

LTGINRTEPEVLCSIIAGLLHFLYLACFTWMLLEGLHLFLTVRNLKVANYTSTGRFKKRF

MYPVGYGIPAVIIAVSAIVGPQNYGTFTCWLKLDKGFIWSFMGPVAVIILINLVFYFQVL

WILRSKLSSLNKEVSTIQDTRVMTFKAISQLFILGCSWGLGFFMVEEVGKTIGSIIAYSF

TIINTLQGVLLFVVHCLLNRQVRLIILSVISLVPKSN

**>sp|Q86SQ3|AGRE4_HUMAN Putative adhesion G protein-coupled receptor E4P OS=Homo sapiens OX=9606 GN=ADGRE4P PE=5 SV=1 – TL EXPOSED**

SSFAVLVALAPKEDPVLTVITQVGLTISLLCLFLAILTFLLCRPIQNTSTSLHLELSLCLFLAHLLF

LTGINRTEPEVLCSIIAGLLHFLYLACFTWMLLEGLHLFLTVRNLKVANYTSTGRFKKRF

MYPVGYGIPAVIIAVSAIVGPQNYGTFTCWLKLDKGFIWSFMGPVAVIILINLVFYFQVL

WILRSKLSSLNKEVSTIQDTRVMTFKAISQLFILGCSWGLGFFMVEEVGKTIGSIIAYSF

TIINTLQGVLLFVVHCLLNRQVRLIILSVISLVPKSN

**>sp|P48960|AGRE5_HUMAN Adhesion G protein-coupled receptor E5 OS=Homo sapiens OX=9606 GN=ADGRE5 PE=1 SV=4**

MGGRVFLAFCVWLTLPGAETQDSRGCARWCPQNSSCVNATACRCNPGFSSFSEIITTPTE

TCDDINECATPSKVSCGKFSDCWNTEGSYDCVCSPGYEPVSGAKTFKNESENTCQDVDEC

QQNPRLCKSYGTCVNTLGSYTCQCLPGFKFIPEDPKVCTDVNECTSGQNPCHSSTHCLNN

VGSYQCRCRPGWQPIPGSPNGPNNTVCEDVDECSSGQHQCDSSTVCFNTVGSYSCRCRPG

WKPRHGIPNNQKDTVCEDMTFSTWTPPPGVHSQTLSRFFDKVQDLGRDSKTSSAEVTIQN

VIKLVDELMEAPGDVEALAPPVRHLIATQLLSNLEDIMRILAKSLPKGPFTYISPSNTEL

TLMIQERGDKNVTMGQSSARMKLNWAVAAGAEDPGPAVAGILSIQNMTTLLANASLNLHS

KKQAELEEIYESSIRGVQLRRLSAVNSIFLSHNNTKELNSPILFAFSHLESSDGEAGRDP

PAKDVMPGPRQELLCAFWKSDSDRGGHWATEGCQVLGSKNGSTTCQCSHLSSFAILMAHY

DVEDWKLTLITRVGLALSLFCLLLCILTFLLVRPIQGSRTTIHLHLCICLFVGSTIFLAG

IENEGGQVGLRCRLVAGLLHYCFLAAFCWMSLEGLELYFLVVRVFQGQGLSTRWLCLIGY

GVPLLIVGVSAAIYSKGYGRPRYCWLDFEQGFLWSFLGPVTFIILCNAVIFVTTVWKLTQ

KFSEINPDMKKLKKARALTITAIAQLFLLGCTWVFGLFIFDDRSLVLTYVFTILNCLQGA

FLYLLHCLLNKKVREEYRKWACLVAGGSKYSEFTSTTSGTGHNQTRALRASESGI

**>sp|P48960|AGRE5_HUMAN Adhesion G protein-coupled receptor E5 OS=Homo sapiens OX=9606 GN=ADGRE5 PE=1 SV=4 – TL EXPOSED**

SSFAILMAHYDVEDWKLTLITRVGLALSLFCLLLCILTFLLVRPIQGSRTTIHLHLCICLFVGSTIFLAG

IENEGGQVGLRCRLVAGLLHYCFLAAFCWMSLEGLELYFLVVRVFQGQGLSTRWLCLIGY

GVPLLIVGVSAAIYSKGYGRPRYCWLDFEQGFLWSFLGPVTFIILCNAVIFVTTVWKLTQ

KFSEINPDMKKLKKARALTITAIAQLFLLGCTWVFGLFIFDDRSLVLTYVFTILNCLQGA

FLYLLHCLLNKKVREEYRKWACLVAGGSKYSEFTSTTSGTGHNQTRALRASESGI

**>sp|Q5T601|AGRF1_HUMAN Adhesion G-protein coupled receptor F1 OS=Homo sapiens OX=9606 GN=ADGRF1 PE=1 SV=2**

MKVGVLWLISFFTFTDGHGGFLGKNDGIKTKKELIVNKKKHLGPVEEYQLLLQVTYRDSK

EKRDLRNFLKLLKPPLLWSHGLIRIIRAKATTDCNSLNGVLQCTCEDSYTWFPPSCLDPQ

NCYLHTAGALPSCECHLNNLSQSVNFCERTKIWGTFKINERFTNDLLNSSSAIYSKYANG

IEIQLKKAYERIQGFESVQVTQFRNGSIVAGYEVVGSSSASELLSAIEHVAEKAKTALHK

LFPLEDGSFRVFGKAQCNDIVFGFGSKDDEYTLPCSSGYRGNITAKCESSGWQVIRETCV

LSLLEELNKNFSMIVGNATEAAVSSFVQNLSVIIRQNPSTTVGNLASVVSILSNISSLSL

ASHFRVSNSTMEDVISIADNILNSASVTNWTVLLREEKYASSRLLETLENISTLVPPTAL

PLNFSRKFIDWKGIPVNKSQLKRGYSYQIKMCPQNTSIPIRGRVLIGSDQFQRSLPETII

SMASLTLGNILPVSKNGNAQVNGPVISTVIQNYSINEVFLFFSKIESNLSQPHCVFWDFS

HLQWNDAGCHLVNETQDIVTCQCTHLTSFSILMSPFVPSTIFPVVKWITYVGLGISIGSL

ILCLIIEALFWKQIKKSQTSHTRRICMVNIALSLLIADVWFIVGATVDTTVNPSGVCTAA

VFFTHFFYLSLFFWMLMLGILLAYRIILVFHHMAQHLMMAVGFCLGYGCPLIISVITIAV

TQPSNTYKRKDVCWLNWSNGSKPLLAFVVPALAIVAVNFVVVLLVLTKLWRPTVGERLSR

DDKATIIRVGKSLLILTPLLGLTWGFGIGTIVDSQNLAWHVIFALLNAFQGFFILCFGIL

LDSKLRQLLFNKLSALSSWKQTEKQNSSDLSAKPKFSKPFNPLQNKGHYAFSHTGDSSDN

IMLTQFVSNE

**>sp|Q5T601|AGRF1_HUMAN Adhesion G-protein coupled receptor F1 OS=Homo sapiens OX=9606 GN=ADGRF1 PE=1 SV=2** **– TL EXPOSED**

TSFSILMSPFVPSTIFPVVKWITYVGLGISIGSL

ILCLIIEALFWKQIKKSQTSHTRRICMVNIALSLLIADVWFIVGATVDTTVNPSGVCTAA

VFFTHFFYLSLFFWMLMLGILLAYRIILVFHHMAQHLMMAVGFCLGYGCPLIISVITIAV

TQPSNTYKRKDVCWLNWSNGSKPLLAFVVPALAIVAVNFVVVLLVLTKLWRPTVGERLSR

DDKATIIRVGKSLLILTPLLGLTWGFGIGTIVDSQNLAWHVIFALLNAFQGFFILCFGIL

LDSKLRQLLFNKLSALSSWKQTEKQNSSDLSAKPKFSKPFNPLQNKGHYAFSHTGDSSDN

IMLTQFVSNE

**>sp|Q8IZF7|AGRF2_HUMAN Adhesion G-protein coupled receptor F2 OS=Homo sapiens OX=9606 GN=ADGRF2 PE=2 SV=1**

MGLTAYGNRRVQPGELPFGANLTLIHTRAQPVICSKLLLTKRVSPISFFLSKFQNSWGED

GWVQLDQLPSPNAVSSDQVHCSAGCTHRKCGWAASKSKEKVPARPHGVCDGVCTDYSQCT

QPCPPDTQGNMGFSCRQKTWHKITDTCQTLNALNIFEEDSRLVQPFEDNIKISVYTGKSE

TITDMLLQKCPTDLSCVIRNIQQSPWIPGNIAVIVQLLHNISTAIWTGVDEAKMQSYSTI

ANHILNSKSISNWTFIPDRNSSYILLHSVNSFARRLFIDKHPVDISDVFIHTMGTTISGD

NIGKNFTFSMRINDTSNEVTGRVLISRDELRKVPSPSQVISIAFPTIGAILEASLLENVT

VNGLVLSAILPKELKRISLIFEKISKSEERRTQCVGWHSVENRWDQQACKMIQENSQQAV

CKCRPSKLFTSFSILMSPHILESLILTYITYVGLGISICSLILCLSIEVLVWSQVTKTEI

TYLRHVCIVNIAATLLMADVWFIVASFLSGPITHHKGCVAATFFVHFFYLSVFFWMLAKA

LLILYGIMIVFHTLPKSVLVASLFSVGYGCPLAIAAITVAATEPGKGYLRPEICWLNWDM

TKALLAFVIPALAIVVVNLITVTLVIVKTQRAAIGNSMFQEVRAIVRISKNIAILTPLLG

LTWGFGVATVIDDRSLAFHIIFSLLNAFQVSPDASDQVQSERIHEDVL

**>sp|Q8IZF7|AGRF2_HUMAN Adhesion G-protein coupled receptor F2 OS=Homo sapiens OX=9606 GN=ADGRF2 PE=2 SV=1 – TL EXPOSED**

TSFSILMSPHILESLILTYITYVGLGISICSLILCLSIEVLVWSQVTKTEI

TYLRHVCIVNIAATLLMADVWFIVASFLSGPITHHKGCVAATFFVHFFYLSVFFWMLAKA

LLILYGIMIVFHTLPKSVLVASLFSVGYGCPLAIAAITVAATEPGKGYLRPEICWLNWDM

TKALLAFVIPALAIVVVNLITVTLVIVKTQRAAIGNSMFQEVRAIVRISKNIAILTPLLG

LTWGFGVATVIDDRSLAFHIIFSLLNAFQVSPDASDQVQSERIHEDVL

**>sp|Q8IZF5|AGRF3_HUMAN Adhesion G-protein coupled receptor F3 OS=Homo sapiens OX=9606 GN=ADGRF3 PE=2 SV=1**

MVCSAAPLLLLATTLPLLGSPVAQASQPVSETGVRPREGLQRRQWGPLIGRDKAWNERID

RPFPACPIPLSSSFGRWPKGQTMWAQTSTLTLTEEELGQSQAGGESGSGQLLDQENGAGE

SALVSVYVHLDFPDKTWPPELSRTLTLPAASASSSPRPLLTGLRLTTECNVNHKGNFYCA

CLSGYQWNTSICLHYPPCQSLHNHQPCGCLVFSHPEPGYCQLLPPGSPVTCLPAVPGILN

LNSQLQMPGDTLSLTLHLSQEATNLSWFLRHPGSPSPILLQPGTQVSVTSSHGQAALSVS

NMSHHWAGEYMSCFEAQGFKWNLYEVVRVPLKATDVARLPYQLSISCATSPGFQLSCCIP

STNLAYTAAWSPGEGSKASSFNESGSQCFVLAVQRCPMADTTYACDLQSLGLAPLRVPIS

ITIIQDGDITCPEDASVLTWNVTKAGHVAQAPCPESKRGIVRRLCGADGVWGPVHSSCTD

ARLLALFTRTKLLQAGQGSPAEEVPQILAQLPGQAAEASSPSDLLTLLSTMKYVAKVVAE

ARIQLDRRALKNLLIATDKVLDMDTRSLWTLAQARKPWAGSTLLLAVETLACSLCPQDHP

FAFSLPNVLLQSQLFGPTFPADYSISFPTRPPLQAQIPRHSLAPLVRNGTEISITSLVLR

KLDHLLPSNYGQGLGDSLYATPGLVLVISIMAGDRAFSQGEVIMDFGNTDGSPHCVFWDH

SLFQGRGGWSKEGCQAQVASASPTAQCLCQHLTAFSVLMSPHTVPEEPALALLTQVGLGA

SILALLVCLGVYWLVWRVVVRNKISYFRHAALLNMVFCLLAADTCFLGAPFLSPGPRSPL

CLAAAFLCHFLYLATFFWMLAQALVLAHQLLFVFHQLAKHRVLPLMVLLGYLCPLGLAGV

TLGLYLPQGQYLREGECWLDGKGGALYTFVGPVLAIIGVNGLVLAMAMLKLLRPSLSEGP

PAEKRQALLGVIKALLILTPIFGLTWGLGLATLLEEVSTVPHYIFTILNTLQGVFILLFG

CLMDRKIQEALRKRFCRAQAPSSTISLVSCCLQILSCASKSMSEGIPWPSSEDMGTARS

**>sp|Q8IZF5|AGRF3_HUMAN Adhesion G-protein coupled receptor F3 OS=Homo sapiens OX=9606 GN=ADGRF3 PE=2 SV=1- TL EXPOSED**

TAFSVLMSPHTVPEEPALALLTQVGLGASILALLVCLGVYWLVWRVVVRNKISYFRHAAL

LNMVFCLLAADTCFLGAPFLSPGPRSPLCLAAAFLCHFLYLATFFWMLAQALVLAHQLLF

VFHQLAKHRVLPLMVLLGYLCPLGLAGVTLGLYLPQGQYLREGECWLDGKGGALYTFVGP

VLAIIGVNGLVLAMAMLKLLRPSLSEGPPAEKRQALLGVIKALLILTPIFGLTWGLGLAT

LLEEVSTVPHYIFTILNTLQGVFILLFGCLMDRKIQEALRKRFCRAQAPSSTISLVSCCL

QILSCASKSMSEGIPWPSSEDMGTARS

**>sp|Q8IZF3|AGRF4_HUMAN Adhesion G protein-coupled receptor F4 OS=Homo sapiens OX=9606 GN=ADGRF4 PE=1 SV=3**

MKMKSQATMICCLVFFLSTECSHYRSKIHLKAGDKLQSPEGKPKTGRIQEKCEGPCISSS

NCSQPCAKDFHGEIGFTCNQKKWQKSAETCTSLSVEKLFKDSTGASRLSVAAPSIPLHIL

DFRAPETIESVAQGIRKNCPFDYACITDMVKSSETTSGNIAFIVELLKNISTDLSDNVTR

EKMKSYSEVANHILDTAAISNWAFIPNKNASSDLLQSVNLFARQLHIHNNSENIVNELFI

QTKGFHINHNTSEKSLNFSMSMNNTTEDILGMVQIPRQELRKLWPNASQAISIAFPTLGA

ILREAHLQNVSLPRQVNGLVLSVVLPERLQEIILTFEKINKTRNARAQCVGWHSKKRRWD

EKACQMMLDIRNEVKCRCNYTSVVMSFSILMSSKSMTDKVLDYITCIGLSVSILSLVLCL

IIEATVWSRVVVTEISYMRHVCIVNIAVSLLTANVWFIIGSHFNIKAQDYNMCVAVTFFS

HFFYLSLFFWMLFKALLIIYGILVIFRRMMKSRMMVIGFAIGYGCPLIIAVTTVAITEPE

KGYMRPEACWLNWDNTKALLAFAIPAFVIVAVNLIVVLVVAVNTQRPSIGSSKSQDVVII

MRISKNVAILTPLLGLTWGFGIATLIEGTSLTFHIIFALLNAFQGFFILLFGTIMDHKIR

DALRMRMSSLKGKSRAAENASLGPTNGSKLMNRQG

**>sp|Q8IZF3|AGRF4_HUMAN Adhesion G protein-coupled receptor F4 OS=Homo sapiens OX=9606 GN=ADGRF4 PE=1 SV=3- TL EXPOSED**

MSFSILMSSKSMTDKVLDYITCIGLSVSILSLVLCLIIEATVWSRVVVTEISYMRHVCIV

NIAVSLLTANVWFIIGSHFNIKAQDYNMCVAVTFFSHFFYLSLFFWMLFKALLIIYGILV

IFRRMMKSRMMVIGFAIGYGCPLIIAVTTVAITEPEKGYMRPEACWLNWDNTKALLAFAI

PAFVIVAVNLIVVLVVAVNTQRPSIGSSKSQDVVIIMRISKNVAILTPLLGLTWGFGIAT

LIEGTSLTFHIIFALLNAFQGFFILLFGTIMDHKIRDALRMRMSSLKGKSRAAENASLGP

TNGSKLMNRQG

**>sp|Q8IZF2|AGRF5_HUMAN Adhesion G protein-coupled receptor F5 OS=Homo sapiens OX=9606 GN=ADGRF5 PE=1 SV=3**

MKSPRRTTLCLMFIVIYSSKAALNWNYESTIHPLSLHEHEPAGEEALRQKRAVATKSPTA

EEYTVNIEISFENASFLDPIKAYLNSLSFPIHGNNTDQITDILSINVTTVCRPAGNEIWC

SCETGYGWPRERCLHNLICQERDVFLPGHHCSCLKELPPNGPFCLLQEDVTLNMRVRLNV

GFQEDLMNTSSALYRSYKTDLETAFRKGYGILPGFKGVTVTGFKSGSVVVTYEVKTTPPS

LELIHKANEQVVQSLNQTYKMDYNSFQAVTINESNFFVTPEIIFEGDTVSLVCEKEVLSS

NVSWRYEEQQLEIQNSSRFSIYTALFNNMTSVSKLTIHNITPGDAGEYVCKLILDIFEYE

CKKKIDVMPIQILANEEMKVMCDNNPVSLNCCSQGNVNWSKVEWKQEGKINIPGTPETDI

DSSCSRYTLKADGTQCPSGSSGTTVIYTCEFISAYGARGSANIKVTFISVANLTITPDPI

SVSEGQNFSIKCISDVSNYDEVYWNTSAGIKIYQRFYTTRRYLDGAESVLTVKTSTREWN

GTYHCIFRYKNSYSIATKDVIVHPLPLKLNIMVDPLEATVSCSGSHHIKCCIEEDGDYKV

TFHTGSSSLPAAKEVNKKQVCYKHNFNASSVSWCSKTVDVCCHFTNAANNSVWSPSMKLN

LVPGENITCQDPVIGVGEPGKVIQKLCRFSNVPSSPESPIGGTITYKCVGSQWEEKRNDC

ISAPINSLLQMAKALIKSPSQDEMLPTYLKDLSISIDKAEHEISSSPGSLGAIINILDLL

STVPTQVNSEMMTHVLSTVNVILGKPVLNTWKVLQQQWTNQSSQLLHSVERFSQALQSGD

SPPLSFSQTNVQMSSMVIKSSHPETYQQRFVFPYFDLWGNVVIDKSYLENLQSDSSIVTM

AFPTLQAILAQDIQENNFAESLVMTTTVSHNTTMPFRISMTFKNNSPSGGETKCVFWNFR

LANNTGGWDSSGCYVEEGDGDNVTCICDHLTSFSILMSPDSPDPSSLLGILLDIISYVGV

GFSILSLAACLVVEAVVWKSVTKNRTSYMRHTCIVNIAASLLVANTWFIVVAAIQDNRYI

LCKTACVAATFFIHFFYLSVFFWMLTLGLMLFYRLVFILHETSRSTQKAIAFCLGYGCPL

AISVITLGATQPREVYTRKNVCWLNWEDTKALLAFAIPALIIVVVNITITIVVITKILRP

SIGDKPCKQEKSSLFQISKSIGVLTPLLGLTWGFGLTTVFPGTNLVFHIIFAILNVFQGL

FILLFGCLWDLKVQEALLNKFSLSRWSSQHSKSTSLGSSTPVFSMSSPISRRFNNLFGKT

GTYNVSTPEATSSSLENSSSASSLLN

**>sp|Q8IZF2|AGRF5_HUMAN Adhesion G protein-coupled receptor F5 OS=Homo sapiens OX=9606 GN=ADGRF5 PE=1 SV=3- TL EXPOSED**

TSFSILMSPDSPDPSSLLGILLDIISYVGVGFSILSLAACLVVEAVVWKSVTKNRTSYMR

HTCIVNIAASLLVANTWFIVVAAIQDNRYILCKTACVAATFFIHFFYLSVFFWMLTLGLM

LFYRLVFILHETSRSTQKAIAFCLGYGCPLAISVITLGATQPREVYTRKNVCWLNWEDTK

ALLAFAIPALIIVVVNITITIVVITKILRPSIGDKPCKQEKSSLFQISKSIGVLTPLLGL

TWGFGLTTVFPGTNLVFHIIFAILNVFQGLFILLFGCLWDLKVQEALLNKFSLSRWSSQH

SKSTSLGSSTPVFSMSSPISRRFNNLFGKTGTYNVSTPEATSSSLENSSSASSLLN

**>sp|Q9Y653|AGRG1_HUMAN Adhesion G-protein coupled receptor G1 OS=Homo sapiens OX=9606 GN=ADGRG1 PE=1 SV=2**

MTPQSLLQTTLFLLSLLFLVQGAHGRGHREDFRFCSQRNQTHRSSLHYKPTPDLRISIEN

SEEALTVHAPFPAAHPASRSFPDPRGLYHFCLYWNRHAGRLHLLYGKRDFLLSDKASSLL

CFQHQEESLAQGPPLLATSVTSWWSPQNISLPSAASFTFSFHSPPHTAAHNASVDMCELK

RDLQLLSQFLKHPQKASRRPSAAPASQQLQSLESKLTSVRFMGDMVSFEEDRINATVWKL

QPTAGLQDLHIHSRQEEEQSEIMEYSVLLPRTLFQRTKGRSGEAEKRLLLVDFSSQALFQ

DKNSSQVLGEKVLGIVVQNTKVANLTEPVVLTFQHQLQPKNVTLQCVFWVEDPTLSSPGH

WSSAGCETVRRETQTSCFCNHLTYFAVLMVSSVEVDAVHKHYLSLLSYVGCVVSALACLV

TIAAYLCSRVPLPCRRKPRDYTIKVHMNLLLAVFLLDTSFLLSEPVALTGSEAGCRASAI

FLHFSLLTCLSWMGLEGYNLYRLVVEVFGTYVPGYLLKLSAMGWGFPIFLVTLVALVDVD

NYGPIILAVHRTPEGVIYPSMCWIRDSLVSYITNLGLFSLVFLFNMAMLATMVVQILRLR

PHTQKWSHVLTLLGLSLVLGLPWALIFFSFASGTFQLVVLYLFSIITSFQGFLIFIWYWS

MRLQARGGPSPLKSNSDSARLPISSGSTSSSRI

**>sp|Q9Y653|AGRG1_HUMAN Adhesion G-protein coupled receptor G1 OS=Homo sapiens OX=9606 GN=ADGRG1 PE=1 SV=2- TL EXPOSED**

TYFAVLMVSSVEVDAVHKHYLSLLSYVGCVVSALACLV

TIAAYLCSRVPLPCRRKPRDYTIKVHMNLLLAVFLLDTSFLLSEPVALTGSEAGCRASAI

FLHFSLLTCLSWMGLEGYNLYRLVVEVFGTYVPGYLLKLSAMGWGFPIFLVTLVALVDVD

NYGPIILAVHRTPEGVIYPSMCWIRDSLVSYITNLGLFSLVFLFNMAMLATMVVQILRLR

PHTQKWSHVLTLLGLSLVLGLPWALIFFSFASGTFQLVVLYLFSIITSFQGFLIFIWYWS

MRLQARGGPSPLKSNSDSARLPISSGSTSSSRI

**>sp|Q8IZP9|AGRG2_HUMAN Adhesion G-protein coupled receptor G2 OS=Homo sapiens OX=9606 GN=ADGRG2 PE=1 SV=2**

MVFSVRQCGHVGRTEEVLLTFKIFLVIICLHVVLVTSLEEDTDNSSLSPPPAKLSVVSFA

PSSNGTPEVETTSLNDVTLSLLPSNETEKTKITIVKTFNASGVKPQRNICNLSSICNDSA

FFRGEIMFQYDKESTVPQNQHITNGTLTGVLSLSELKRSELNKTLQTLSETYFIMCATAE

AQSTLNCTFTIKLNNTMNACAVIAALERVKIRPMEHCCCSVRIPCPSSPEELEKLQCDLQ

DPIVCLADHPRGPPFSSSQSIPVVPRATVLSQVPKATSFAEPPDYSPVTHNVPSPIGEIQ

PLSPQPSAPIASSPAIDMPPQSETISSPMPQTHVSGTPPPVKASFSSPTVSAPANVNTTS

APPVQTDIVNTSSISDLENQVLQMEKALSLGSLEPNLAGEMINQVSRLLHSPPDMLAPLA

QRLLKVVDDIGLQLNFSNTTISLTSPSLALAVIRVNASSFNTTTFVAQDPANLQVSLETQ

APENSIGTITLPSSLMNNLPAHDMELASRVQFNFFETPALFQDPSLENLSLISYVISSSV

ANLTVRNLTRNVTVTLKHINPSQDELTVRCVFWDLGRNGGRGGWSDNGCSVKDRRLNETI

CTCSHLTSFGVLLDLSRTSVLPAQMMALTFITYIGCGLSSIFLSVTLVTYIAFEKIRRDY

PSKILIQLCAALLLLNLVFLLDSWIALYKMQGLCISVAVFLHYFLLVSFTWMGLEAFHMY

LALVKVFNTYIRKYILKFCIVGWGVPAVVVTIILTISPDNYGLGSYGKFPNGSPDDFCWI

NNNAVFYITVVGYFCVIFLLNVSMFIVVLVQLCRIKKKKQLGAQRKTSIQDLRSIAGLTF

LLGITWGFAFFAWGPVNVTFMYLFAIFNTLQGFFIFIFYCVAKENVRKQWRRYLCCGKLR

LAENSDWSKTATNGLKKQTVNQGVSSSSNSLQSSSNSTNSTTLLVNNDCSVHASGNGNAS

TERNGVSFSVQNGDVCLHDFTGKQHMFNEKEDSCNGKGRMALRRTSKRGSLHFIEQM

**>sp|Q8IZP9|AGRG2_HUMAN Adhesion G-protein coupled receptor G2 OS=Homo sapiens OX=9606 GN=ADGRG2 PE=1 SV=2 – TL EXPOSED**

TSFGVLLDLSRTSVLPAQMMALTFITYIGCGLSSIFLSVTLVTYIAFEKIRRDY

PSKILIQLCAALLLLNLVFLLDSWIALYKMQGLCISVAVFLHYFLLVSFTWMGLEAFHMY

LALVKVFNTYIRKYILKFCIVGWGVPAVVVTIILTISPDNYGLGSYGKFPNGSPDDFCWI

NNNAVFYITVVGYFCVIFLLNVSMFIVVLVQLCRIKKKKQLGAQRKTSIQDLRSIAGLTF

LLGITWGFAFFAWGPVNVTFMYLFAIFNTLQGFFIFIFYCVAKENVRKQWRRYLCCGKLR

LAENSDWSKTATNGLKKQTVNQGVSSSSNSLQSSSNSTNSTTLLVNNDCSVHASGNGNAS

TERNGVSFSVQNGDVCLHDFTGKQHMFNEKEDSCNGKGRMALRRTSKRGSLHFIEQM

**>sp|Q86Y34|AGRG3_HUMAN Adhesion G protein-coupled receptor G3 OS=Homo sapiens OX=9606 GN=ADGRG3 PE=1 SV=1**

MATPRGLGALLLLLLLPTSGQEKPTEGPRNTCLGSNNMYDIFNLNDKALCFTKCRQSGSD

SCNVENLQRYWLNYEAHLMKEGLTQKVNTPFLKALVQNLSTNTAEDFYFSLEPSQVPRQV

MKDEDKPPDRVRLPKSLFRSLPGNRSVVRLAVTILDIGPGTLFKGPRLGLGDGSGVLNNR

LVGLSVGQMHVTKLAEPLEIVFSHQRPPPNMTLTCVFWDVTKGTTGDWSSEGCSTEVRPE

GTVCCCDHLTFFALLLRPTLDQSTVHILTRISQAGCGVSMIFLAFTIILYAFLRLSRERF

KSEDAPKIHVALGGSLFLLNLAFLVNVGSGSKGSDAACWARGAVFHYFLLCAFTWMGLEA

FHLYLLAVRVFNTYFGHYFLKLSLVGWGLPALMVIGTGSANSYGLYTIRDRENRTSLELC

WFREGTTMYALYITVHGYFLITFLFGMVVLALVVWKIFTLSRATAVKERGKNRKKVLTLL

GLSSLVGVTWGLAIFTPLGLSTVYIFALFNSLQGVFICCWFTILYLPSQSTTVSSSTARL

DQAHSASQE

**>sp|Q86Y34|AGRG3_HUMAN Adhesion G protein-coupled receptor G3 OS=Homo sapiens OX=9606 GN=ADGRG3 PE=1 SV=1- TL EXPOSED**

TFFALLLRPTLDQSTVHILTRISQAGCGVSMIFLAFTIILYAFLRLSRERF

KSEDAPKIHVALGGSLFLLNLAFLVNVGSGSKGSDAACWARGAVFHYFLLCAFTWMGLEA

FHLYLLAVRVFNTYFGHYFLKLSLVGWGLPALMVIGTGSANSYGLYTIRDRENRTSLELC

WFREGTTMYALYITVHGYFLITFLFGMVVLALVVWKIFTLSRATAVKERGKNRKKVLTLL

GLSSLVGVTWGLAIFTPLGLSTVYIFALFNSLQGVFICCWFTILYLPSQSTTVSSSTARL

DQAHSASQE

**>sp|Q8IZF6|AGRG4_HUMAN Adhesion G-protein coupled receptor G4 OS=Homo sapiens OX=9606 GN=ADGRG4 PE=2 SV=2**

MKEHIIYQKLYGLILMSSFIFLSDTLSLKGKKLDFFGRGDTYVSLIDTIPELSRFTACID

LVFMDDNSRYWMAFSYITNNALLGREDIDLGLAGDHQQLILYRLGKTFSIRHHLASFQWH

TICLIWDGVKGKLELFLNKERILEVTDQPHNLTPHGTLFLGHFLKNESSEVKSMMRSFPG

SLYYFQLWDHILENEEFMKCLDGNIVSWEEDVWLVNKIIPTVDRTLRCFVPENMTIQEKS

TTVSQQIDMTTPSQITGVKPQNTAHSSTLLSQSIPIFATDYTTISYSNTTSPPLETMTAQ

KILKTLVDETATFAVDVLSTSSAISLPTQSISIDNTTNSMKKTKSPSSESTKTTKMVEAM

ATEIFQPPTPSNFLSTSRFTKNSVVSTTSAIKSQSAVTKTTSLFSTIESTSMSTTPCLKQ

KSTNTGALPISTAGQEFIESTAAGTVPWFTVEKTSPASTHVGTASSFPPEPVLISTAAPV

DSVFPRNQTAFPLATTDMKIAFTVHSLTLPTRLIETTPAPRTAETELTSTNFQDVSLPRV

EDAMSTSMSKETSSKTFSFLTSFSFTGTESVQTVIDAEATRTALTPEITLASTVAETMLS

STITGRVYTQNTPTADGHLLTLMSTRSASTSKAPESGPTSTTDEAAHLFSSNETIWTSRP

DQALLASMNTTTILTFVPNENFTSAFHENTTYTEYLSATTNITPLKASPEGKGTTANDAT

TARYTTAVSKLTSPWFANFSIVSGTTSITNMPEFKLTTLLLKTIPMSTKPANELPLTPRE

TVVPSVDIISTLACIQPNFSTEESASETTQTEINGAIVFGGTTTPVPKSATTQRLNATVT

RKEATSHYLMRKSTIAAVAEVSPFSTMLEVTDESAQRVTASVTVSSFPDIEKLSTPLDNK

TATTEVRESWLLTKLVKTTPRSSYNEMTEMFNFNHTYVAHWTSETSEGISAGSPTSGSTH

IFGEPLGASTTRISETSFSTTPTDRTATSLSDGILPPQPTAAHSSATPVPVTHMFSLPVN

GSSVVAEETEVTMSEPSTLARAFSTSVLSDVSNLSSTTMTTALVPPLDQTASTTIVIVPT

HGDLIRTTSEATVISVRKTSMAVPSLTETPFHSLRLSTPVTAKAETTLFSTSVDTVTPST

HTLVCSKPPPDNIPPASSTHVISTTSTPEATQPISQVEETSTYALSFPYTFSGGGVVASL

ATGTTETSVVDETTPSHISANKLTTSVNSHISSSATYRVHTPVSIQLVTSTSVLSSDKDQ

MTISLGKTPRTMEVTEMSPSKNSFISYSRGTPSLEMTDTGFPETTKISSHQTHSPSEIPL

GTPSDGNLASSPTSGSTQITPTLTSSNTVGVHIPEMSTSLGKTALPSQALTITTFLCPEK

ESTSALPAYTPRTVEMIVNSTYVTHSVSYGQDTSFVDTTTSSSTRISNPMDINTTFSHLH

SLRTQPEVTSVASFISESTQTFPESLSLSTAGLYNDGFTVLSDRITTAFSVPNVPTMLPR

ESSMATSTPIYQMSSLPVNVTAFTSKKVSDTPPIVITKSSKTMHPGCLKSPCTATSGPMS

EMSSIPVNNSAFTPATVSSDTSTRVGLFSTLLSSVTPRTTMTMQTSTLDVTPVIYAGATS

KNKMVSSAFTTEMIEAPSRITPTTFLSPTEPTLPFVKTVPTTIMAGIVTPFVGTTAFSPL

SSKSTGAISSIPKTTFSPFLSATQQSSQADEATTLGILSGITNRSLSTVNSGTGVALTDT

YSRITVPENMLSPTHADSLHTSFNIQVSPSLTSFKSASGPTKNVKTTTNCFSSNTRKMTS

LLEKTSLTNYATSLNTPVSYPPWTPSSATLPSLTSFVYSPHSTEAEISTPKTSPPPTSQM

VEFPVLGTRMTSSNTQPLLMTSWNIPTAEGSQFPISTTINVPTSNEMETETLHLVPGPLS

TFTASQTGLVSKDVMAMSSIPMSGILPNHGLSENPSLSTSLRAITSTLADVKHTFEKMTT

SVTPGTTLPSILSGATSGSVISKSPILTWLLSSLPSGSPPATVSNAPHVMTSSTVEVSKS

TFLTSDMISAHPFTNLTTLPSATMSTILTRTIPTPTLGGITTGFPTSLPMSINVTDDIVY

ISTHPEASSRTTITANPRTVSHPSSFSRKTMSPSTTDHTLSVGAMPLPSSTITSSWNRIP

TASSPSTLIIPKPTLDSLLNIMTTTSTVPGASFPLISTGVTYPFTATVSSPISSFFETTW

LDSTPSFLSTEASTSPTATKSTVSFYNVEMSFSVFVEEPRIPITSVINEFTENSLNSIFQ

NSEFSLATLETQIKSRDISEEEMVMDRAILEQREGQEMATISYVPYSCVCQVIIKASSSL

ASSELMRKIKSKIHGNFTHGNFTQDQLTLLVNCEHVAVKKLEPGNCKADETASKYKGTYK

WLLTNPTETAQTRCIKNEDGNATRFCSISINTGKSQWEKPKFKQCKLLQELPDKIVDLAN

ITISDENAEDVAEHILNLINESPALGKEETKIIVSKISDISQCDEISMNLTHVMLQIINV

VLEKQNNSASDLHEISNEILRIIERTGHKMEFSGQIANLTVAGLALAVLRGDHTFDGMAF

SIHSYEEGTDPEIFLGNVPVGGILASIYLPKSLTERIPLSNLQTILFNFFGQTSLFKTKN

VTKALTTYVVSASISDDMFIQNLADPVVITLQHIGGNQNYGQVHCAFWDFENNNGLGGWN

SSGCKVKETNVNYTICQCDHLTHFGVLMDLSRSTVDSVNEQILALITYTGCGISSIFLGV

AVVTYIAFHKLRKDYPAKILINLCTALLMLNLVFLINSWLSSFQKVGVCITAAVALHYFL

LVSFTWMGLEAVHMYLALVKVFNIYIPNYILKFCLVGWGIPAIMVAITVSVKKDLYGTLS

PTTPFCWIKDDSIFYISVVAYFCLIFLMNLSMFCTVLVQLNSVKSQIQKTRRKMILHDLK

GTMSLTFLLGLTWGFAFFAWGPMRNFFLYLFAIFNTLQGFFIFVFHCVMKESVREQWQIH

LCCGWLRLDNSSDGSSRCQIKVGYKQEGLKKIFEHKLLTPSLKSTATSSTFKSLGSAQGT

PSEISFPNDDFDKDPYCSSP

**>sp|Q8IZF6|AGRG4_HUMAN Adhesion G-protein coupled receptor G4 OS=Homo sapiens OX=9606 GN=ADGRG4 PE=2 SV=2 – TL EXPOSED**

THFGVLMDLSRSTVDSVNEQILALITYTGCGISSIFLGV

AVVTYIAFHKLRKDYPAKILINLCTALLMLNLVFLINSWLSSFQKVGVCITAAVALHYFL

LVSFTWMGLEAVHMYLALVKVFNIYIPNYILKFCLVGWGIPAIMVAITVSVKKDLYGTLS

PTTPFCWIKDDSIFYISVVAYFCLIFLMNLSMFCTVLVQLNSVKSQIQKTRRKMILHDLK

GTMSLTFLLGLTWGFAFFAWGPMRNFFLYLFAIFNTLQGFFIFVFHCVMKESVREQWQIH

LCCGWLRLDNSSDGSSRCQIKVGYKQEGLKKIFEHKLLTPSLKSTATSSTFKSLGSAQGT

PSEISFPNDDFDKDPYCSSP

**>sp|Q8IZF4|AGRG5_HUMAN Adhesion G-protein coupled receptor G5 OS=Homo sapiens OX=9606 GN=ADGRG5 PE=2 SV=3**

MDHCGALFLCLCLLTLQNATTETWEELLSYMENMQVSRGRSSVFSSRQLHQLEQMLLNTS

FPGYNLTLQTPTIQSLAFKLSCDFSGLSLTSATLKRVPQAGGQHARGQHAMQFPAELTRD

ACKTRPRELRLICIYFSNTHFFKDENNSSLLNNYVLGAQLSHGHVNNLRDPVNISFWHNQ

SLEGYTLTCVFWKEGARKQPWGGWSPEGCRTEQPSHSQVLCRCNHLTYFAVLMQLSPALV

PAELLAPLTYISLVGCSISIVASLITVLLHFHFRKQSDSLTRIHMNLHASVLLLNIAFLL

SPAFAMSPVPGSACTALAAALHYALLSCLTWMAIEGFNLYLLLGRVYNIYIRRYVFKLGV

LGWGAPALLVLLSLSVKSSVYGPCTIPVFDSWENGTGFQNMSICWVRSPVVHSVLVMGYG

GLTSLFNLVVLAWALWTLRRLRERADAPSVRACHDTVTVLGLTVLLGTTWALAFFSFGVF

LLPQLFLFTILNSLYGFFLFLWFCSQRCRSEAEAKAQIEAFSSSQTTQ

**>sp|Q8IZF4|AGRG5_HUMAN Adhesion G-protein coupled receptor G5 OS=Homo sapiens OX=9606 GN=ADGRG5 PE=2 SV=3- TL EXPOSED**

TYFAVLMQLSPALVPAELLAPLTYISLVGCSISIVASLITVLLHFHFRKQSDSLTRIHMN

LHASVLLLNIAFLLSPAFAMSPVPGSACTALAAALHYALLSCLTWMAIEGFNLYLLLGRV

YNIYIRRYVFKLGVLGWGAPALLVLLSLSVKSSVYGPCTIPVFDSWENGTGFQNMSICWV

RSPVVHSVLVMGYGGLTSLFNLVVLAWALWTLRRLRERADAPSVRACHDTVTVLGLTVLL

GTTWALAFFSFGVFLLPQLFLFTILNSLYGFFLFLWFCSQRCRSEAEAKAQIEAFSSSQT

TQ

**>sp|Q86SQ4|AGRG6_HUMAN Adhesion G-protein coupled receptor G6 OS=Homo sapiens OX=9606 GN=ADGRG6 PE=1 SV=3**

MMFRSDRMWSCHWKWKPSPLLFLFALYIMCVPHSVWGCANCRVVLSNPSGTFTSPCYPND

YPNSQACMWTLRAPTGYIIQITFNDFDIEEAPNCIYDSLSLDNGESQTKFCGATAKGLSF

NSSANEMHVSFSSDFSIQKKGFNASYIRVAVSLRNQKVILPQTSDAYQVSVAKSISIPEL

SAFTLCFEATKVGHEDSDWTAFSYSNASFTQLLSFGKAKSGYFLSISDSKCLLNNALPVK

EKEDIFAESFEQLCLVWNNSLGSIGVNFKRNYETVPCDSTISKVIPGNGKLLLGSNQNEI

VSLKGDIYNFRLWNFTMNAKILSNLSCNVKGNVVDWQNDFWNIPNLALKAESNLSCGSYL

IPLPAAELASCADLGTLCQATVNSPSTTPPTVTTNMPVTNRIDKQRNDGIIYRISVVIQN

ILRHPEVKVQSKVAEWLNSTFQNWNYTVYVVNISFHLSAGEDKIKVKRSLEDEPRLVLWA

LLVYNATNNTNLEGKIIQQKLLKNNESLDEGLRLHTVNVRQLGHCLAMEEPKGYYWPSIQ

PSEYVLPCPDKPGFSASRICFYNATNPLVTYWGPVDISNCLKEANEVANQILNLTADGQN

LTSANITNIVEQVKRIVNKEENIDITLGSTLMNIFSNILSSSDSDLLESSSEALKTIDEL

AFKIDLNSTSHVNITTRNLALSVSSLLPGTNAISNFSIGLPSNNESYFQMDFESGQVDPL

ASVILPPNLLENLSPEDSVLVRRAQFTFFNKTGLFQDVGPQRKTLVSYVMACSIGNITIQ

NLKDPVQIKIKHTRTQEVHHPICAFWDLNKNKSFGGWNTSGCVAHRDSDASETVCLCNHF

THFGVLMDLPRSASQLDARNTKVLTFISYIGCGISAIFSAATLLTYVAFEKLRRDYPSKI

LMNLSTALLFLNLLFLLDGWITSFNVDGLCIAVAVLLHFFLLATFTWMGLEAIHMYIALV

KVFNTYIRRYILKFCIIGWGLPALVVSVVLASRNNNEVYGKESYGKEKGDEFCWIQDPVI

FYVTCAGYFGVMFFLNIAMFIVVMVQICGRNGKRSNRTLREEVLRNLRSVVSLTFLLGMT

WGFAFFAWGPLNIPFMYLFSIFNSLQGLFIFIFHCAMKENVQKQWRQHLCCGRFRLADNS

DWSKTATNIIKKSSDNLGKSLSSSSIGSNSTYLTSKSKSSSTTYFKRNSHTDNVSYEHSF

NKSGSLRQCFHGQVLVKTGPC

**>sp|Q86SQ4|AGRG6_HUMAN Adhesion G-protein coupled receptor G6 OS=Homo sapiens OX=9606 GN=ADGRG6 PE=1 SV=3- TL EXPOSED**

THFGVLMDLPRSASQLDARNTKVLTFISYIGCGISAIFSAATLLTYVAFEKLRRDYPSKI

LMNLSTALLFLNLLFLLDGWITSFNVDGLCIAVAVLLHFFLLATFTWMGLEAIHMYIALV

KVFNTYIRRYILKFCIIGWGLPALVVSVVLASRNNNEVYGKESYGKEKGDEFCWIQDPVI

FYVTCAGYFGVMFFLNIAMFIVVMVQICGRNGKRSNRTLREEVLRNLRSVVSLTFLLGMT

WGFAFFAWGPLNIPFMYLFSIFNSLQGLFIFIFHCAMKENVQKQWRQHLCCGRFRLADNS

DWSKTATNIIKKSSDNLGKSLSSSSIGSNSTYLTSKSKSSSTTYFKRNSHTDNVSYEHSF

NKSGSLRQCFHGQVLVKTGPC

**>sp|Q96K78|AGRG7_HUMAN Adhesion G-protein coupled receptor G7 OS=Homo sapiens OX=9606 GN=ADGRG7 PE=1 SV=2**

MASCRAWNLRVLVAVVCGLLTGIILGLGIWRIVIRIQRGKSTSSSSTPTEFCRNGGTWEN

GRCICTEEWKGLRCTIANFCENSTYMGFTFARIPVGRYGPSLQTCGKDTPNAGNPMAVRL

CSLSLYGEIELQKVTIGNCNENLETLEKQVKDVTAPLNNISSEVQILTSDANKLTAENIT

SATRVVGQIFNTSRNASPEAKKVAIVTVSQLLDASEDAFQRVAATANDDALTTLIEQMET

YSLSLGNQSVVEPNIAIQSANFSSENAVGPSNVRFSVQKGASSSLVSSSTFIHTNVDGLN

PDAQTELQVLLNMTKNYTKTCGFVVYQNDKLFQSKTFTAKSDFSQKIISSKTDENEQDQS

ASVDMVFSPKYNQKEFQLYSYACVYWNLSAKDWDTYGCQKDKGTDGFLRCRCNHTTNFAV

LMTFKKDYQYPKSLDILSNVGCALSVTGLALTVIFQIVTRKVRKTSVTWVLVNLCISMLI

FNLLFVFGIENSNKNLQTSDGDINNIDFDNNDIPRTDTINIPNPMCTAIAALLHYFLLVT

FTWNALSAAQLYYLLIRTMKPLPRHFILFISLIGWGVPAIVVAITVGVIYSQNGNNPQWE

LDYRQEKICWLAIPEPNGVIKSPLLWSFIVPVTIILISNVVMFITISIKVLWKNNQNLTS

TKKVSSMKKIVSTLSVAVVFGITWILAYLMLVNDDSIRIVFSYIFCLFNTTQGLQIFILY

TVRTKVFQSEASKVLMLLSSIGRRKSLPSVTRPRLRVKMYNFLRSLPTLHERFRLLETSP

STEEITLSESDNAKESI

**>sp|Q96K78|AGRG7_HUMAN Adhesion G-protein coupled receptor G7 OS=Homo sapiens OX=9606 GN=ADGRG7 PE=1 SV=2 – TL EXPOSED**

TNFAVLMTFKKDYQYPKSLDILSNVGCALSVTGLALTVIFQIVTRKVRKTSVTWVLVNLCISMLI

FNLLFVFGIENSNKNLQTSDGDINNIDFDNNDIPRTDTINIPNPMCTAIAALLHYFLLVT

FTWNALSAAQLYYLLIRTMKPLPRHFILFISLIGWGVPAIVVAITVGVIYSQNGNNPQWE

LDYRQEKICWLAIPEPNGVIKSPLLWSFIVPVTIILISNVVMFITISIKVLWKNNQNLTS

TKKVSSMKKIVSTLSVAVVFGITWILAYLMLVNDDSIRIVFSYIFCLFNTTQGLQIFILY

TVRTKVFQSEASKVLMLLSSIGRRKSLPSVTRPRLRVKMYNFLRSLPTLHERFRLLETSP

STEEITLSESDNAKESI

**>sp|O94910|AGRL1_HUMAN Adhesion G protein-coupled receptor L1 OS=Homo sapiens OX=9606 GN=ADGRL1 PE=1 SV=1**

MARLAAVLWNLCVTAVLVTSATQGLSRAGLPFGLMRRELACEGYPIELRCPGSDVIMVEN

ANYGRTDDKICDADPFQMENVQCYLPDAFKIMSQRCNNRTQCVVVAGSDAFPDPCPGTYK

YLEVQYDCVPYKVEQKVFVCPGTLQKVLEPTSTHESEHQSGAWCKDPLQAGDRIYVMPWI

PYRTDTLTEYASWEDYVAARHTTTYRLPNRVDGTGFVVYDGAVFYNKERTRNIVKYDLRT

RIKSGETVINTANYHDTSPYRWGGKTDIDLAVDENGLWVIYATEGNNGRLVVSQLNPYTL

RFEGTWETGYDKRSASNAFMVCGVLYVLRSVYVDDDSEAAGNRVDYAFNTNANREEPVSL

TFPNPYQFISSVDYNPRDNQLYVWNNYFVVRYSLEFGPPDPSAGPATSPPLSTTTTARPT

PLTSTASPAATTPLRRAPLTTHPVGAINQLGPDLPPATAPVPSTRRPPAPNLHVSPELFC

EPREVRRVQWPATQQGMLVERPCPKGTRGIASFQCLPALGLWNPRGPDLSNCTSPWVNQV

AQKIKSGENAANIASELARHTRGSIYAGDVSSSVKLMEQLLDILDAQLQALRPIERESAG

KNYNKMHKRERTCKDYIKAVVETVDNLLRPEALESWKDMNATEQVHTATMLLDVLEEGAF

LLADNVREPARFLAAKENVVLEVTVLNTEGQVQELVFPQEEYPRKNSIQLSAKTIKQNSR

NGVVKVVFILYNNLGLFLSTENATVKLAGEAGPGGPGGASLVVNSQVIAASINKESSRVF

LMDPVIFTVAHLEDKNHFNANCSFWNYSERSMLGYWSTQGCRLVESNKTHTTCACSHLTN

FAVLMAHREIYQGRINELLLSVITWVGIVISLVCLAICISTFCFLRGLQTDRNTIHKNLC

INLFLAELLFLVGIDKTQYEIACPIFAGLLHYFFLAAFSWLCLEGVHLYLLLVEVFESEY

SRTKYYYLGGYCFPALVVGIAAAIDYRSYGTEKACWLRVDNYFIWSFIGPVSFVIVVNLV

FLMVTLHKMIRSSSVLKPDSSRLDNIKSWALGAIALLFLLGLTWAFGLLFINKESVVMAY

LFTTFNAFQGVFIFVFHCALQKKVHKEYSKCLRHSYCCIRSPPGGTHGSLKTSAMRSNTR

YYTGTQSRIRRMWNDTVRKQTESSFMAGDINSTPTLNRGTMGNHLLTNPVLQPRGGTSPY

NTLIAESVGFNPSSPPVFNSPGSYREPKHPLGGREACGMDTLPLNGNFNNSYSLRSGDFP

PGDGGPEPPRGRNLADAAAFEKMIISELVHNNLRGSSSAAKGPPPPEPPVPPVPGGGGEE

EAGGPGGADRAEIELLYKALEEPLLLPRAQSVLYQSDLDESESCTAEDGATSRPLSSPPG

RDSLYASGANLRDSPSYPDSSPEGPSEALPPPPPAPPGPPEIYYTSRPPALVARNPLQGY

YQVRRPSHEGYLAAPGLEGPGPDGDGQMQLVTSL

**>sp|O94910|AGRL1_HUMAN Adhesion G protein-coupled receptor L1 OS=Homo sapiens OX=9606 GN=ADGRL1 PE=1 SV=1**

TNFAVLMAHREIYQGRINELLLSVITWVGIVISLVCLAICISTFCFLRGLQTDRNTIHKNLC

INLFLAELLFLVGIDKTQYEIACPIFAGLLHYFFLAAFSWLCLEGVHLYLLLVEVFESEY

SRTKYYYLGGYCFPALVVGIAAAIDYRSYGTEKACWLRVDNYFIWSFIGPVSFVIVVNLV

FLMVTLHKMIRSSSVLKPDSSRLDNIKSWALGAIALLFLLGLTWAFGLLFINKESVVMAY

LFTTFNAFQGVFIFVFHCALQKKVHKEYSKCLRHSYCCIRSPPGGTHGSLKTSAMRSNTR

YYTGTQSRIRRMWNDTVRKQTESSFMAGDINSTPTLNRGTMGNHLLTNPVLQPRGGTSPY

NTLIAESVGFNPSSPPVFNSPGSYREPKHPLGGREACGMDTLPLNGNFNNSYSLRSGDFP

PGDGGPEPPRGRNLADAAAFEKMIISELVHNNLRGSSSAAKGPPPPEPPVPPVPGGGGEE

EAGGPGGADRAEIELLYKALEEPLLLPRAQSVLYQSDLDESESCTAEDGATSRPLSSPPG

RDSLYASGANLRDSPSYPDSSPEGPSEALPPPPPAPPGPPEIYYTSRPPALVARNPLQGY

YQVRRPSHEGYLAAPGLEGPGPDGDGQMQLVTSL

**>sp|O95490|AGRL2_HUMAN Adhesion G protein-coupled receptor L2 OS=Homo sapiens OX=9606 GN=ADGRL2 PE=1 SV=2**

MVSSGCRMRSLWFIIVISFLPNTEGFSRAALPFGLVRRELSCEGYSIDLRCPGSDVIMIE

SANYGRTDDKICDADPFQMENTDCYLPDAFKIMTQRCNNRTQCIVVTGSDVFPDPCPGTY

KYLEVQYECVPYIFVCPGTLKAIVDSPCIYEAEQKAGAWCKDPLQAADKIYFMPWTPYRT

DTLIEYASLEDFQNSRQTTTYKLPNRVDGTGFVVYDGAVFFNKERTRNIVKFDLRTRIKS

GEAIINYANYHDTSPYRWGGKTDIDLAVDENGLWVIYATEQNNGMIVISQLNPYTLRFEA

TWETVYDKRAASNAFMICGVLYVVRSVYQDNESETGKNSIDYIYNTRLNRGEYVDVPFPN

QYQYIAAVDYNPRDNQLYVWNNNFILRYSLEFGPPDPAQVPTTAVTITSSAELFKTIIST

TSTTSQKGPMSTTVAGSQEGSKGTKPPPAVSTTKIPPITNIFPLPERFCEALDSKGIKWP

QTQRGMMVERPCPKGTRGTASYLCMISTGTWNPKGPDLSNCTSHWVNQLAQKIRSGENAA

SLANELAKHTKGPVFAGDVSSSVRLMEQLVDILDAQLQELKPSEKDSAGRSYNKLQKREK

TCRAYLKAIVDTVDNLLRPEALESWKHMNSSEQAHTATMLLDTLEEGAFVLADNLLEPTR

VSMPTENIVLEVAVLSTEGQIQDFKFPLGIKGAGSSIQLSANTVKQNSRNGLAKLVFIIY

RSLGQFLSTENATIKLGADFIGRNSTIAVNSHVISVSINKESSRVYLTDPVLFTLPHIDP

DNYFNANCSFWNYSERTMMGYWSTQGCKLVDTNKTRTTCACSHLTNFAILMAHREIAYKD

GVHELLLTVITWVGIVISLVCLAICIFTFCFFRGLQSDRNTIHKNLCINLFIAEFIFLIG

IDKTKYAIACPIFAGLLHFFFLAAFAWMCLEGVQLYLMLVEVFESEYSRKKYYYVAGYLF

PATVVGVSAAIDYKSYGTEKACWLHVDNYFIWSFIGPVTFIILLNIIFLVITLCKMVKHS

NTLKPDSSRLENIKSWVLGAFALLCLLGLTWSFGLLFINEETIVMAYLFTIFNAFQGVFI

FIFHCALQKKVRKEYGKCFRHSYCCGGLPTESPHSSVKASTTRTSARYSSGTQSRIRRMW

NDTVRKQSESSFISGDINSTSTLNQGMTGNYLLTNPLLRPHGTNNPYNTLLAETVVCNAP

SAPVFNSPGHSLNNARDTSAMDTLPLNGNFNNSYSLHKGDYNDSVQVVDCGLSLNDTAFE

KMIISELVHNNLRGSSKTHNLELTLPVKPVIGGSSSEDDAIVADASSLMHSDNPGLELHH

KELEAPLIPQRTHSLLYQPQKKVKSEGTDSYVSQLTAEAEDHLQSPNRDSLYTSMPNLRD

SPYPESSPDMEEDLSPSRRSENEDIYYKSMPNLGAGHQLQMCYQISRGNSDGYIIPINKE

GCIPEGDVREGQMQLVTSL

**>sp|O95490|AGRL2_HUMAN Adhesion G protein-coupled receptor L2 OS=Homo sapiens OX=9606 GN=ADGRL2 PE=1 SV=2 – TL EXPOSED**

TNFAILMAHREIAYKDGVHELLLTVITWVGIVISLVCLAICIFTFCFFRGLQSDRNTIHK

NLCINLFIAEFIFLIGIDKTKYAIACPIFAGLLHFFFLAAFAWMCLEGVQLYLMLVEVFE

SEYSRKKYYYVAGYLFPATVVGVSAAIDYKSYGTEKACWLHVDNYFIWSFIGPVTFIILL

NIIFLVITLCKMVKHSNTLKPDSSRLENIKSWVLGAFALLCLLGLTWSFGLLFINEETIV

MAYLFTIFNAFQGVFIFIFHCALQKKVRKEYGKCFRHSYCCGGLPTESPHSSVKASTTRT

SARYSSGTQSRIRRMWNDTVRKQSESSFISGDINSTSTLNQGMTGNYLLTNPLLRPHGTN

NPYNTLLAETVVCNAPSAPVFNSPGHSLNNARDTSAMDTLPLNGNFNNSYSLHKGDYNDS

VQVVDCGLSLNDTAFEKMIISELVHNNLRGSSKTHNLELTLPVKPVIGGSSSEDDAIVAD

ASSLMHSDNPGLELHHKELEAPLIPQRTHSLLYQPQKKVKSEGTDSYVSQLTAEAEDHLQ

SPNRDSLYTSMPNLRDSPYPESSPDMEEDLSPSRRSENEDIYYKSMPNLGAGHQLQMCYQ

ISRGNSDGYIIPINKEGCIPEGDVREGQMQLVTSL

**>sp|Q9HAR2|AGRL3_HUMAN Adhesion G protein-coupled receptor L3 OS=Homo sapiens OX=9606 GN=ADGRL3 PE=1 SV=2**

MWPSQLLIFMMLLAPIIHAFSRAPIPMAVVRRELSCESYPIELRCPGTDVIMIESANYGR

TDDKICDSDPAQMENIRCYLPDAYKIMSQRCNNRTQCAVVAGPDVFPDPCPGTYKYLEVQ

YECVPYKVEQKVFLCPGLLKGVYQSEHLFESDHQSGAWCKDPLQASDKIYYMPWTPYRTD

TLTEYSSKDDFIAGRPTTTYKLPHRVDGTGFVVYDGALFFNKERTRNIVKFDLRTRIKSG

EAIIANANYHDTSPYRWGGKSDIDLAVDENGLWVIYATEQNNGKIVISQLNPYTLRIEGT

WDTAYDKRSASNAFMICGILYVVKSVYEDDDNEATGNKIDYIYNTDQSKDSLVDVPFPNS

YQYIAAVDYNPRDNLLYVWNNYHVVKYSLDFGPLDSRSGQAHHGQVSYISPPIHLDSELE

RPSVKDISTTGPLGMGSTTTSTTLRTTTLSPGRSTTPSVSGRRNRSTSTPSPAVEVLDDM

TTHLPSASSQIPALEESCEAVEAREIMWFKTRQGQIAKQPCPAGTIGVSTYLCLAPDGIW

DPQGPDLSNCSSPWVNHITQKLKSGETAANIARELAEQTRNHLNAGDITYSVRAMDQLVG

LLDVQLRNLTPGGKDSAARSLNKAMVETVNNLLQPQALNAWRDLTTSDQLRAATMLLHTV

EESAFVLADNLLKTDIVRENTDNIKLEVARLSTEGNLEDLKFPENMGHGSTIQLSANTLK

QNGRNGEIRVAFVLYNNLGPYLSTENASMKLGTEALSTNHSVIVNSPVITAAINKEFSNK

VYLADPVVFTVKHIKQSEENFNPNCSFWSYSKRTMTGYWSTQGCRLLTTNKTHTTCSCNH

LTNFAVLMAHVEVKHSDAVHDLLLDVITWVGILLSLVCLLICIFTFCFFRGLQSDRNTIH

KNLCISLFVAELLFLIGINRTDQPIACAVFAALLHFFFLAAFTWMFLEGVQLYIMLVEVF

ESEHSRRKYFYLVGYGMPALIVAVSAAVDYRSYGTDKVCWLRLDTYFIWSFIGPATLIIM

LNVIFLGIALYKMFHHTAILKPESGCLDNIKSWVIGAIALLCLLGLTWAFGLMYINESTV

IMAYLFTIFNSLQGMFIFIFHCVLQKKVRKEYGKCLRTHCCSGKSTESSIGSGKTSGSRT

PGRYSTGSQSRIRRMWNDTVRKQSESSFITGDINSSASLNREGLLNNARDTSVMDTLPLN

GNHGNSYSIASGEYLSNCVQIIDRGYNHNETALEKKILKELTSNYIPSYLNNHERSSEQN

RNLMNKLVNNLGSGREDDAIVLDDATSFNHEESLGLELIHEESDAPLLPPRVYSTENHQP

HHYTRRRIPQDHSESFFPLLTNEHTEDLQSPHRDSLYTSMPTLAGVAATESVTTSTQTEP

PPAKCGDAEDVYYKSMPNLGSRNHVHQLHTYYQLGRGSSDGFIVPPNKDGTPPEGSSKGP

AHLVTSL

**>sp|Q9HAR2|AGRL3_HUMAN Adhesion G protein-coupled receptor L3 OS=Homo sapiens OX=9606 GN=ADGRL3 PE=1 SV=2 – TL EXPOSED**

TNFAVLMAHVEVKHSDAVHDLLLDVITWVGILLSLVCLLICIFTFCFFRGLQSDRNTIH

KNLCISLFVAELLFLIGINRTDQPIACAVFAALLHFFFLAAFTWMFLEGVQLYIMLVEVF

ESEHSRRKYFYLVGYGMPALIVAVSAAVDYRSYGTDKVCWLRLDTYFIWSFIGPATLIIM

LNVIFLGIALYKMFHHTAILKPESGCLDNIKSWVIGAIALLCLLGLTWAFGLMYINESTV

IMAYLFTIFNSLQGMFIFIFHCVLQKKVRKEYGKCLRTHCCSGKSTESSIGSGKTSGSRT

PGRYSTGSQSRIRRMWNDTVRKQSESSFITGDINSSASLNREGLLNNARDTSVMDTLPLN

GNHGNSYSIASGEYLSNCVQIIDRGYNHNETALEKKILKELTSNYIPSYLNNHERSSEQN

RNLMNKLVNNLGSGREDDAIVLDDATSFNHEESLGLELIHEESDAPLLPPRVYSTENHQP

HHYTRRRIPQDHSESFFPLLTNEHTEDLQSPHRDSLYTSMPTLAGVAATESVTTSTQTEP

PPAKCGDAEDVYYKSMPNLGSRNHVHQLHTYYQLGRGSSDGFIVPPNKDGTPPEGSSKGP

AHLVTSL

**>sp|Q9HBW9|AGRL4_HUMAN Adhesion G protein-coupled receptor L4 OS=Homo sapiens OX=9606 GN=ADGRL4 PE=1 SV=3**

MKRLPLLVVFSTLLNCSYTQNCTKTPCLPNAKCEIRNGIEACYCNMGFSGNGVTICEDDN

ECGNLTQSCGENANCTNTEGSYYCMCVPGFRSSSNQDRFITNDGTVCIENVNANCHLDNV

CIAANINKTLTKIRSIKEPVALLQEVYRNSVTDLSPTDIITYIEILAESSSLLGYKNNTI

SAKDTLSNSTLTEFVKTVNNFVQRDTFVVWDKLSVNHRRTHLTKLMHTVEQATLRISQSF

QKTTEFDTNSTDIALKVFFFDSYNMKHIHPHMNMDGDYINIFPKRKAAYDSNGNVAVAFV

YYKSIGPLLSSSDNFLLKPQNYDNSEEEERVISSVISVSMSSNPPTLYELEKITFTLSHR

KVTDRYRSLCAFWNYSPDTMNGSWSSEGCELTYSNETHTSCRCNHLTHFAILMSSGPSIG

IKDYNILTRITQLGIIISLICLAICIFTFWFFSEIQSTRTTIHKNLCCSLFLAELVFLVG

INTNTNKLFCSIIAGLLHYFFLAAFAWMCIEGIHLYLIVVGVIYNKGFLHKNFYIFGYLS

PAVVVGFSAALGYRYYGTTKVCWLSTENNFIWSFIGPACLIILVNLLAFGVIIYKVFRHT

AGLKPEVSCFENIRSCARGALALLFLLGTTWIFGVLHVVHASVVTAYLFTVSNAFQGMFI

FLFLCVLSRKIQEEYYRLFKNVPCCFGCLR

**>sp|Q9HBW9|AGRL4_HUMAN Adhesion G protein-coupled receptor L4 OS=Homo sapiens OX=9606 GN=ADGRL4 PE=1 SV=3** **– TL EXPOSED**

THFAILMSSGPSIGIKDYNILTRITQLGIIISLICLAICIFTFWFFSEIQSTRTTIHKNL

CCSLFLAELVFLVGINTNTNKLFCSIIAGLLHYFFLAAFAWMCIEGIHLYLIVVGVIYNK

GFLHKNFYIFGYLSPAVVVGFSAALGYRYYGTTKVCWLSTENNFIWSFIGPACLIILVNL

LAFGVIIYKVFRHTAGLKPEVSCFENIRSCARGALALLFLLGTTWIFGVLHVVHASVVTA

YLFTVSNAFQGMFIFLFLCVLSRKIQEEYYRLFKNVPCCFGCLR

**>sp|Q9NYQ6|CELR1_HUMAN Cadherin EGF LAG seven-pass G-type receptor 1 OS=Homo sapiens OX=9606 GN=CELSR1 PE=1 SV=1, ADGRC1**

MAPPPPPVLPVLLLLAAAAALPAMGLRAAAWEPRVPGGTRAFALRPGCTYAVGAACTPRA

PRELLDVGRDGRLAGRRRVSGAGRPLPLQVRLVARSAPTALSRRLRARTHLPGCGARARL

CGTGARLCGALCFPVPGGCAAAQHSALAAPTTLPACRCPPRPRPRCPGRPICLPPGGSVR

LRLLCALRRAAGAVRVGLALEAATAGTPSASPSPSPPLPPNLPEARAGPARRARRGTSGR

GSLKFPMPNYQVALFENEPAGTLILQLHAHYTIEGEEERVSYYMEGLFDERSRGYFRIDS

ATGAVSTDSVLDRETKETHVLRVKAVDYSTPPRSATTYITVLVKDTNDHSPVFEQSEYRE

RVRENLEVGYEVLTIRASDRDSPINANLRYRVLGGAWDVFQLNESSGVVSTRAVLDREEA

AEYQLLVEANDQGRNPGPLSATATVYIEVEDENDNYPQFSEQNYVVQVPEDVGLNTAVLR

VQATDRDQGQNAAIHYSILSGNVAGQFYLHSLSGILDVINPLDFEDVQKYSLSIKAQDGG

RPPLINSSGVVSVQVLDVNDNEPIFVSSPFQATVLENVPLGYPVVHIQAVDADSGENARL

HYRLVDTASTFLGGGSAGPKNPAPTPDFPFQIHNSSGWITVCAELDREEVEHYSFGVEAV

DHGSPPMSSSTSVSITVLDVNDNDPVFTQPTYELRLNEDAAVGSSVLTLQARDRDANSVI

TYQLTGGNTRNRFALSSQRGGGLITLALPLDYKQEQQYVLAVTASDGTRSHTAHVLINVT

DANTHRPVFQSSHYTVSVSEDRPVGTSIATLSANDEDTGENARITYVIQDPVPQFRIDPD

SGTMYTMMELDYENQVAYTLTIMAQDNGIPQKSDTTTLEILILDANDNAPQFLWDFYQGS

IFEDAPPSTSILQVSATDRDSGPNGRLLYTFQGGDDGDGDFYIEPTSGVIRTQRRLDREN

VAVYNLWALAVDRGSPTPLSASVEIQVTILDINDNAPMFEKDELELFVEENNPVGSVVAK

IRANDPDEGPNAQIMYQIVEGDMRHFFQLDLLNGDLRAMVELDFEVRREYVLVVQATSAP

LVSRATVHILLVDQNDNPPVLPDFQILFNNYVTNKSNSFPTGVIGCIPAHDPDVSDSLNY

TFVQGNELRLLLLDPATGELQLSRDLDNNRPLEALMEVSVSDGIHSVTAFCTLRVTIITD

DMLTNSITVRLENMSQEKFLSPLLALFVEGVAAVLSTTKDDVFVFNVQNDTDVSSNILNV

TFSALLPGGVRGQFFPSEDLQEQIYLNRTLLTTISTQRVLPFDDNICLREPCENYMKCVS

VLRFDSSAPFLSSTTVLFRPIHPINGLRCRCPPGFTGDYCETEIDLCYSDPCGANGRCRS

REGGYTCECFEDFTGEHCEVDARSGRCANGVCKNGGTCVNLLIGGFHCVCPPGEYERPYC

EVTTRSFPPQSFVTFRGLRQRFHFTISLTFATQERNGLLLYNGRFNEKHDFIALEIVDEQ

VQLTFSAGETTTTVAPKVPSGVSDGRWHSVQVQYYNKPNIGHLGLPHGPSGEKMAVVTVD

DCDTTMAVRFGKDIGNYSCAAQGTQTGSKKSLDLTGPLLLGGVPNLPEDFPVHNRQFVGC

MRNLSVDGKNVDMAGFIANNGTREGCAARRNFCDGRRCQNGGTCVNRWNMYLCECPLRFG

GKNCEQAMPHPQLFSGESVVSWSDLNIIISVPWYLGLMFRTRKEDSVLMEATSGGPTSFR

LQILNNYLQFEVSHGPSDVESVMLSGLRVTDGEWHHLLIELKNVKEDSEMKHLVTMTLDY

GMDQNKADIGGMLPGLTVRSVVVGGASEDKVSVRRGFRGCMQGVRMGGTPTNVATLNMNN

ALKVRVKDGCDVDDPCTSSPCPPNSRCHDAWEDYSCVCDKGYLGINCVDACHLNPCENMG

ACVRSPGSPQGYVCECGPSHYGPYCENKLDLPCPRGWWGNPVCGPCHCAVSKGFDPDCNK

TNGQCQCKENYYKLLAQDTCLPCDCFPHGSHSRTCDMATGQCACKPGVIGRQCNRCDNPF

AEVTTLGCEVIYNGCPKAFEAGIWWPQTKFGQPAAVPCPKGSVGNAVRHCSGEKGWLPPE

LFNCTTISFVDLRAMNEKLSRNETQVDGARALQLVRALRSATQHTGTLFGNDVRTAYQLL

GHVLQHESWQQGFDLAATQDADFHEDVIHSGSALLAPATRAAWEQIQRSEGGTAQLLRRL

EGYFSNVARNVRRTYLRPFVIVTANMILAVDIFDKFNFTGARVPRFDTIHEEFPRELESS

VSFPADFFRPPEEKEGPLLRPAGRRTTPQTTRPGPGTEREAPISRRRRHPDDAGQFAVAL

VIIYRTLGQLLPERYDPDRRSLRLPHRPIINTPMVSTLVYSEGAPLPRPLERPVLVEFAL

LEVEERTKPVCVFWNHSLAVGGTGGWSARGCELLSRNRTHVACQCSHTASFAVLMDISRR

ENGEVLPLKIVTYAAVSLSLAALLVAFVLLSLVRMLRSNLHSIHKHLAVALFLSQLVFVI

GINQTENPFLCTVVAILLHYIYMSTFAWTLVESLHVYRMLTEVRNIDTGPMRFYYVVGWG

IPAIVTGLAVGLDPQGYGNPDFCWLSLQDTLIWSFAGPIGAVIIINTVTSVLSAKVSCQR

KHHYYGKKGIVSLLRTAFLLLLLISATWLLGLLAVNRDALSFHYLFAIFSGLQGPFVLLF

HCVLNQEVRKHLKGVLGGRKLHLEDSATTRATLLTRSLNCNTTFGDGPDMLRTDLGESTA

SLDSIVRDEGIQKLGVSSGLVRGSHGEPDASLMPRSCKDPPGHDSDSDSELSLDEQSSSY

ASSHSSDSEDDGVGAEEKWDPARGAVHSTPKGDAVANHVPAGWPDQSLAESDSEDPSGKP

RLKVETKVSVELHREEQGSHRGEYPPDQESGGAARLASSQPPEQRKGILKNKVTYPPPLT

LTEQTLKGRLREKLADCEQSPTSSRTSSLGSGGPDCAITVKSPGREPGRDHLNGVAMNVRTGSAQADGSDSEKP

**>sp|Q9NYQ6|CELR1_HUMAN Cadherin EGF LAG seven-pass G-type receptor 1 OS=Homo sapiens OX=9606 GN=CELSR1 PE=1 SV=1, ADGRC1 – TL EXPOSED**

ASFAVLMDISRRENGEVLPLKIVTYAAVSLSLAALLVAFVLLSLVRMLRSNLHSIHKHLAVALFLSQLVFVI

GINQTENPFLCTVVAILLHYIYMSTFAWTLVESLHVYRMLTEVRNIDTGPMRFYYVVGWG

IPAIVTGLAVGLDPQGYGNPDFCWLSLQDTLIWSFAGPIGAVIIINTVTSVLSAKVSCQR

KHHYYGKKGIVSLLRTAFLLLLLISATWLLGLLAVNRDALSFHYLFAIFSGLQGPFVLLF

HCVLNQEVRKHLKGVLGGRKLHLEDSATTRATLLTRSLNCNTTFGDGPDMLRTDLGESTA

SLDSIVRDEGIQKLGVSSGLVRGSHGEPDASLMPRSCKDPPGHDSDSDSELSLDEQSSSY

ASSHSSDSEDDGVGAEEKWDPARGAVHSTPKGDAVANHVPAGWPDQSLAESDSEDPSGKP

RLKVETKVSVELHREEQGSHRGEYPPDQESGGAARLASSQPPEQRKGILKNKVTYPPPLT

LTEQTLKGRLREKLADCEQSPTSSRTSSLGSGGPDCAITVKSPGREPGRDHLNGVAMNVR

TGSAQADGSDSEKP

**>sp|Q9HCU4|CELR2_HUMAN Cadherin EGF LAG seven-pass G-type receptor 2 OS=Homo sapiens OX=9606 GN=CELSR2 PE=1 SV=1**

MRSPATGVPLPTPPPPLLLLLLLLLPPPLLGDQVGPCRSLGSRGRGSSGACAPMGWLCPS

SASNLWLYTSRCRDAGTELTGHLVPHHDGLRVWCPESEAHIPLPPAPEGCPWSCRLLGIG

GHLSPQGKLTLPEEHPCLKAPRLRCQSCKLAQAPGLRAGERSPEESLGGRRKRNVNTAPQ

FQPPSYQATVPENQPAGTPVASLRAIDPDEGEAGRLEYTMDALFDSRSNQFFSLDPVTGA

VTTAEELDRETKSTHVFRVTAQDHGMPRRSALATLTILVTDTNDHDPVFEQQEYKESLRE

NLEVGYEVLTVRATDGDAPPNANILYRLLEGSGGSPSEVFEIDPRSGVIRTRGPVDREEV

ESYQLTVEASDQGRDPGPRSTTAAVFLSVEDDNDNAPQFSEKRYVVQVREDVTPGAPVLR

VTASDRDKGSNAVVHYSIMSGNARGQFYLDAQTGALDVVSPLDYETTKEYTLRVRAQDGG

RPPLSNVSGLVTVQVLDINDNAPIFVSTPFQATVLESVPLGYLVLHVQAIDADAGDNARL

EYRLAGVGHDFPFTINNGTGWISVAAELDREEVDFYSFGVEARDHGTPALTASASVSVTV

LDVNDNNPTFTQPEYTVRLNEDAAVGTSVVTVSAVDRDAHSVITYQITSGNTRNRFSITS

QSGGGLVSLALPLDYKLERQYVLAVTASDGTRQDTAQIVVNVTDANTHRPVFQSSHYTVN

VNEDRPAGTTVVLISATDEDTGENARITYFMEDSIPQFRIDADTGAVTTQAELDYEDQVS

YTLAITARDNGIPQKSDTTYLEILVNDVNDNAPQFLRDSYQGSVYEDVPPFTSVLQISAT

DRDSGLNGRVFYTFQGGDDGDGDFIVESTSGIVRTLRRLDRENVAQYVLRAYAVDKGMPP

ARTPMEVTVTVLDVNDNPPVFEQDEFDVFVEENSPIGLAVARVTATDPDEGTNAQIMYQI

VEGNIPEVFQLDIFSGELTALVDLDYEDRPEYVLVIQATSAPLVSRATVHVRLLDRNDNP

PVLGNFEILFNNYVTNRSSSFPGGAIGRVPAHDPDISDSLTYSFERGNELSLVLLNASTG

ELKLSRALDNNRPLEAIMSVLVSDGVHSVTAQCALRVTIITDEMLTHSITLRLEDMSPER

FLSPLLGLFIQAVAATLATPPDHVVVFNVQRDTDAPGGHILNVSLSVGQPPGPGGGPPFL

PSEDLQERLYLNRSLLTAISAQRVLPFDDNICLREPCENYMRCVSVLRFDSSAPFIASSS

VLFRPIHPVGGLRCRCPPGFTGDYCETEVDLCYSRPCGPHGRCRSREGGYTCLCRDGYTG

EHCEVSARSGRCTPGVCKNGGTCVNLLVGGFKCDCPSGDFEKPYCQVTTRSFPAHSFITF

RGLRQRFHFTLALSFATKERDGLLLYNGRFNEKHDFVALEVIQEQVQLTFSAGESTTTVS

PFVPGGVSDGQWHTVQLKYYNKPLLGQTGLPQGPSEQKVAVVTVDGCDTGVALRFGSVLG

NYSCAAQGTQGGSKKSLDLTGPLLLGGVPDLPESFPVRMRQFVGCMRNLQVDSRHIDMAD

FIANNGTVPGCPAKKNVCDSNTCHNGGTCVNQWDAFSCECPLGFGGKSCAQEMANPQHFL

GSSLVAWHGLSLPISQPWYLSLMFRTRQADGVLLQAITRGRSTITLQLREGHVMLSVEGT

GLQASSLRLEPGRANDGDWHHAQLALGASGGPGHAILSFDYGQQRAEGNLGPRLHGLHLS

NITVGGIPGPAGGVARGFRGCLQGVRVSDTPEGVNSLDPSHGESINVEQGCSLPDPCDSN

PCPANSYCSNDWDSYSCSCDPGYYGDNCTNVCDLNPCEHQSVCTRKPSAPHGYTCECPPN

YLGPYCETRIDQPCPRGWWGHPTCGPCNCDVSKGFDPDCNKTSGECHCKENHYRPPGSPT

CLLCDCYPTGSLSRVCDPEDGQCPCKPGVIGRQCDRCDNPFAEVTTNGCEVNYDSCPRAI

EAGIWWPRTRFGLPAAAPCPKGSFGTAVRHCDEHRGWLPPNLFNCTSITFSELKGFAERL

QRNESGLDSGRSQQLALLLRNATQHTAGYFGSDVKVAYQLATRLLAHESTQRGFGLSATQ

DVHFTENLLRVGSALLDTANKRHWELIQQTEGGTAWLLQHYEAYASALAQNMRHTYLSPF

TIVTPNIVISVVRLDKGNFAGAKLPRYEALRGEQPPDLETTVILPESVFRETPPVVRPAG

PGEAQEPEELARRQRRHPELSQGEAVASVIIYRTLAGLLPHNYDPDKRSLRVPKRPIINT

PVVSISVHDDEELLPRALDKPVTVQFRLLETEERTKPICVFWNHSILVSGTGGWSARGCE

VVFRNESHVSCQCNHMTSFAVLMDVSRRENGEILPLKTLTYVALGVTLAALLLTFFFLTL

LRILRSNQHGIRRNLTAALGLAQLVFLLGINQADLPFACTVIAILLHFLYLCTFSWALLE

ALHLYRALTEVRDVNTGPMRFYYMLGWGVPAFITGLAVGLDPEGYGNPDFCWLSIYDTLI

WSFAGPVAFAVSMSVFLYILAARASCAAQRQGFEKKGPVSGLQPSFAVLLLLSATWLLAL

LSVNSDTLLFHYLFATCNCIQGPFIFLSYVVLSKEVRKALKLACSRKPSPDPALTTKSTL

TSSYNCPSPYADGRLYQPYGDSAGSLHSTSRSGKSQPSYIPFLLREESALNPGQGPPGLG

DPGSLFLEGQDQQHDPDTDSDSDLSLEDDQSGSYASTHSSDSEEEEEEEEEEAAFPGEQG

WDSLLGPGAERLPLHSTPKDGGPGPGKAPWPGDFGTTAKESSGNGAPEERLRENGDALSR

EGSLGPLPGSSAQPHKGILKKKCLPTISEKSSLLRLPLEQCTGSSRGSSASEGSRGGPPP

RPPPRQSLQEQLNGVMPIAMSIKAGTVDEDSSGSEFLFFNFLH

**>sp|Q9HCU4|CELR2_HUMAN Cadherin EGF LAG seven-pass G-type receptor 2 OS=Homo sapiens OX=9606 GN=CELSR2 PE=1 SV=1 – TL EXPOSED**

TSFAVLMDVSRRENGEILPLKTLTYVALGVTLAALLLTFFFLTL

LRILRSNQHGIRRNLTAALGLAQLVFLLGINQADLPFACTVIAILLHFLYLCTFSWALLE

ALHLYRALTEVRDVNTGPMRFYYMLGWGVPAFITGLAVGLDPEGYGNPDFCWLSIYDTLI

WSFAGPVAFAVSMSVFLYILAARASCAAQRQGFEKKGPVSGLQPSFAVLLLLSATWLLAL

LSVNSDTLLFHYLFATCNCIQGPFIFLSYVVLSKEVRKALKLACSRKPSPDPALTTKSTL

TSSYNCPSPYADGRLYQPYGDSAGSLHSTSRSGKSQPSYIPFLLREESALNPGQGPPGLG

DPGSLFLEGQDQQHDPDTDSDSDLSLEDDQSGSYASTHSSDSEEEEEEEEEEAAFPGEQG

WDSLLGPGAERLPLHSTPKDGGPGPGKAPWPGDFGTTAKESSGNGAPEERLRENGDALSR

EGSLGPLPGSSAQPHKGILKKKCLPTISEKSSLLRLPLEQCTGSSRGSSASEGSRGGPPP

RPPPRQSLQEQLNGVMPIAMSIKAGTVDEDSSGSEFLFFNFLH

**>sp|Q91ZI0|CELR3_MOUSE Cadherin EGF LAG seven-pass G-type receptor 3 OS=Mus musculus OX=10090 GN=Celsr3 PE=2 SV=2**

MARRPLWWGLPGPSTPVLLLLLLSLFPFSREELGGGGDQDWDPGVATTTGPRAQIGSGAV

ALCPESPGVWEDGDPGLGVREPVFMRLRVGRQNARNGRGAPEQPNAEVVVQALGSREQEA

GQGPGYLLCWHPEISSCGRTGPLRRGSLPLDALSPGDSDLRNSSPHPSELLAQPDGSRPV

AFQRNARRSIRKRVETSRCCGKLWEPGHKGQGERSATSTVDRGPFRRDCLPGSLGSGLGE

DSAPRAVRTAPTPGSAPRESRTAPGRMRSRGLFRRRFLFERPGPRPPGFPTGPEAKQILS

TNQARPRRAANRHPQFPQYNYQTLVPENEAAGTSVLRVVAQDPDPGEAGRLIYSLAALMN

SRSLELFSIDPQSGLIRTAAALDRESMERHYLRVTAQDHGSPRLSATTMVAVTVADRNDH

APVFEQAQYRETLRENVEEGYPILQLRATDGDAPPNANLRYRFVGSPAVRTAAAAAFEID

PRSGLISTSGRVDREHMESYELVVEASDQGQEPGPRSATVRVHITVLDENDNAPQFSEKR

YVAQVREDVRPHTVVLRVTATDKDKDANGLVHYNIISGNSRGHFAIDSLTGEIQVMAPLD

FEAEREYALRIRAQDAGRPPLSNNTGLASIQVVDINDHAPIFVSTPFQVSVLENAPLGHS

VIHIQAVDADHGENSRLEYSLTGVASDTPFVINSATGWVSVSGPLDRESVEHYFFGVEAR

DHGSPPLSASASVTVTVLDVNDNRPEFTMKEYHLRLNEDAAVGTSVVSVTAVDRDANSAI

SYQITGGNTRNRFAISTQGGVGLVTLALPLDYKQERYFKLVLTASDRALHDHCYVHINIT

DANTHRPVFQSAHYSVSMNEDRPVGSTVVVISASDDDVGENARITYLLEDNLPQFRIDAD

SGAITLQAPLDYEDQVTYTLAITARDNGIPQKADTTYVEVMVNDVNDNAPQFVASHYTGL

VSEDAPPFTSVLQISATDRDAHANGRVQYTFQNGEDGDGDFTIEPTSGIVRTVRRLDREA

VPVYELTAYAVDRGVPPLRTPVSIQVTVQDVNDNAPVFPAEEFEVRVKENSIVGSVVAQI

TAVDPDDGPNAHIMYQIVEGNIPELFQMDIFSGELTALIDLDYEARQEYVIVVQATSAPL

VSRATVHVRLVDQNDNSPVLNNFQILFNNYVSNRSDTFPSGIIGRIPAYDPDVSDHLFYS

FERGNELQLLVVNRTSGELRLSRKLDNNRPLVASMLVTVTDGLHSVTAQCVLRVVIITEE

LLANSLTVRLENMWQERFLSPLLGHFLEGVAAVLATPTEDVFIFNIQNDTDVGGTVLNVS

FSALAPRGAGAGAAGPWFSSEELQEQLYVRRAALAARSLLDVLPFDDNVCLREPCENYMK

CVSVLRFDSSAPFLASTSTLFRPIQPIAGLRCRCPPGFTGDFCETELDLCYSNPCRNGGA

CARREGGYTCVCRPRFTDCELDTEAGRCVPGVCRNGGTCTNAPNGGFRCQCPAGGAFEGP

RCEVAARSFPPSSFVMFRGLRQRFHLTLSLSFATVQPSGLLFYNGRLNEKHDFLALELVA

GQVRLTYSTGESNTVVSPTVPGGLSDGQWHTVHLRYYNKPRTDALGGAQGPSKDKVAVLS

VDDCNVAVALQFGAEIGNYSCAAAGVQTSSKKSLDLTGPLLLGGVPNLPENFPVSHKDFI

GCMRDLHIDGRRMDMAAFVANNGTMAGCQAKSHFCASGPCKNNGFCSERWGGFSCDCPVG

FGGKDCRLTMAHPYHFQGNGTLSWDFGNDMAVSVPWYLGLSFRTRATKGILMQVQLGPHS

VLLCKLDRGLLSVTLNRASGHTVHLLLDQMTVSDGRWHDLRLELQEEPGGRRGHHIFMVS

LDFTLFQDTMAMGGELQGLKVKQLHVGGLPPSSKEEGHQGLVGCIQGVWIGFTPFGSSAL

LPPSHRVNVEPGCTVTNPCASGPCPPHADCKDLWQTFSCTCRPGYYGPGCVDACLLNPCQ

NQGSCRHLQGAPHGYTCDCVSGYFGQHCEHRVDQQCPRGWWGSPTCGPCNCDVHKGFDPN

CNKTNGQCHCKEFHYRPRGSDSCLPCDCYPVGSTSRSCAPHSGQCPCRPGALGRQCNSCD

SPFAEVTASGCRVLYDACPKSLRSGVWWPQTKFGVLATVPCPRGALGAAVRLCDEDQGWL

EPDLFNCTSPAFRELSLLLDGLELNKTALDTVEAKKLAQRLREVTGQTDHYFSQDVRVTA

RLLAYLLAFESHQQGFGLTATQDAHFNENLLWAGSALLAPETGHLWAALGQRAPGGSPGS

AGLVQHLEEYAATLARNMELTYLNPVGLVTPNIMLSIDRMEHPSSTQGARRYPRYHSNLF

RGQDAWDPHTHVLLPSQASQPSPSEVLPTSSNAENATASSVVSPPAPLEPESEPGISIVI

LLVYRALGGLLPAQFQAERRGARLPQNPVMNSPVVSVAVFHGRNFLRGVLVSPINLEFRL

LQTANRSKAICVQWDPPGPTDQHGMWTARDCELVHRNGSHARCRCSRTGTFGVLMDASPR

ERLEGDLELLAVFTHVVVAVSVTALVLTAAVLLSLRSLKSNVRGIHANVAAALGVAELLF

LLGIHRTHNQLLCTAVAILLHYFFLSTFAWLLVQGLHLYRMQVEPRNVDRGAMRFYHALG

WGVPAVLLGLAVGLDPEGYGNPDFCWISIHEPLIWSFAGPIVLVIVMNGTMFLLAARTSC

STGQREAKKTSVLTLRSSFLLLLLVSASWLFGLLAVNHSILAFHYLHAGLCGLQGLAVLL

LFCVLNADARAAWTPACLGKKAAPEETRPAPGPGSGAYNNTALFEESGLIRITLGASTVS

SVSSARSGRAQDQDSQRGRSYLRDNVLVRHGSTAEHTERSLQAHAGPTDLDVAMFHRDAG

ADSDSDSDLSLEEERSLSIPSSESEDNGRTRGRFQRPLRRAAQSERLLAHPKDVDGNDLL

SYWPALGECEAAPCALQAWGSERRLGLDSNKDAANNNQPELALTSGDETSLGRAQRQRKG

ILKNRLQYPLVPQSRGTPELSWCRAATLGHRAVPAASYGRIYAGGGTGSLSQPASRYSSR

EQLDLLLRRQLSKERLEEVPVPAPVLHPLSRPGSQERLDTAPARLEARDRGSTLPRRQPP

RDYPGTMAGRFGSRDALDLGAPREWLSTLPPPRRNRDLDPQHPPLPLSPQRQLSRDPLLP

SRPLDSLSRISNSREGLDQVPSRHPSREALGPAPQLLRAREDPASGPSHGPSTEQLDILS

SILASFNSSALSSVQSSSTPSGPHTTATASALGPSTPRSATSHSISELSPDSEVPRSEGHS

**>sp|Q91ZI0|CELR3_MOUSE Cadherin EGF LAG seven-pass G-type receptor 3 OS=Mus musculus OX=10090 GN=Celsr3 PE=2 SV=2 – TL EXPOSED**

GTFGVLMDASPRERLEGDLELLAVFTHVVVAVSVTALVLTAAVLLSLRSLKSNVRGIHAN

VAAALGVAELLFLLGIHRTHNQLLCTAVAILLHYFFLSTFAWLLVQGLHLYRMQVEPRNV

DRGAMRFYHALGWGVPAVLLGLAVGLDPEGYGNPDFCWISIHEPLIWSFAGPIVLVIVMN

GTMFLLAARTSCSTGQREAKKTSVLTLRSSFLLLLLVSASWLFGLLAVNHSILAFHYLHA

GLCGLQGLAVLLLFCVLNADARAAWTPACLGKKAAPEETRPAPGPGSGAYNNTALFEESG

LIRITLGASTVSSVSSARSGRAQDQDSQRGRSYLRDNVLVRHGSTAEHTERSLQAHAGPT

DLDVAMFHRDAGADSDSDSDLSLEEERSLSIPSSESEDNGRTRGRFQRPLRRAAQSERLL

AHPKDVDGNDLLSYWPALGECEAAPCALQAWGSERRLGLDSNKDAANNNQPELALTSGDE

TSLGRAQRQRKGILKNRLQYPLVPQSRGTPELSWCRAATLGHRAVPAASYGRIYAGGGTG

SLSQPASRYSSREQLDLLLRRQLSKERLEEVPVPAPVLHPLSRPGSQERLDTAPARLEAR

DRGSTLPRRQPPRDYPGTMAGRFGSRDALDLGAPREWLSTLPPPRRNRDLDPQHPPLPLS

PQRQLSRDPLLPSRPLDSLSRISNSREGLDQVPSRHPSREALGPAPQLLRAREDPASGPS

HGPSTEQLDILSSILASFNSSALSSVQSSSTPSGPHTTATASALGPSTPRSATSHSISEL

SPDSEVPRSEGHS

**>sp|Q8WXG9|AGRV1_HUMAN Adhesion G-protein coupled receptor V1 OS=Homo sapiens OX=9606 GN=ADGRV1 PE=1 SV=2**

MSVFLGPGMPSASLLVNLLSALLILFVFGETEIRFTGQTEFVVNETSTTVIRLIIERIGE

PANVTAIVSLYGEDAGDFFDTYAAAFIPAGETNRTVYIAVCDDDLPEPDETFIFHLTLQK

PSANVKLGWPRTVTVTILSNDNAFGIISFNMLPSIAVSEPKGRNESMPLTLIREKGTYGM

VMVTFEVEGGPNPPDEDLSPVKGNITFPPGRATVIYNLTVLDDEVPENDEIFLIQLKSVE

GGAEINTSRNSIEIIIKKNDSPVRFLQSIYLVPEEDHILIIPVVRGKDNNGNLIGSDEYE

VSISYAVTTGNSTAHAQQNLDFIDLQPNTTVVFPPFIHESHLKFQIVDDTIPEIAESFHI

MLLKDTLQGDAVLISPSVVQVTIKPNDKPYGVLSFNSVLFERTVIIDEDRISRYEEITVV

RNGGTHGNVSANWVLTRNSTDPSPVTADIRPSSGVLHFAQGQMLATIPLTVVDDDLPEEA

EAYLLQILPHTIRGGAEVSEPAELLFYIQDSDDVYGLITFFPMENQKIESSPGERYLSLS

FTRLGGTKGDVRLLYSVLYIPAGAVDPLQAKEGILNISRRNDLIFPEQKTQVTTKLPIRN

DAFLQNGAHFLVQLETVELLNIIPLIPPISPRFGEICNISLLVTPAIANGEIGFLSNLPI

ILHEPEDFAAEVVYIPLHRDGTDGQATVYWSLKPSGFNSKAVTPDDIGPFNGSVLFLSGQ

SDTTINITIKGDDIPEMNETVTLSLDRVNVENQVLKSGYTSRDLIILENDDPGGVFEFSP

ASRGPYVIKEGESVELHIIRSRGSLVKQFLHYRVEPRDSNEFYGNTGVLEFKPGEREIVI

TLLARLDGIPELDEHYWVVLSSHGERESKLGSATIVNITILKNDDPHGIIEFVSDGLIVM

INESKGDAIYSAVYDVVRNRGNFGDVSVSWVVSPDFTQDVFPVQGTVVFGDQEFSKNITI

YSLPDEIPEEMEEFTVILLNGTGGAKVGNRTTATLRIRRNDDPIYFAEPRVVRVQEGETA

NFTVLRNGSVDVTCMVQYATKDGKATARERDFIPVEKGETLIFEVGSRQQSISIFVNEDG

IPETDEPFYIILLNSTGDTVVYQYGVATVIIEANDDPNGIFSLEPIDKAVEEGKTNAFWI

LRHRGYFGSVSVSWQLFQNDSALQPGQEFYETSGTVNFMDGEEAKPIILHAFPDKIPEFN

EFYFLKLVNISGGSPGPGGQLAETNLQVTVMVPFNDDPFGVFILDPECLEREVAEDVLSE

DDMSYITNFTILRQQGVFGDVQLGWEILSSEFPAGLPPMIDFLLVGIFPTTVHLQQHMRR

HHSGTDALYFTGLEGAFGTVNPKYHPSRNNTIANFTFSAWVMPNANTNGFIIAKDDGNGS

IYYGVKIQTNESHVTLSLHYKTLGSNATYIAKTTVMKYLEESVWLHLLIILEDGIIEFYL

DGNAMPRGIKSLKGEAITDGPGILRIGAGINGNDRFTGLMQDVRSYERKLTLEEIYELHA

MPAKSDLHPISGYLEFRQGETNKSFIISARDDNDEEGEELFILKLVSVYGGARISEENTT

ARLTIQKSDNANGLFGFTGACIPEIAEEGSTISCVVERTRGALDYVHVFYTISQIETDGI

NYLVDDFANASGTITFLPWQRSEVLNIYVLDDDIPELNEYFRVTLVSAIPGDGKLGSTPT

SGASIDPEKETTDITIKASDHPYGLLQFSTGLPPQPKDAMTLPASSVPHITVEEEDGEIR

LLVIRAQGLLGRVTAEFRTVSLTAFSPEDYQNVAGTLEFQPGERYKYIFINITDNSIPEL

EKSFKVELLNLEGGVAELFRVDGSGSGDGDMEFFLPTIHKRASLGVASQILVTIAASDHA

HGVFEFSPESLFVSGTEPEDGYSTVTLNVIRHHGTLSPVTLHWNIDSDPDGDLAFTSGNI

TFEIGQTSANITVEILPDEDPELDKAFSVSVLSVSSGSLGAHINATLTVLASDDPYGIFI

FSEKNRPVKVEEATQNITLSIIRLKGLMGKVLVSYATLDDMEKPPYFPPNLARATQGRDY

IPASGFALFGANQSEATIAISILDDDEPERSESVFIELLNSTLVAKVQSRSIPNSPRLGP

KVETIAQLIIIANDDAFGTLQLSAPIVRVAENHVGPIINVTRTGGAFADVSVKFKAVPIT

AIAGEDYSIASSDVVLLEGETSKAVPIYVINDIYPELEESFLVQLMNETTGGARLGALTE

AVIIIEASDDPYGLFGFQITKLIVEEPEFNSVKVNLPIIRNSGTLGNVTVQWVATINGQL

ATGDLRVVSGNVTFAPGETIQTLLLEVLADDVPEIEEVIQVQLTDASGGGTIGLDRIANI

IIPANDDPYGTVAFAQMVYRVQEPLERSSCANITVRRSGGHFGRLLLFYSTSDIDVVALA

MEEGQDLLSYYESPIQGVPDPLWRTWMNVSAVGEPLYTCATLCLKEQACSAFSFFSASEG

PQCFWMTSWISPAVNNSDFWTYRKNMTRVASLFSGQAVAGSDYEPVTRQWAIMQEGDEFA

NLTVSILPDDFPEMDESFLISLLEVHLMNISASLKNQPTIGQPNISTVVIALNGDAFGVF

VIYNISPNTSEDGLFVEVQEQPQTLVELMIHRTGGSLGQVAVEWRVVGGTATEGLDFIGA

GEILTFAEGETKKTVILTILDDSEPEDDESIIVSLVYTEGGSRILPSSDTVRVNILANDN

VAGIVSFQTASRSVIGHEGEILQFHVIRTFPGRGNVTVNWKIIGQNLELNFANFSGQLFF

PEGSLNTTLFVHLLDDNIPEEKEVYQVILYDVRTQGVPPAGIALLDAQGYAAVLTVEASD

EPHGVLNFALSSRFVLLQEANITIQLFINREFGSLGAINVTYTTVPGMLSLKNQTVGNLA

EPEVDFVPIIGFLILEEGETAAAINITILEDDVPELEEYFLVNLTYVGLTMAASTSFPPR

LDSEGLTAQVIIDANDGARGVIEWQQSRFEVNETHGSLTLVAQRSREPLGHVSLFVYAQN

LEAQVGLDYIFTPMILHFADGERYKNVNIMILDDDIPEGDEKFQLILTNPSPGLELGKNT

IALIIVLANDDGPGVLSFNNSEHFFLREPTALYVQESVAVLYIVREPAQGLFGTVTVQFI

VTEVNSSNESKDLTPSKGYIVLEEGVRFKALQISAILDTEPEMDEYFVCTLFNPTGGARL

GVHVQTLITVLQNQAPLGLFSISAVENRATSIDIEEANRTVYLNVSRTNGIDLAVSVQWE

TVSETAFGMRGMDVVFSVFQSFLDESASGWCFFTLENLIYGIMLRKSSVTVYRWQGIFIP

VEDLNIENPKTCEAFNIGFSPYFVITHEERNEEKPSLNSVFTFTSGFKLFLVQTIIILES

SQVRYFTSDSQDYLIIASQRDDSELTQVFRWNGGSFVLHQKLPVRGVLTVALFNKGGSVF

LAISQANARLNSLLFRWSGSGFINFQEVPVSGTTEVEALSSANDIYLIFAENVFLGDQNS

IDIFIWEMGQSSFRYFQSVDFAAVNRIHSFTPASGIAHILLIGQDMSALYCWNSERNQFS

FVLEVPSAYDVASVTVKSLNSSKNLIALVGAHSHIYELAYISSHSDFIPSSGELIFEPGE

REATIAVNILDDTVPEKEESFKVQLKNPKGGAEIGINDSVTITILSNDDAYGIVAFAQNS

LYKQVEEMEQDSLVTLNVERLKGTYGRITIAWEADGSISDIFPTSGVILFTEGQVLSTIT

LTILADNIPELSEVVIVTLTRITTEGVEDSYKGATIDQDRSKSVITTLPNDSPFGLVGWR

AASVFIRVAEPKENTTTLQLQIARDKGLLGDIAIHLRAQPNFLLHVDNQATENEDYVLQE

TIIIMKENIKEAHAEVSILPDDLPELEEGFIVTITEVNLVNSDFSTGQPSVRRPGMEIAE

IMIEENDDPRGIFMFHVTRGAGEVITAYEVPPPLNVLQVPVVRLAGSFGAVNVYWKASPD

SAGLEDFKPSHGILEFADKQVTAMIEITIIDDAEFELTETFNISLISVAGGGRLGDDVVV

TVVIPQNDSPFGVFGFEEKTVMIDESLSSDDPDSYVTLTVVRSPGGKGTVRLEWTIDEKA

KHNLSPLNGTLHFDETESQKTIVLHTLQDTVLEEDRRFTIQLISIDEVEISPVKGSASII

IRGDKRASGEVGIAPSSRHILIGEPSAKYNGTAIISLVRGPGILGEVTVFWRIFPPSVGE

FAETSGKLTMRDEQSAVIVVIQALNDDIPEEKSFYEFQLTAVSEGGVLSESSSTANITVV

ASDSPYGRFAFSHEQLRVSEAQRVNITIIRSSGDFGHVRLWYKTMSGTAEAGLDFVPAAG

ELLFEAGEMRKSLHVEILDDDYPEGPEEFSLTITKVELQGRGYDFTIQENGLQIDQPPEI

GNISIVRIIIMKNDNAEGIIEFDPKYTAFEVEEDVGLIMIPVVRLHGTYGYVTADFISQS

SSASPGGVDYILHGSTVTFQHGQNLSFINISIIDDNESEFEEPIEILLTGATGGAVLGRH

LVSRIIIAKSDSPFGVIRFLNQSKISIANPNSTMILSLVLERTGGLLGEIQVNWETVGPN

SQEALLPQNRDIADPVSGLFYFGEGEGGVRTIILTIYPHEEIEVEETFIIKLHLVKGEAK

LDSRAKDVTLTIQEFGDPNGVVQFAPETLSKKTYSEPLALEGPLLITFFVRRVKGTFGEI

MVYWELSSEFDITEDFLSTSGFFTIADGESEASFDVHLLPDEVPEIEEDYVIQLVSVEGG

AELDLEKSITWFSVYANDDPHGVFALYSDRQSILIGQNLIRSIQINITRLAGTFGDVAVG

LRISSDHKEQPIVTENAERQLVVKDGATYKVDVVPIKNQVFLSLGSNFTLQLVTVMLVGG

RFYGMPTILQEAKSAVLPVSEKAANSQVGFESTAFQLMNITAGTSHVMISRRGTYGALSV

AWTTGYAPGLEIPEFIVVGNMTPTLGSLSFSHGEQRKGVFLWTFPSPGWPEAFVLHLSGV

QSSAPGGAQLRSGFIVAEIEPMGVFQFSTSSRNIIVSEDTQMIRLHVQRLFGFHSDLIKV

SYQTTAGSAKPLEDFEPVQNGELFFQKFQTEVDFEITIINDQLSEIEEFFYINLTSVEIR

GLQKFDVNWSPRLNLDFSVAVITILDNDDLAGMDISFPETTVAVAVDTTLIPVETESTTY

LSTSKTTTILQPTNVVAIVTEATGVSAIPEKLVTLHGTPAVSEKPDVATVTANVSIHGTF

SLGPSIVYIEEEMKNGTFNTAEVLIRRTGGFTGNVSITVKTFGERCAQMEPNALPFRGIY

GISNLTWAVEEEDFEEQTLTLIFLDGERERKVSVQILDDDEPEGQEFFYVFLTNPQGGAQ

IVEEKDDTGFAAFAMVIITGSDLHNGIIGFSEESQSGLELREGAVMRRLHLIVTRQPNRA

FEDVKVFWRVTLNKTVVVLQKDGVNLVEELQSVSGTTTCTMGQTKCFISIELKPEKVPQV

EVYFFVELYEATAGAAINNSARFAQIKILESDESQSLVYFSVGSRLAVAHKKATLISLQV

ARDSGTGLMMSVNFSTQELRSAETIGRTIISPAISGKDFVITEGTLVFEPGQRSTVLDVI

LTPETGSLNSFPKRFQIVLFDPKGGARIDKVYGTANITLVSDADSQAIWGLADQLHQPVN

DDILNRVLHTISMKVATENTDEQLSAMMHLIEKITTEGKIQAFSVASRTLFYEILCSLIN

PKRKDTRGFSHFAEVTENFAFSLLTNVTCGSPGEKSKTILDSCPYLSILALHWYPQQING

HKFEGKEGDYIRIPERLLDVQDAEIMAGKSTCKLVQFTEYSSQQWFISGNNLPTLKNKVL

SLSVKGQSSQLLTNDNEVLYRIYAAEPRIIPQTSLCLLWNQAAASWLSDSQFCKVVEETA

DYVECACSHMSVYAVYARTDNLSSYNEAFFTSGFICISGLCLAVLSHIFCARYSMFAAKL

LTHMMAASLGTQILFLASAYASPQLAEESCSAMAAVTHYLYLCQFSWMLIQSVNFWYVLV

MNDEHTERRYLLFFLLSWGLPAFVVILLIVILKGIYHQSMSQIYGLIHGDLCFIPNVYAA

LFTAALVPLTCLVVVFVVFIHAYQVKPQWKAYDDVFRGRTNAAEIPLILYLFALISVTWL

WGGLHMAYRHFWMLVLFVIFNSLQGLYVFMVYFILHNQMCCPMKASYTVEMNGHPGPSTA

FFTPGSGMPPAGGEISKSTQNLIGAMEEVPPDWERASFQQGSQASPDLKPSPQNGATFPS

SGGYGQGSLIADEESQEFDDLIFALKTGAGLSVSDNESGQGSQEGGTLTDSQIVELRRIP

IADTHL

**>sp|Q8WXG9|AGRV1_HUMAN Adhesion G-protein coupled receptor V1 OS=Homo sapiens OX=9606 GN=ADGRV1 PE=1 SV=2** **– TL EXPOSED**

SVYAVYARTDNLSSYNEAFFTSGFICISGLCLAVLSHIFCARYSMFAAKL

LTHMMAASLGTQILFLASAYASPQLAEESCSAMAAVTHYLYLCQFSWMLIQSVNFWYVLV

MNDEHTERRYLLFFLLSWGLPAFVVILLIVILKGIYHQSMSQIYGLIHGDLCFIPNVYAA

LFTAALVPLTCLVVVFVVFIHAYQVKPQWKAYDDVFRGRTNAAEIPLILYLFALISVTWL

WGGLHMAYRHFWMLVLFVIFNSLQGLYVFMVYFILHNQMCCPMKASYTVEMNGHPGPSTA

FFTPGSGMPPAGGEISKSTQNLIGAMEEVPPDWERASFQQGSQASPDLKPSPQNGATFPS

SGGYGQGSLIADEESQEFDDLIFALKTGAGLSVSDNESGQGSQEGGTLTDSQIVELRRIP

IADTHL
