## Supplementary figures and images for "AlphaFold prediction and analysis of Adhesion-family G protein coupled receptor (aGPCR) structures"

### af284_coverage.png

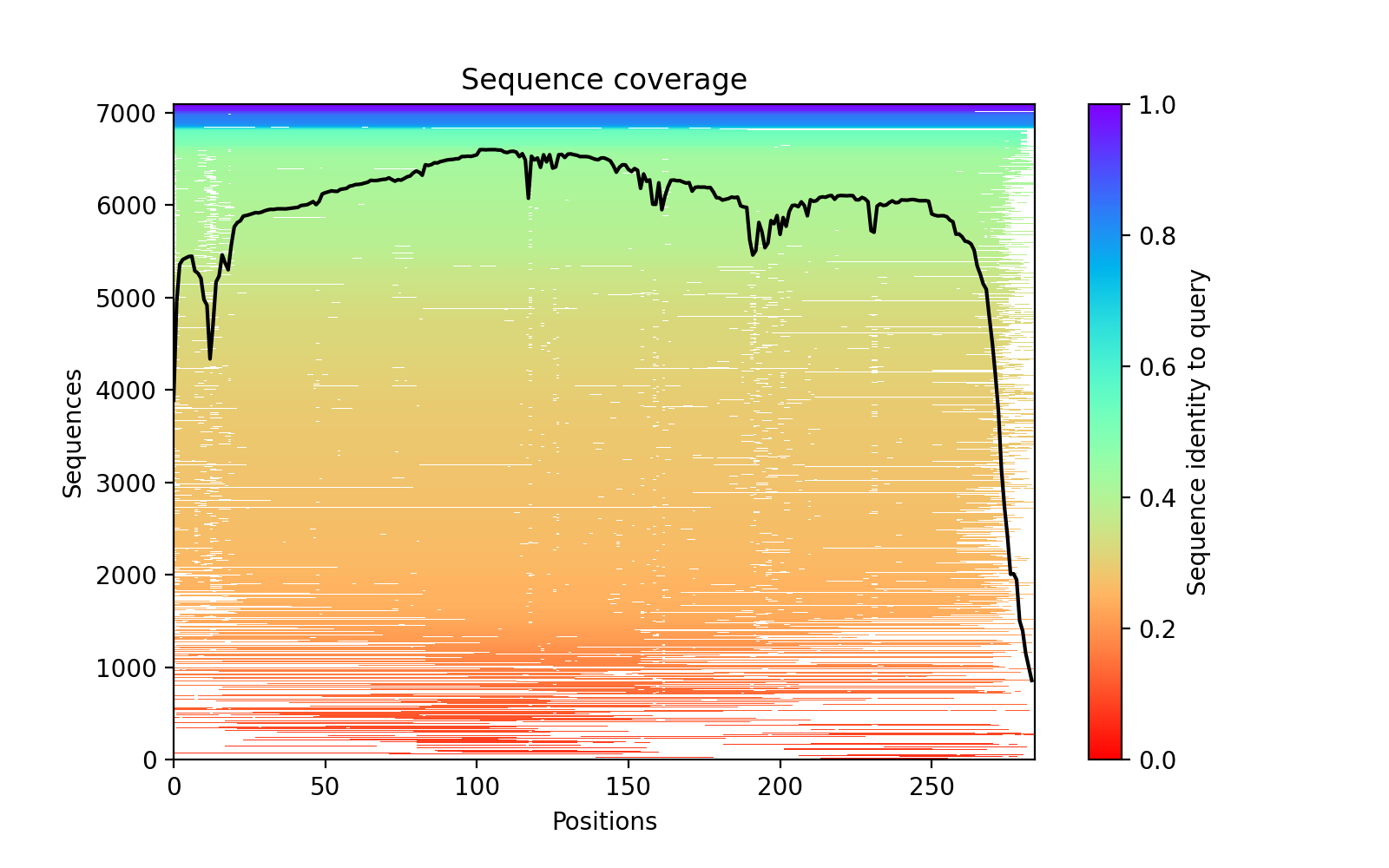

### af284_PAE.png

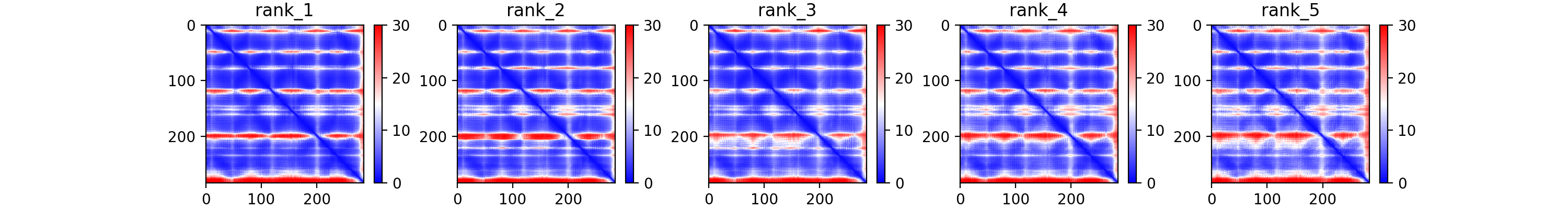

### af284_plddt.png

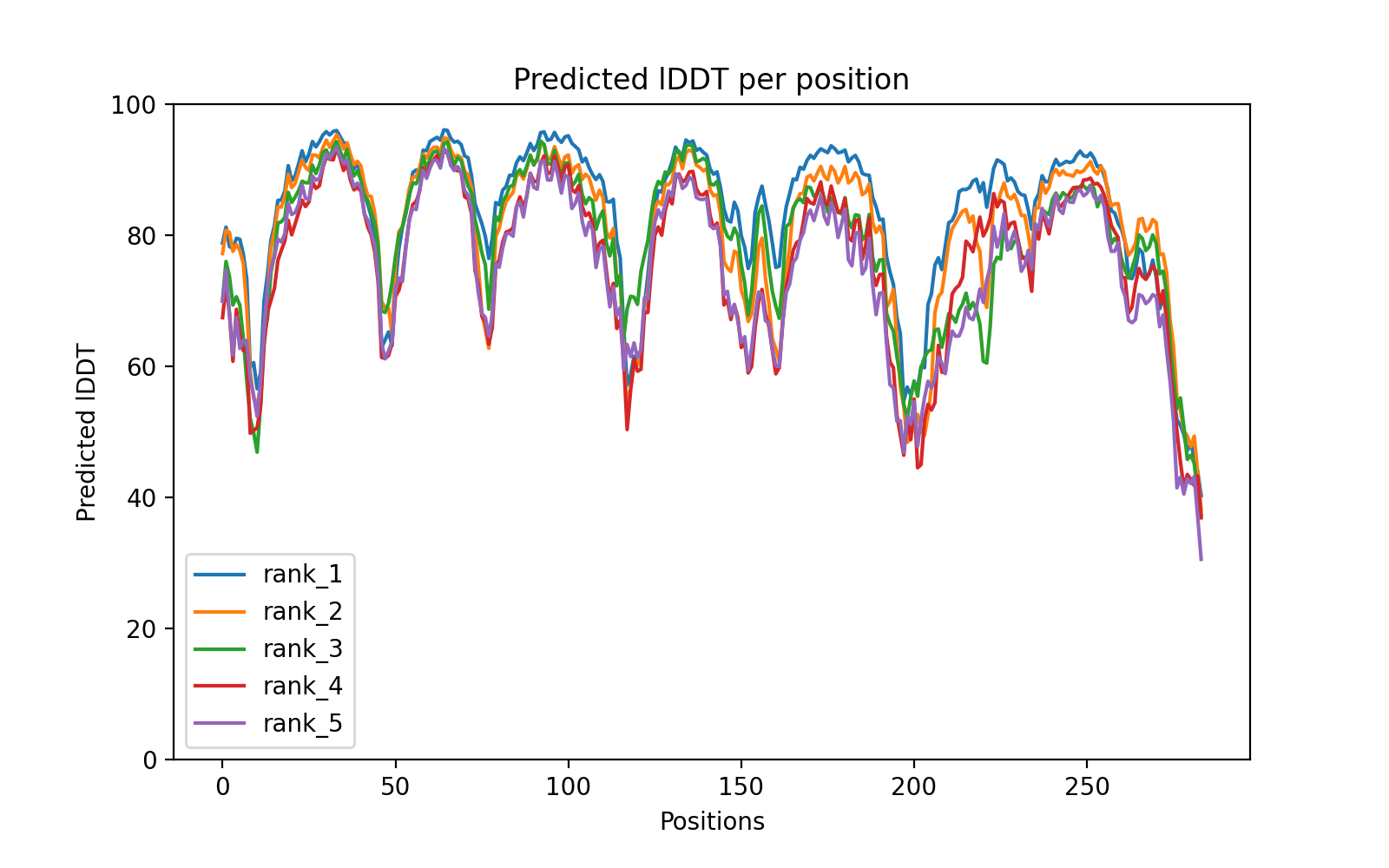

### af327_coverage.png

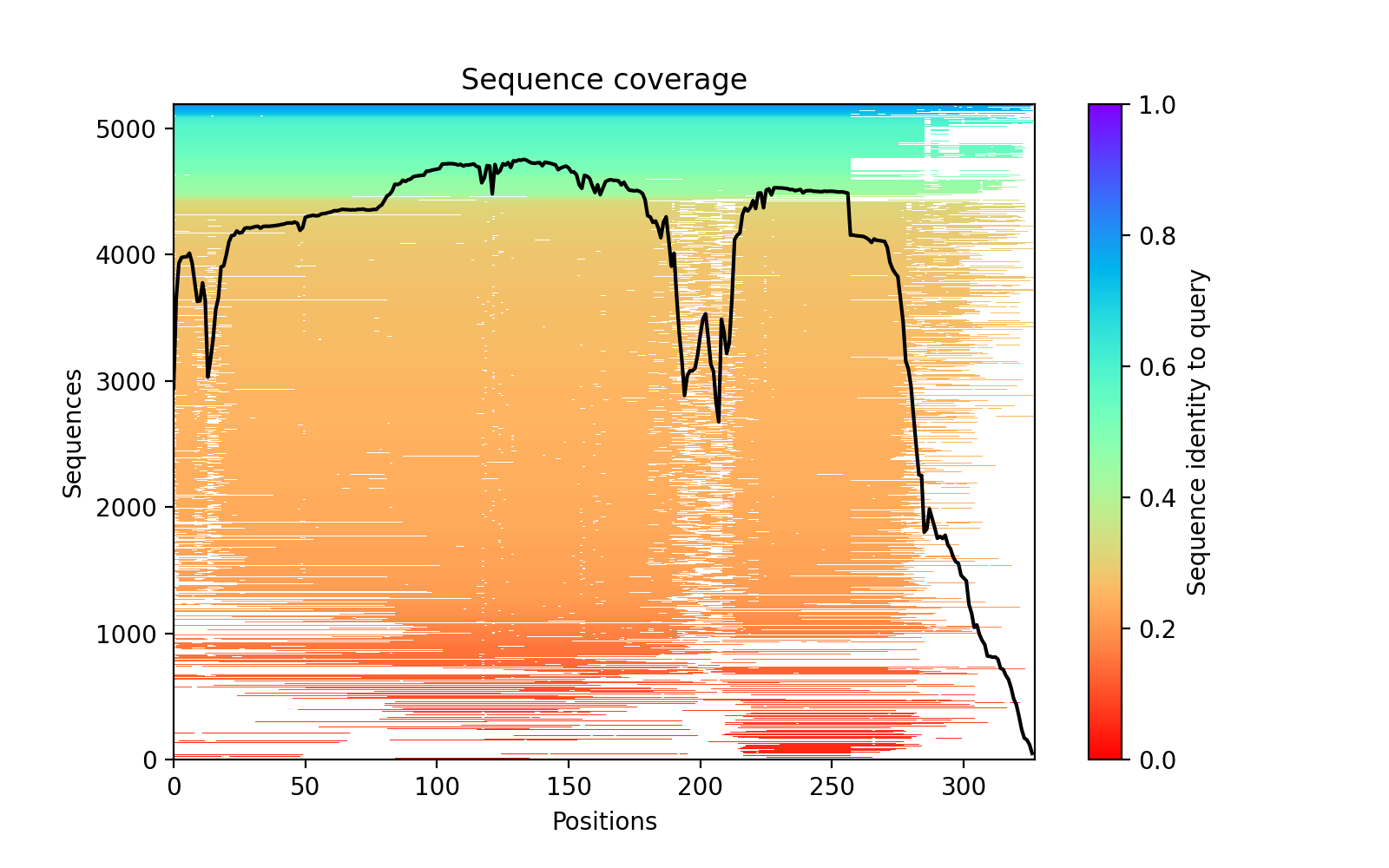

### af327_PAE.png

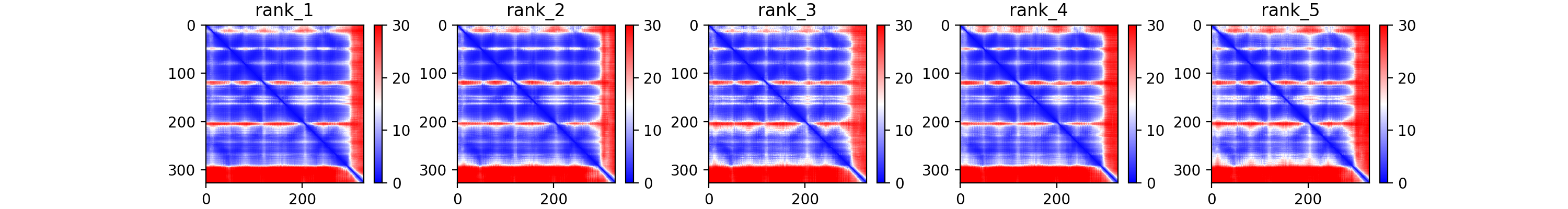

### af327_plddt.png

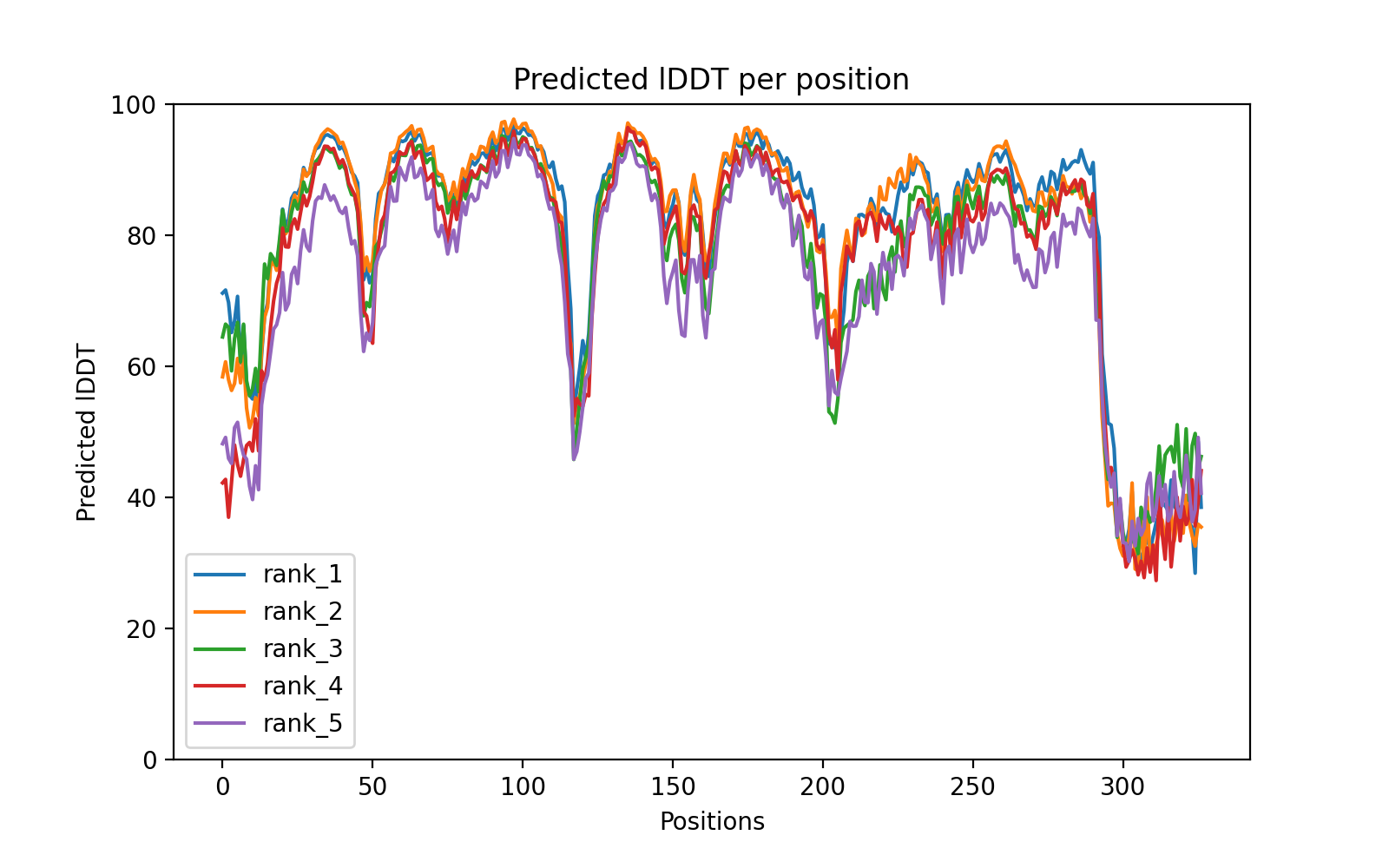

### af330_coverage.png

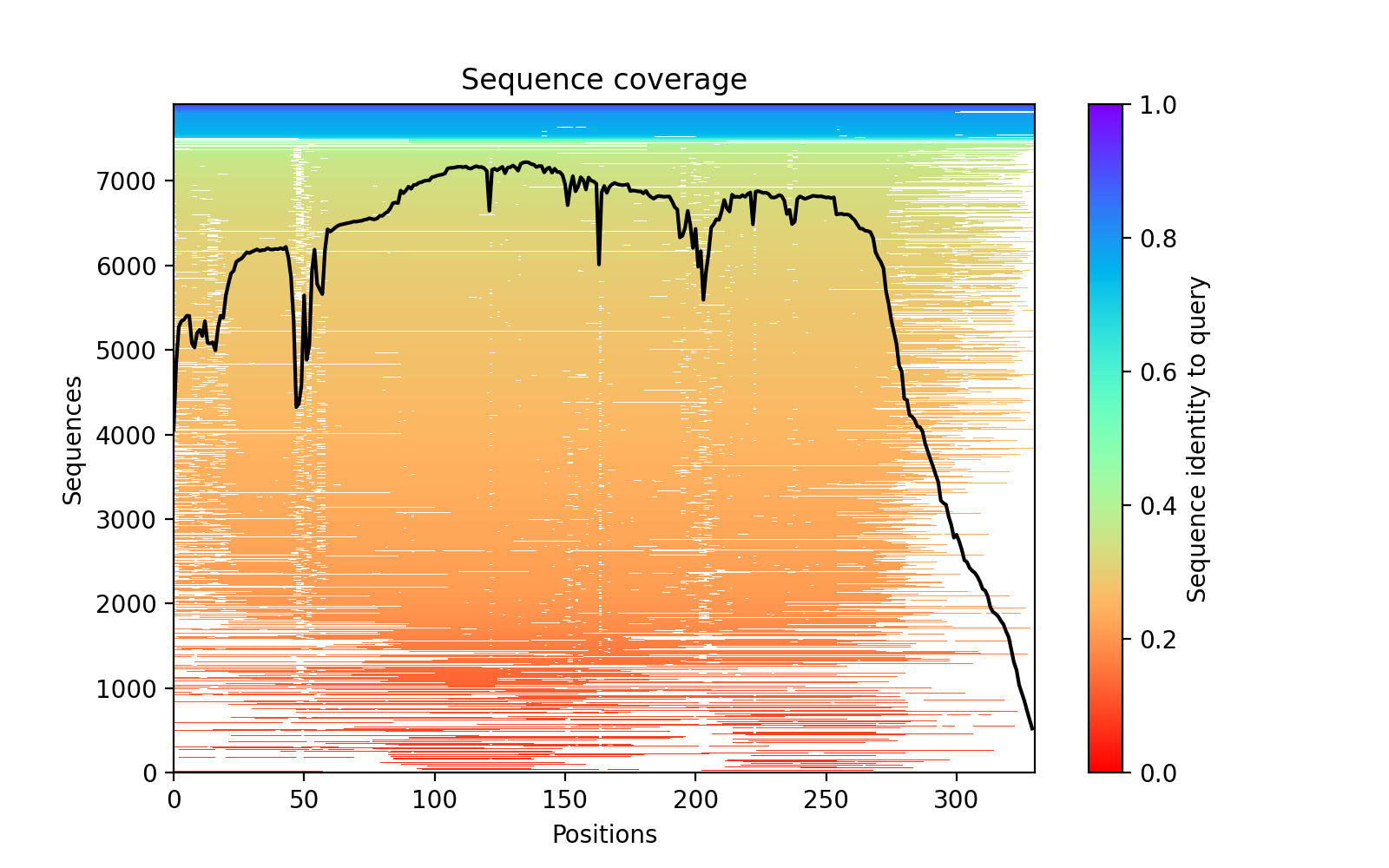

### af330_PAE.png

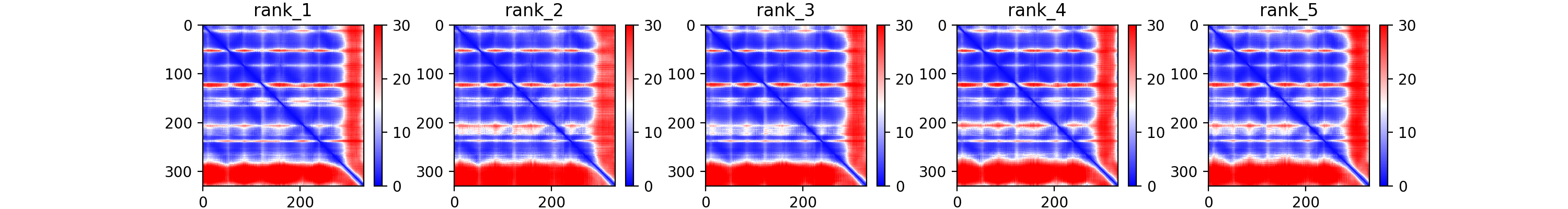

### af330_plddt.png

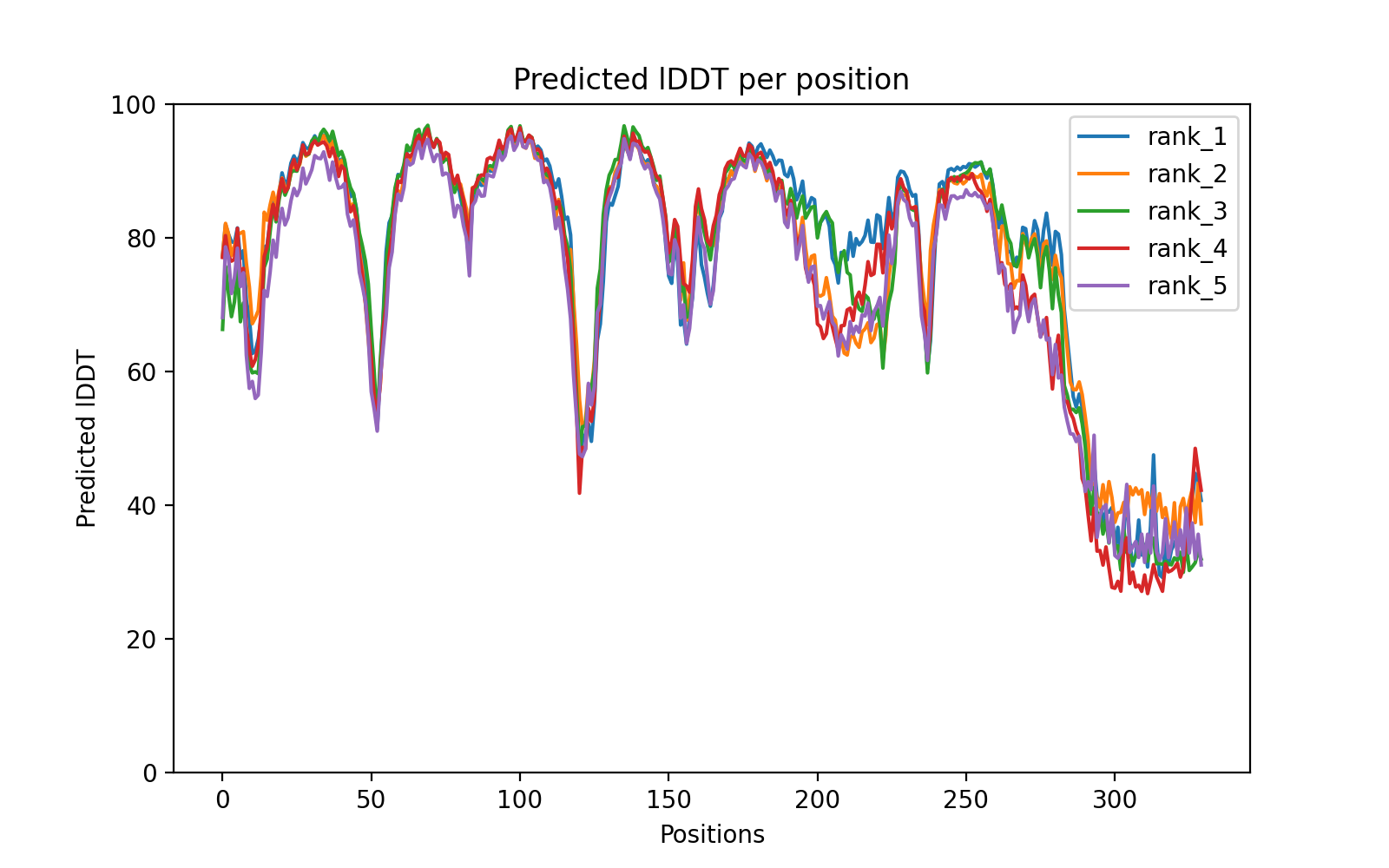

### af416_coverage.png

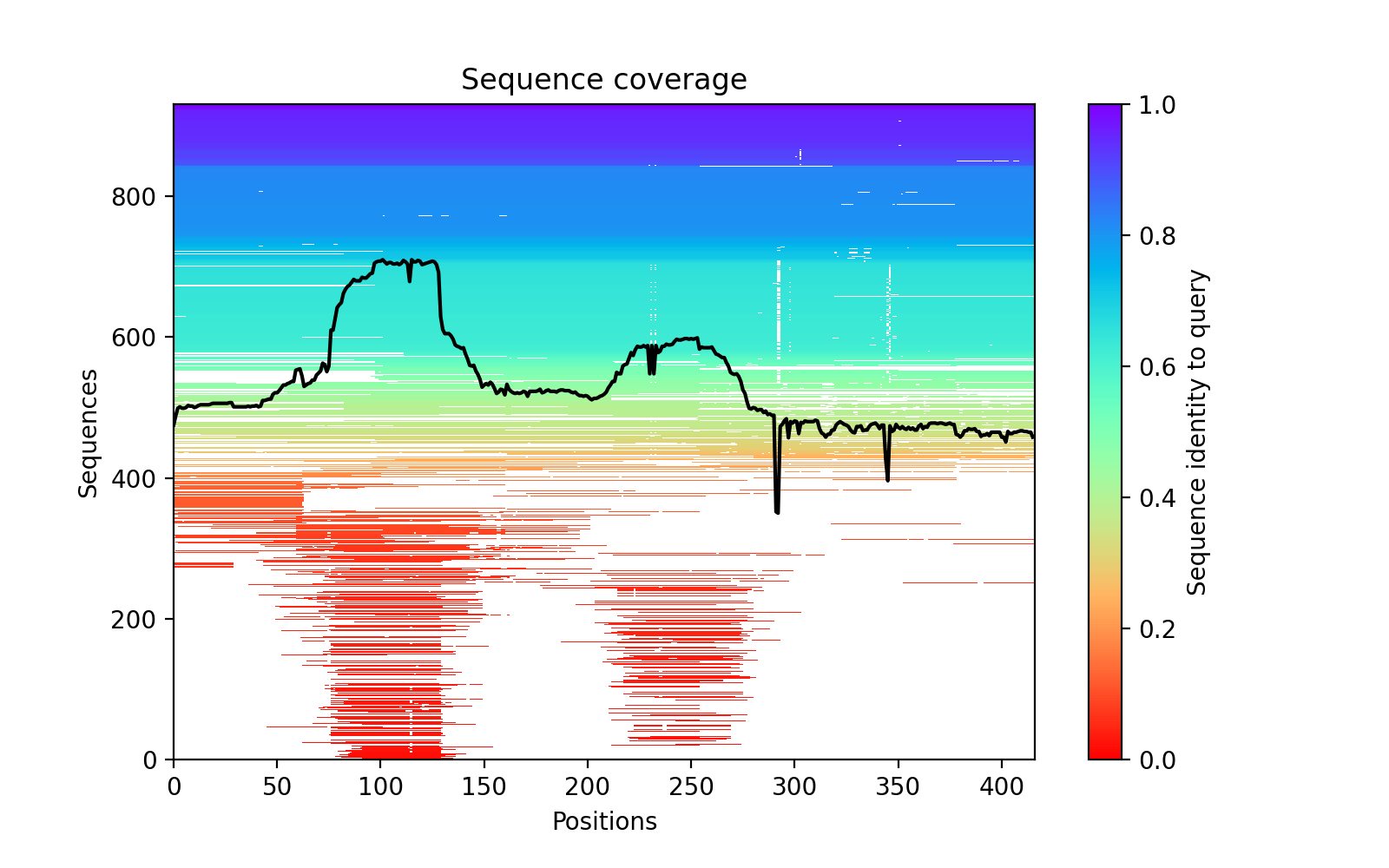

### af416_PAE.png

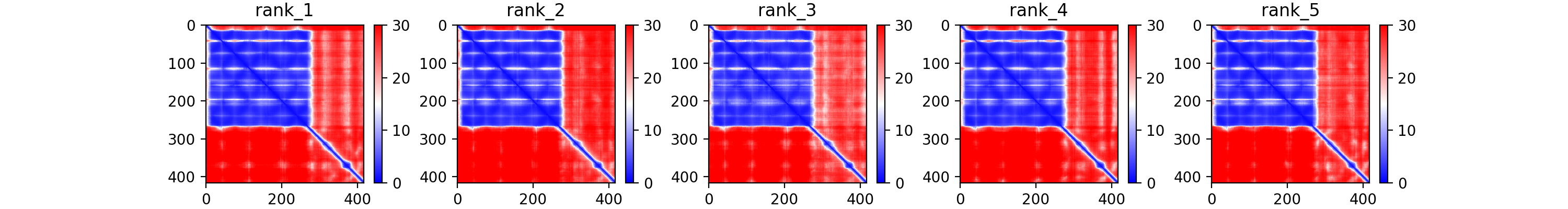

### af416_plddt.png

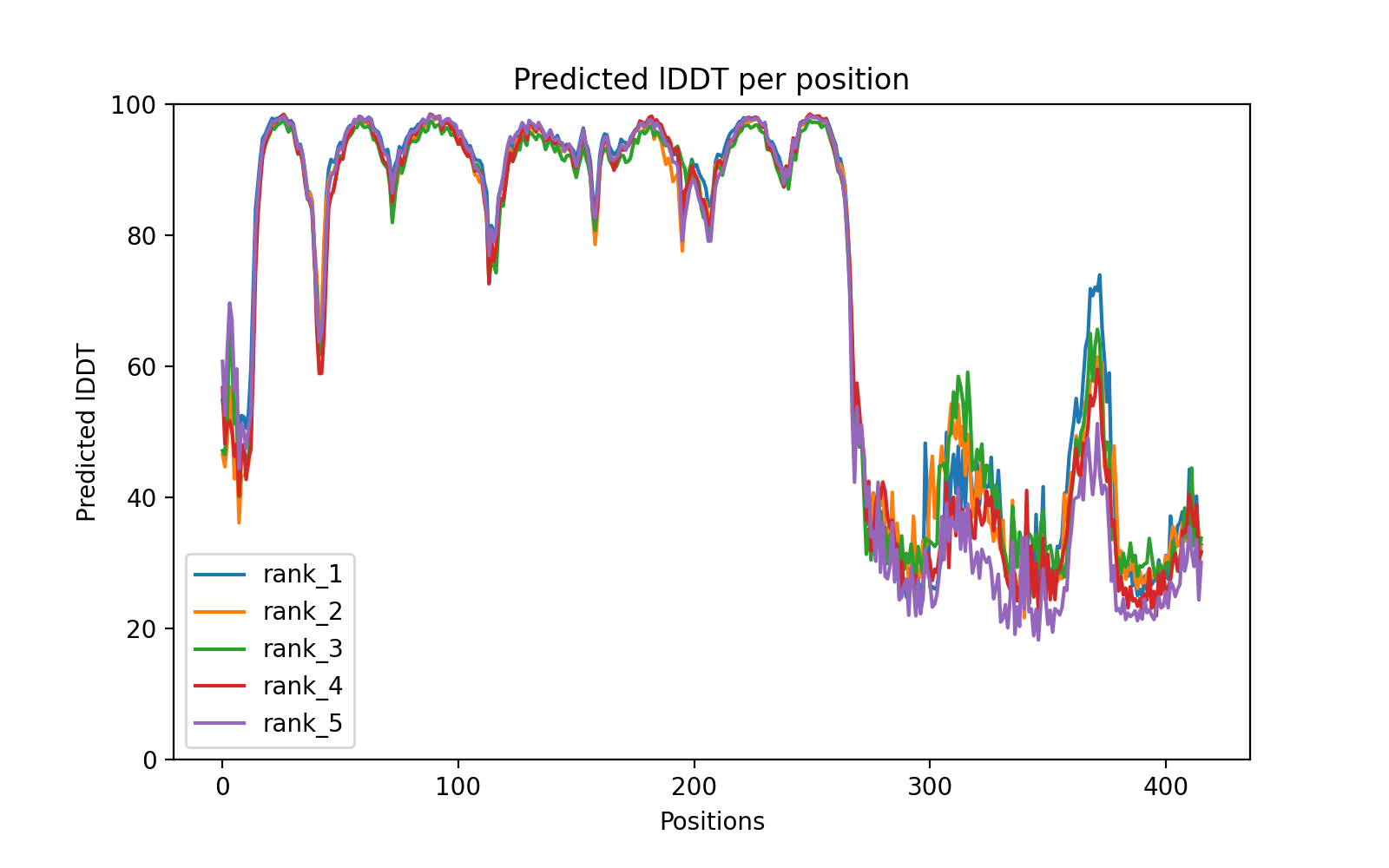

### af566_coverage.png

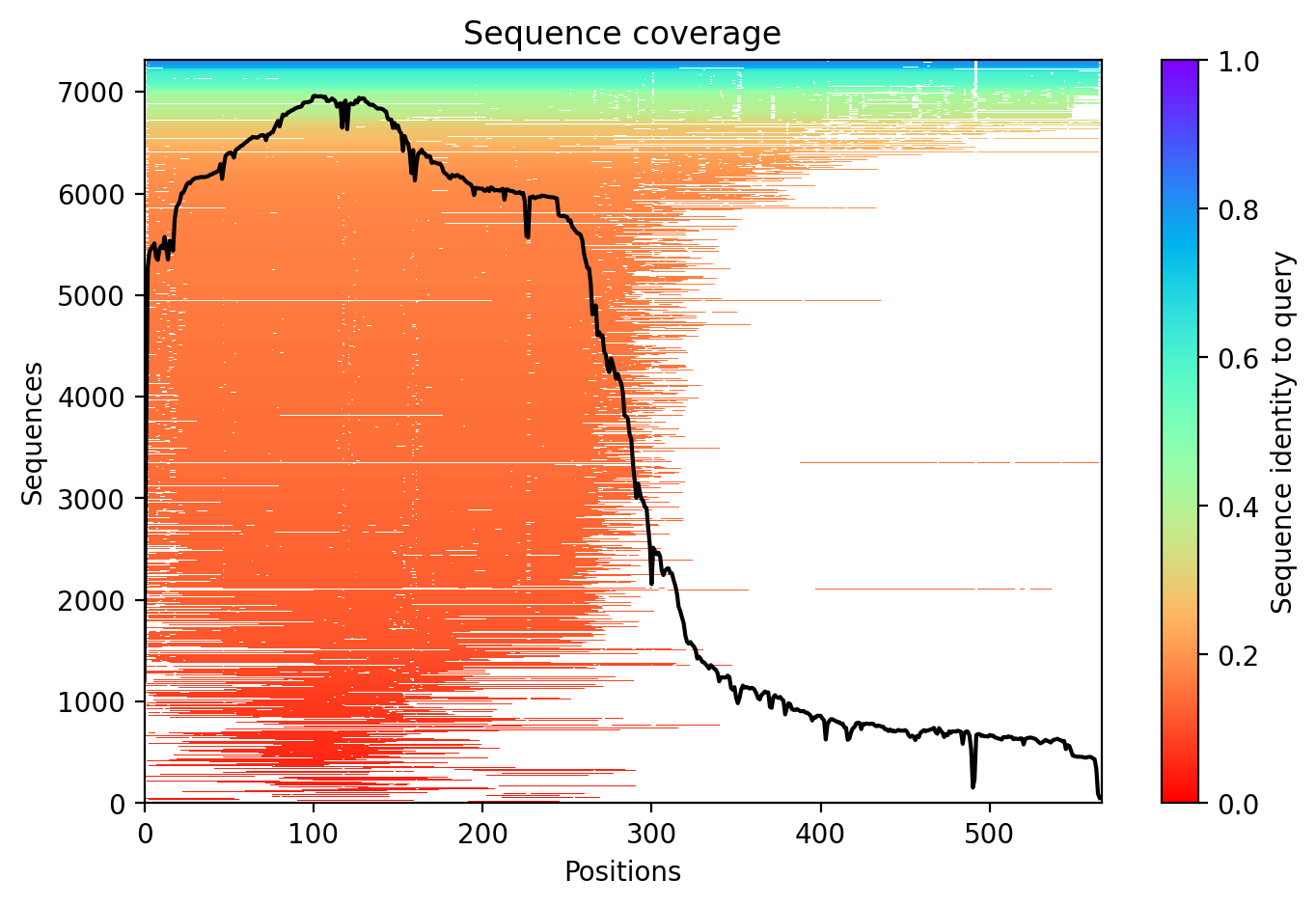

### af566_pae.png

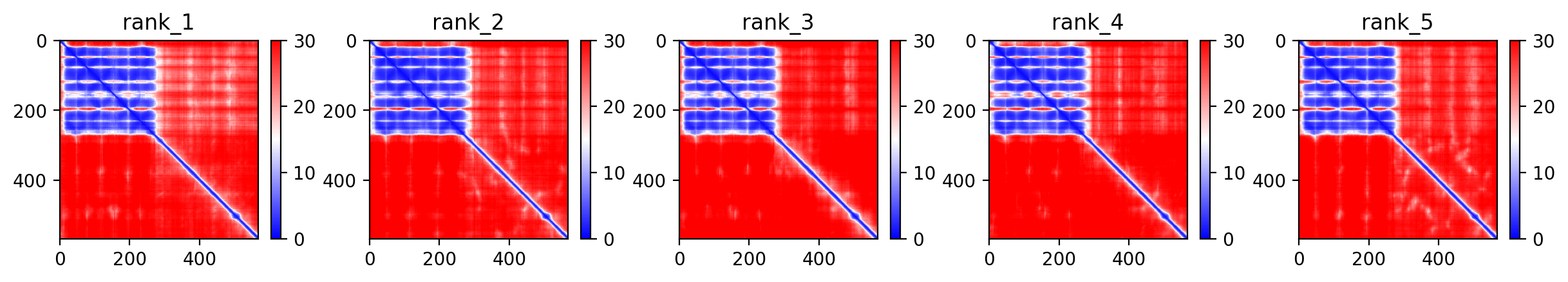

### af566_plddt.png

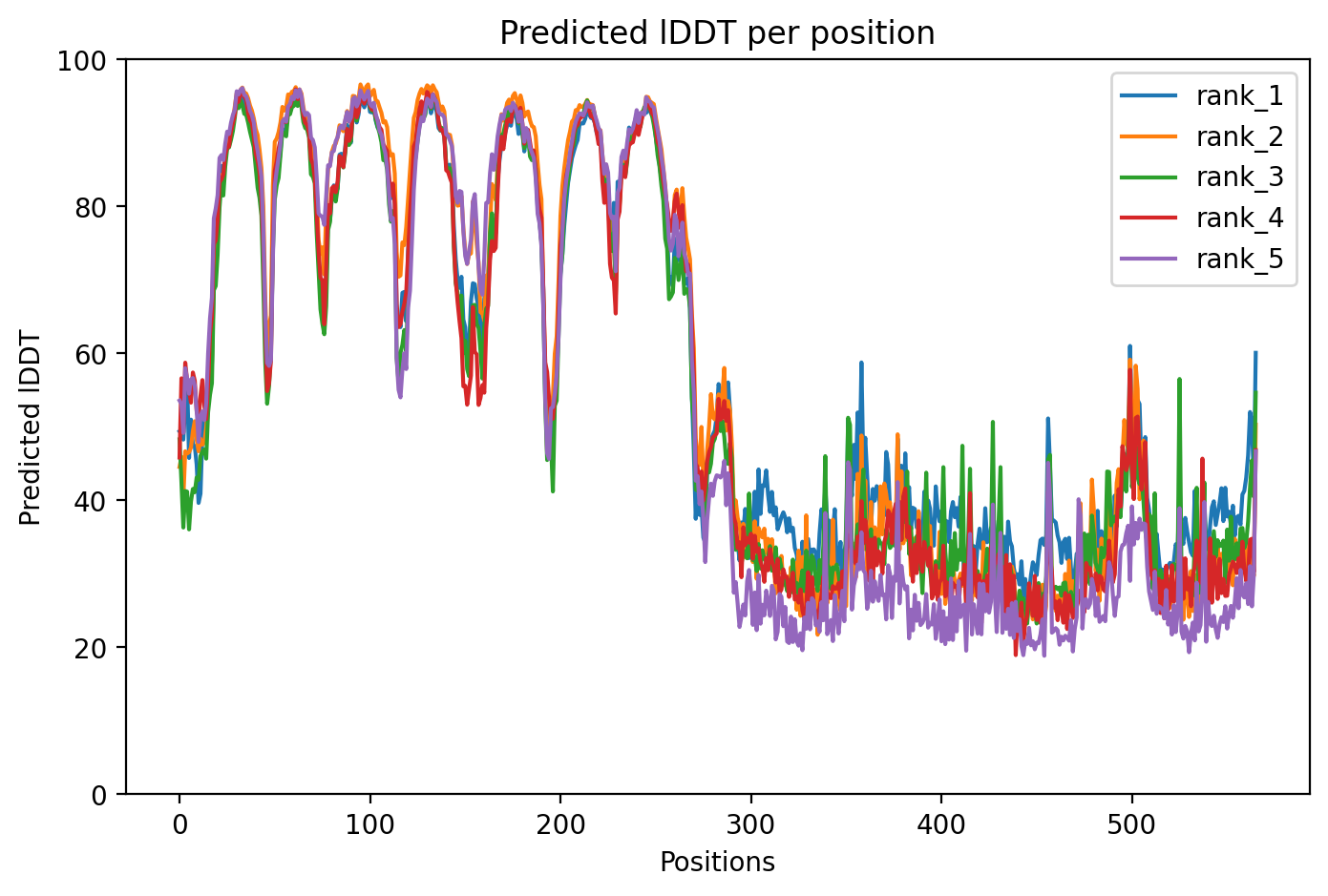

### af567_coverage.png

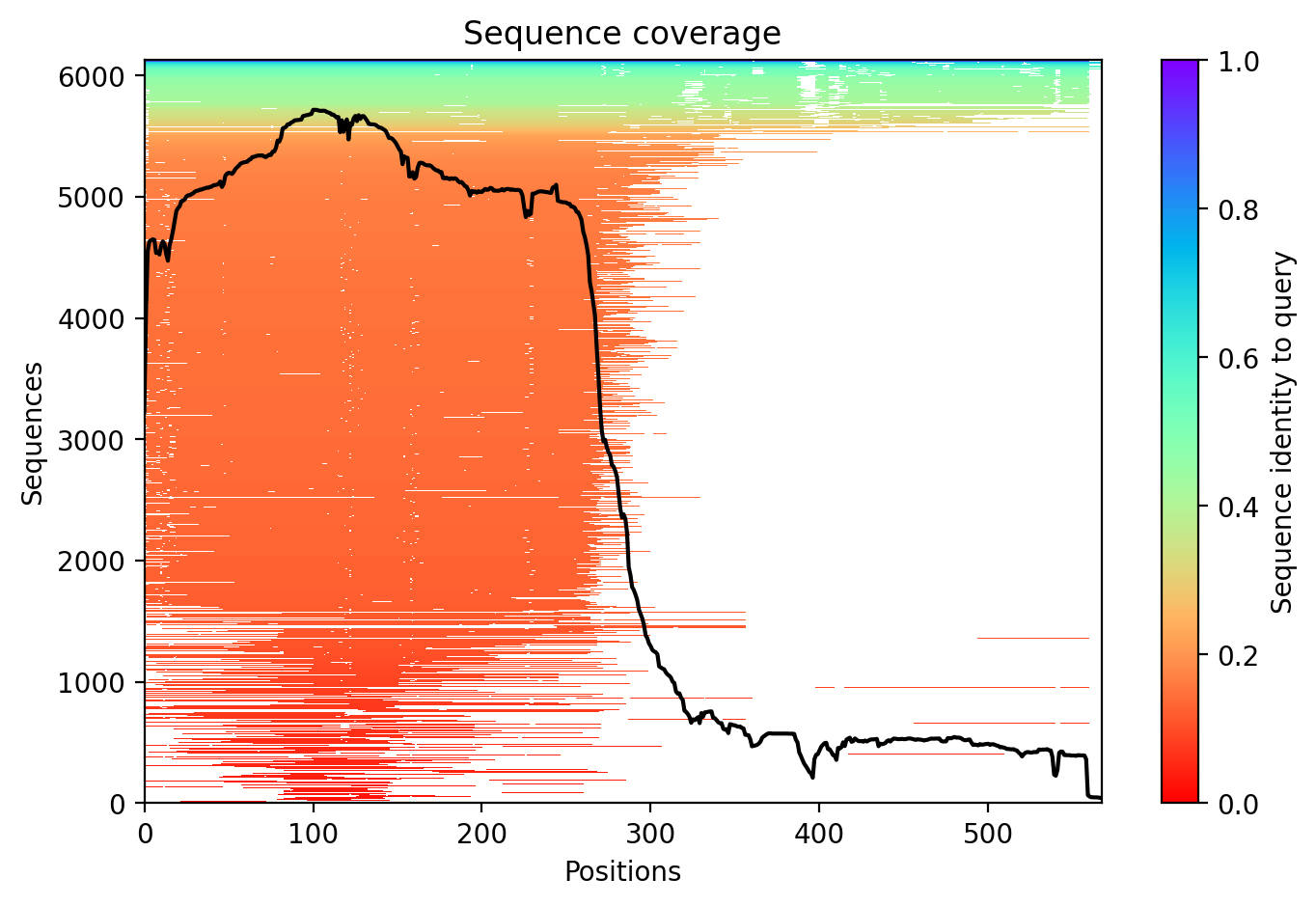

### af567_pae.png

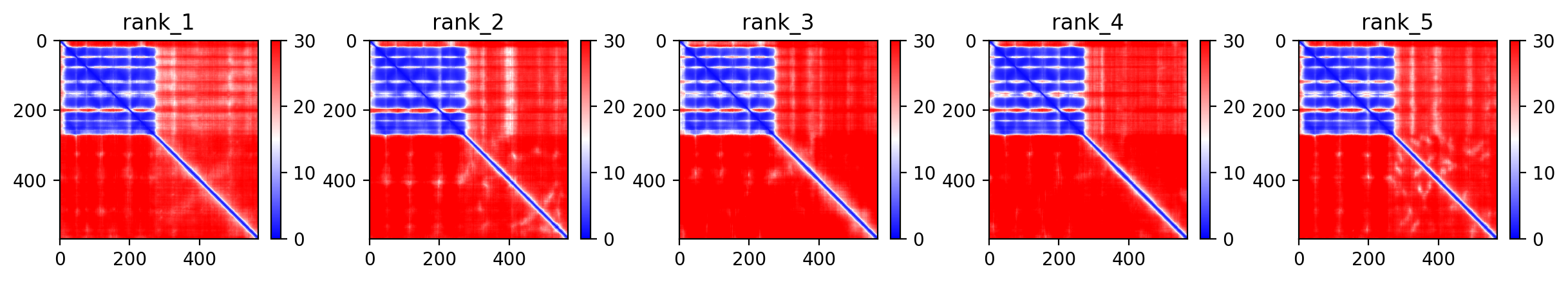

### af567_plddt.png

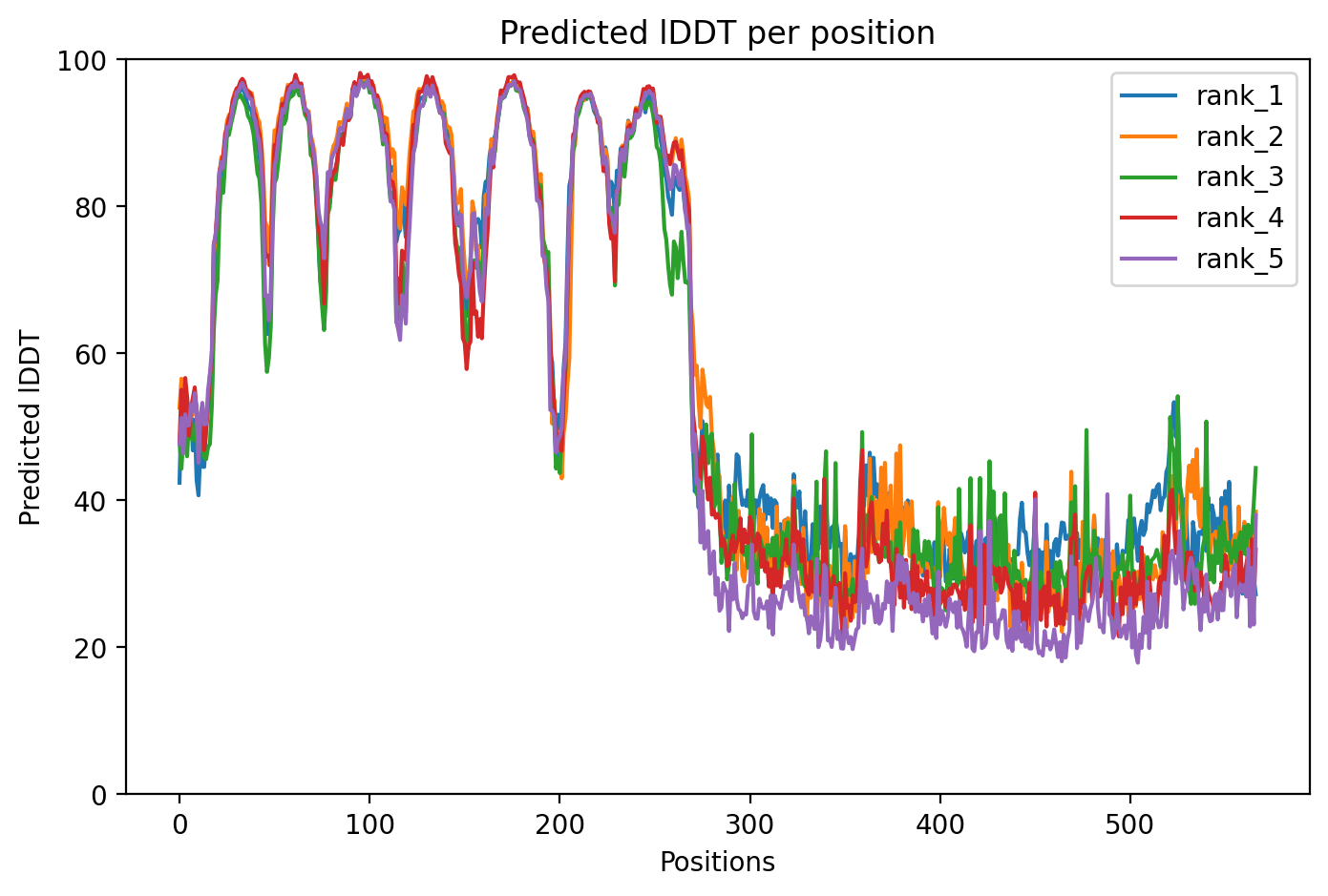

### af636_coverage.png

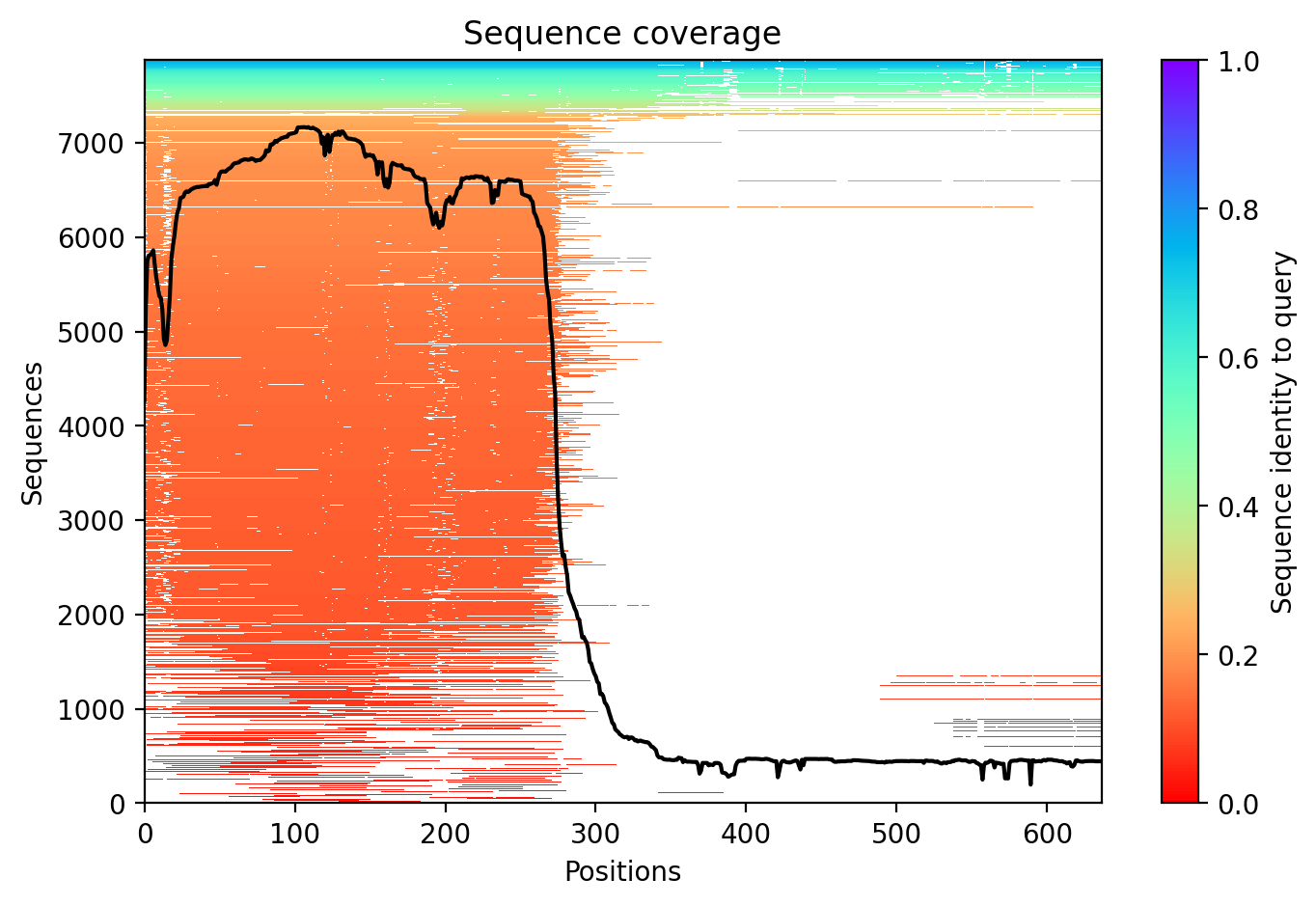

### af636_pae.png

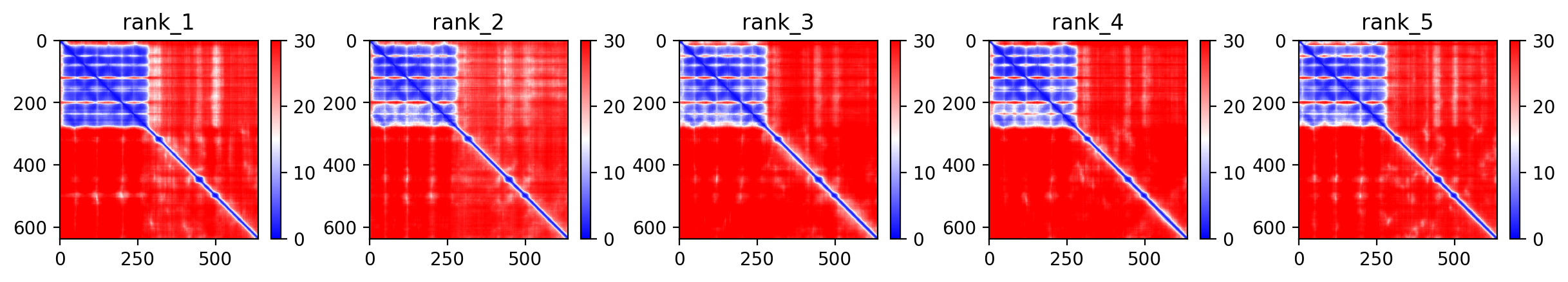

### af636_plddt.png

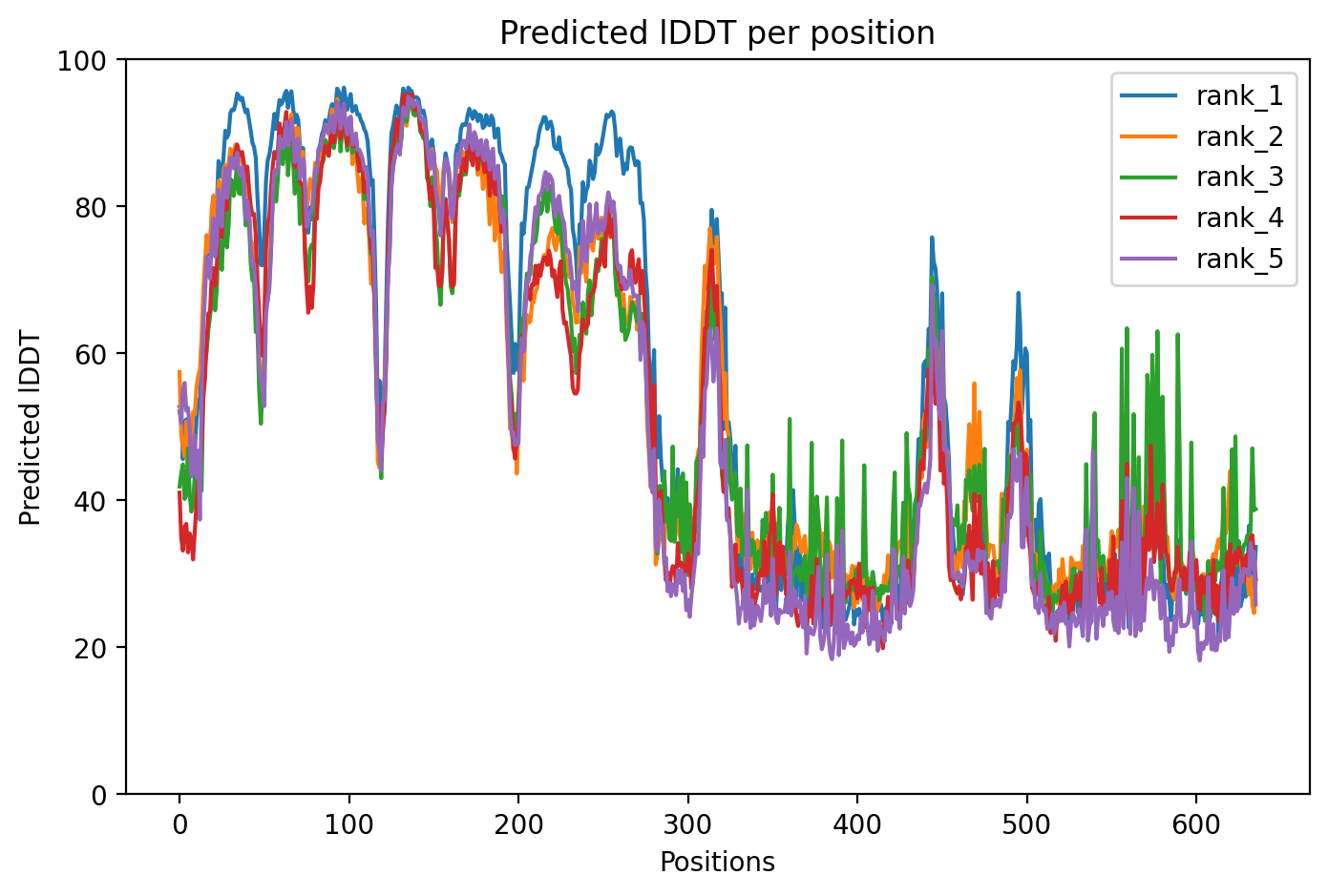

### af793_coverage.png

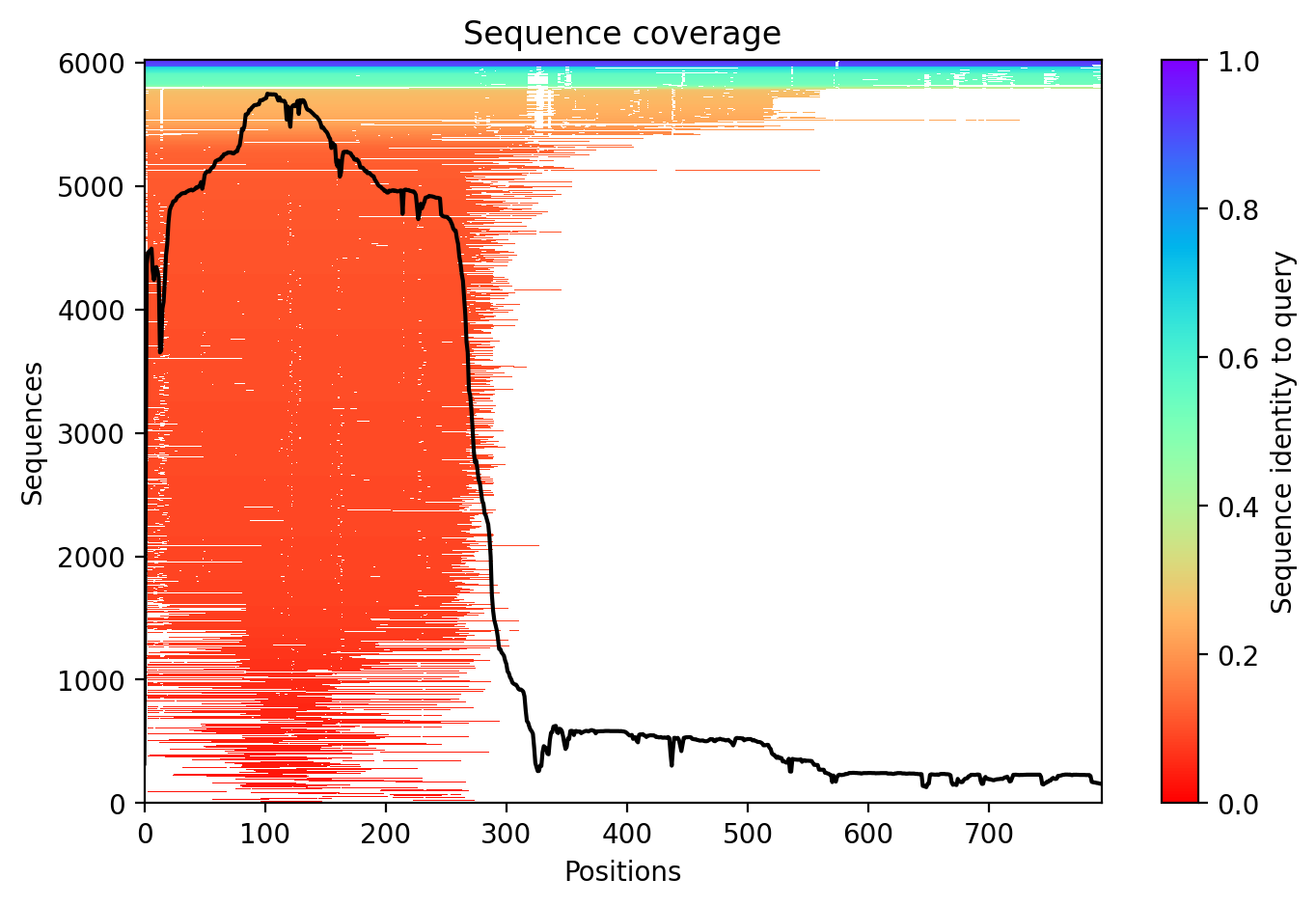

### af793_pae.png

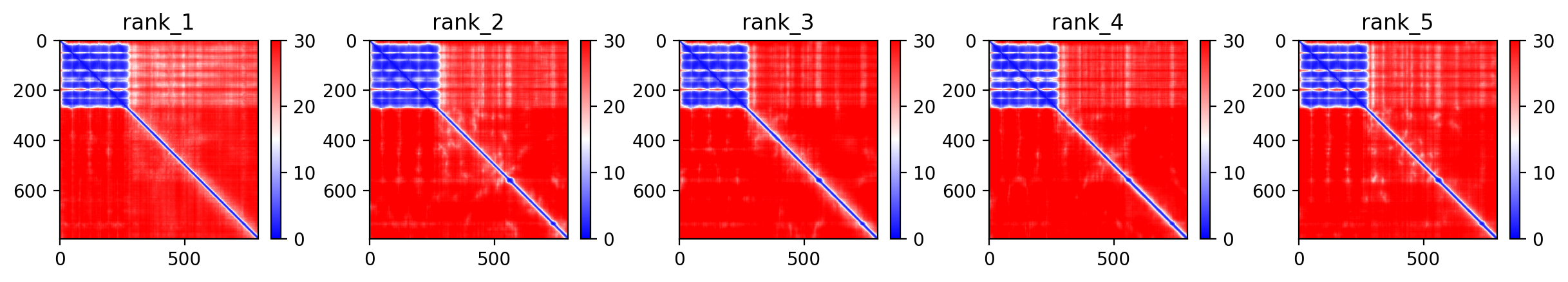

### af793_plddt.png

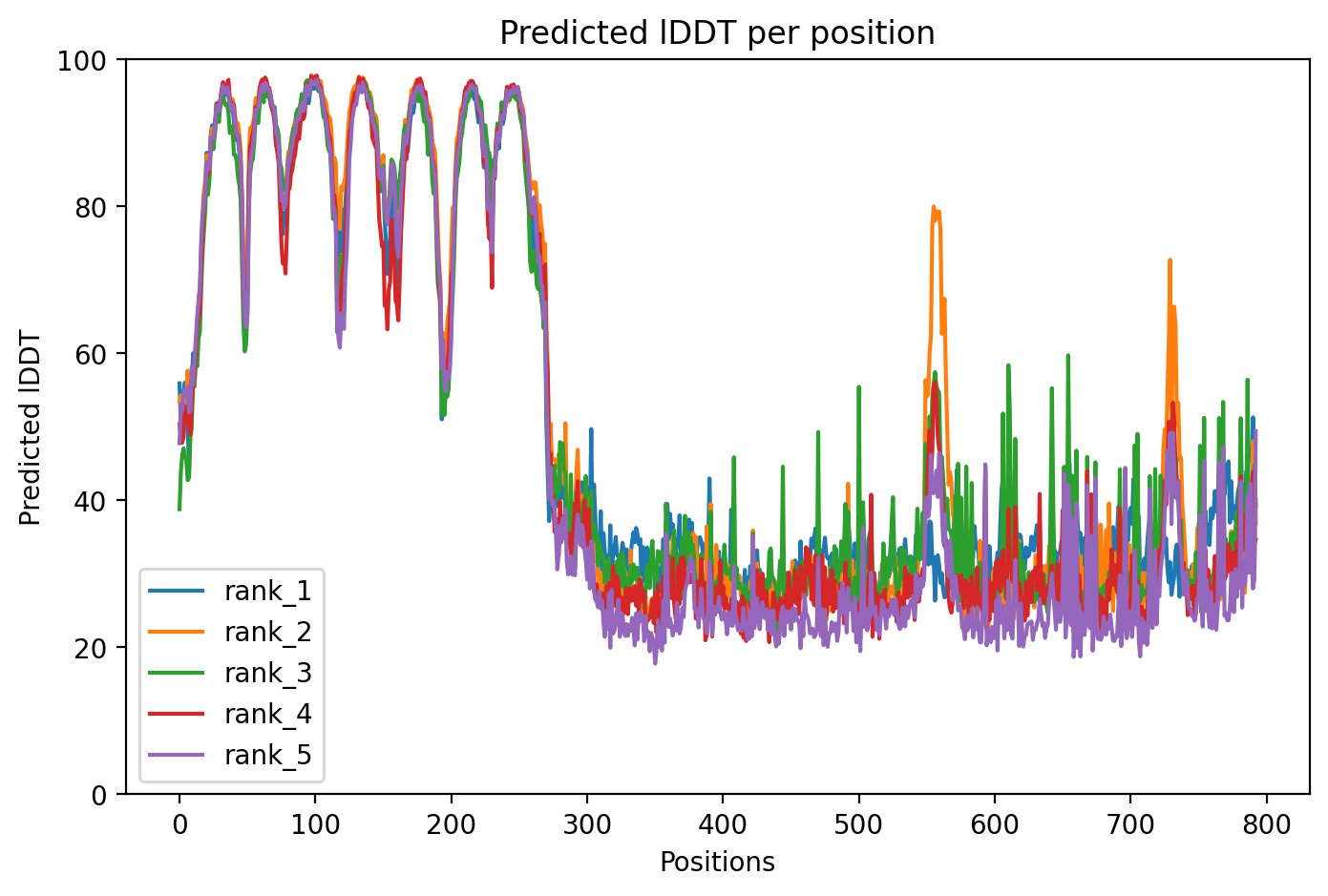
